# GABA facilitates spike propagation through branch points of sensory axons in the spinal cord

**DOI:** 10.1101/2021.01.20.427494

**Authors:** Krishnapriya Hari, Ana M. Lucas-Osma, Krista Metz, Shihao Lin, Noah Pardell, David A. Roszko, Sophie Black, Anna Minarik, Rahul Singla, Marilee J. Stephens, Robert A. Pearce, Karim Fouad, Kelvin E. Jones, Monica A. Gorassini, Keith K. Fenrich, Yaqing Li, David J. Bennett

## Abstract

Movement and posture depend on sensory feedback that is regulated by specialized GABAergic neurons (GAD2^+^) that form axo-axonic contacts onto myelinated proprioceptive sensory axons and are thought to be inhibitory. However, we report here that activating GAD2^+^ neurons, directly with optogenetics or indirectly by cutaneous stimulation, facilitates sensory feedback to motoneurons in awake rodents and humans. GABA_A_ receptors and GAD2^+^ innervation at or near nodes of Ranvier of sensory axons cause this facilitation, preventing spike propagation failure at the many axon branch points, which is otherwise common without GABA. In contrast, GABA_A_ receptors are generally lacking from axon terminals (unlike GABA_B_) and do not presynaptically inhibit transmitter release onto motoneurons. GABAergic innervation near nodes and branch points allows individual branches to function autonomously, with GAD2^+^ neurons regulating which branches conduct, adding a computational layer to the neuronal networks generating movement and likely generalizing to other CNS axons.

## Main

The ease with which animals move defies the complexity of the underlying neuronal circuits, which include corticospinal tracts (CSTs) that coordinate skilled movement, spinal interneurons that form central patterns generators (CPGs) for walking, and motoneurons that ultimately drive the muscles^1^. Sensory feedback ensures the final precision of such motor acts, with proprioceptive feedback to motoneurons producing a major part of the muscle activity in routine movement and posture^2, 3^, without which coordination is poor^4^. Proprioceptive sensory feedback is regulated by specialized GABAergic neurons (GAD2^+^; abbreviated GABA_axo_ neurons) that form axo-axonic connections onto the sensory axon terminals^5–7^. These neurons are thought to produce presynaptic inhibition of sensory feedback to motoneurons^8–10^ and possibly limit inappropriate sensory feedback^3, 6^. However, during movement the CST, CPG and even sensory neurons all augment GABA_axo_ neuron activity^10–15^ right at a time when sensory feedback is known to be increased to ensure precision and postural stability^2, 3^, raising the question of whether GABA_axo_ neurons have a yet undescribed excitatory action.

The long-standing view that GABAergic neurons and associated axonal GABA_A_ receptors produce presynaptic inhibition of proprioceptive sensory axon terminals in adult mammals actually lacks direct evidence. This is largely because of the difficulty in recording from these small terminals^15^ and the technical limitations of previously employed reduced spinal cord preparations (immature)^6, 16^ or anesthetized animals, since anesthetics themselves modulate GABA_A_ receptors^8, 17^. Thus, in this paper we used optogenetic approaches to directly target GABA_axo_ neurons in awake animals and in isolated whole adult spinal cord preparations. Surprisingly, we found that optogenetically activating these GABAergic neurons markedly facilitates sensory axon transmission to motoneurons via axonal GABA_A_ receptors, throwing into doubt the concept of presynaptic inhibition mediated by GABA_A_ receptors.

The mechanism by which GABA_A_ receptors are theorized to produced presynaptic inhibition is rather counterintuitive and based on indirect evidence^10, 17^. That is, sensory axons, like many other axons, have high intracellular chloride concentrations, leading to an outward chloride ion flow through activated GABA_A_ receptors^10, 18, 19^. Thus, GABA_A_ receptors cause a depolarization of sensory axons (primary afferent depolarization, PAD)^10, 15, 20–22^, which is on face value excitatory, rather than inhibitory, sometimes even evoking axon spikes^15^. Nevertheless, PAD and associated GABA_A_ receptors have variously been theorized to cause presynaptic inhibition by depolarization-dependent inactivation or shunting of sodium currents at the sensory axon terminals^21, 23^. However, we do not even know if terminals of large myelinated proprioceptive sensory axons express GABA_A_ receptors at all, despite their demonstrated innervation by GABA_axo_ neurons^5^. These terminals appear to lack the α5 subunit of extrasynaptic GABA_A_ receptors^15^ and the more ubiquitous β2/3 subunits of GABA_A_ receptors^24^, but this leaves open the possibility that they express other GABA_A_ subunits or GABA_B_ receptors. We thus examined this question here and found again that GABA_A_ receptors are generally not at these axon terminals, but are instead near sodium channels (Na_V_) of the nodes of Ranvier throughout the myelinated regions of the axon, together with innervation by GABA_axo_ neurons, consistent with earlier electron microscopy observations of GABAergic innervation of afferent nodes^25^ and imaging of α5 subunits^15^. What then is the function of such GABA_A_ receptors near sodium channels?

An unexplored possibility is that the depolarizing action of GABAA receptors (and GABA_axo_ neurons) near nodes aids sodium spike propagation between axon nodes. This has not previously been considered, as spikes are thought to securely propagate from node to node, at least in the orthodromic direction^15^. Myelinated proprioceptive axons branch extensively in the spinal cord^15^(Fig. 1a) and each branch point poses a theoretical risk for spike propagation failure at downstream nodes^26, 27^. However, branch points are always located at nodes (Na_V_)^15^, likely to minimize this failure. Nevertheless, indirect evidence has suggested that propagation failure can occur^28–31^. Thus, in the present study we sought direct evidence of nodal spike failure near branch points and examined whether GABA_A_ receptors near or at nodes facilitates afferent conduction by preventing this failure. We already know that PAD and GABA_A_ receptors lower the threshold for initiating axon spikes by extracellular stimulation^32^, and even initiate spikes^15^, but do not know whether they aid normal spike propagation. We found that spike propagation depends so heavily on GABA that blocking GABA action makes the majority of proprioceptive sensory axons fail to propagate spikes to motoneurons, and thus GABA near sodium channels provides a powerful mechanism to turn on specific nodes and branches to regulate sensory feedback.

**Fig. 1.**
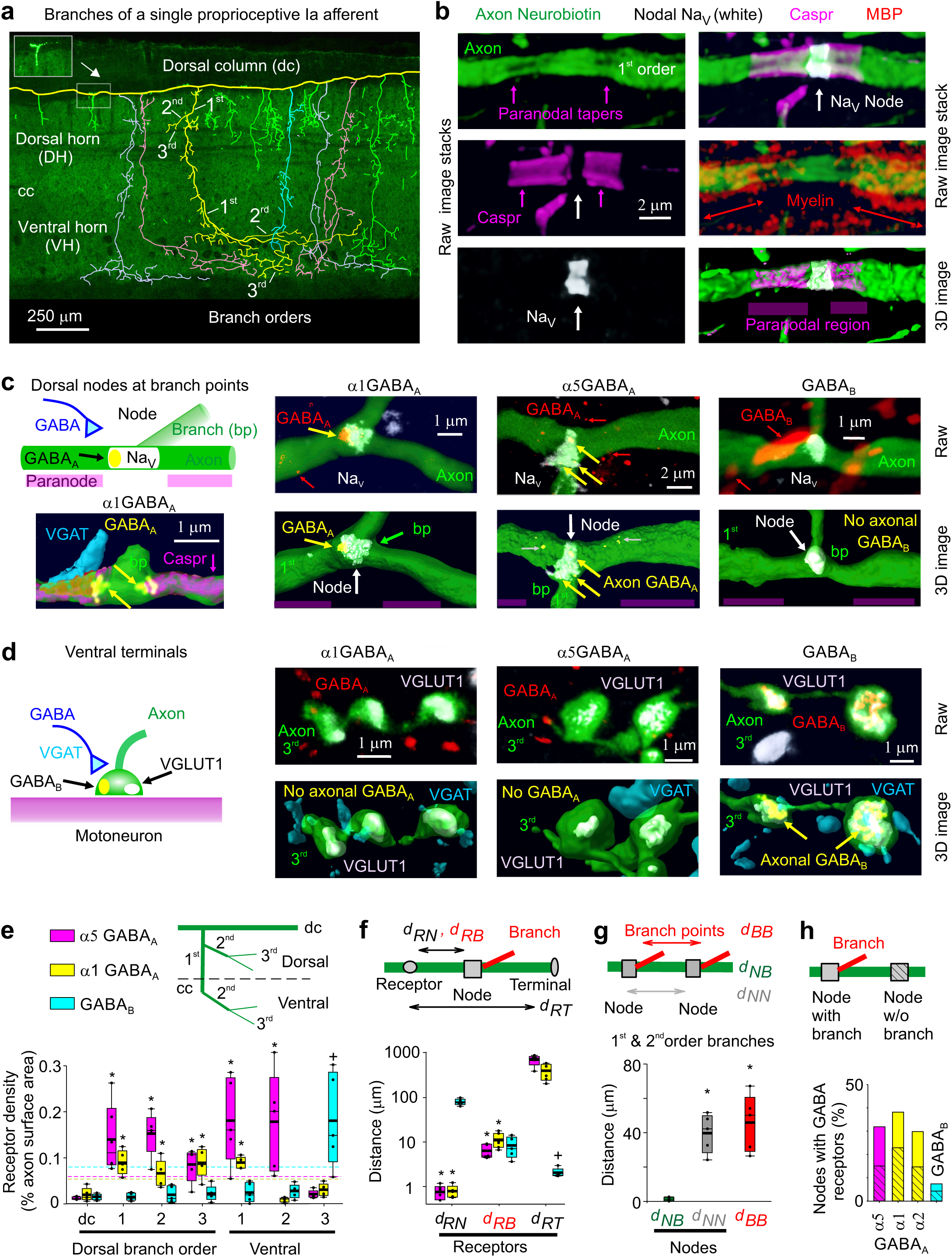
Nodal GABA_A_ and terminal GABA_B_ receptors in rats. **a,** Neurobiotin filled proprioceptive Ia axon in the sacrocaudal rat spinal cord (S4 and Ca1), reconstructed from fluorescent images (inset), with branching order indicated and different primary branches distinguished by color. Some ventral branches truncated for clarity (green). Axon diameter not to scale. Central canal: cc. Dorsal columns: dc. Dorsal and ventral horns: DH and VH. **b**, Node on axon branch in DH immunolabelled for sodium channels (Na_V_), paranodal Caspr and myelin (MBP), shown with raw images (maximum projection of z-stack), with the paranodal taper indicated, and co-labelling within the axon rendered in 3D (bottom). 1^st^ order branch in DH. **c-d**, α1 GABA_A_, α5 GABA_A_ and GABA_B_ receptor immunolabelling in axon branches (raw maximum projection: top row, 3D reconstruction: bottom), with all receptors colocalized with the axon labelled yellow. Receptor clusters specifically in the axon membrane indicated with yellow arrows. Some α5 GABA_A_ receptors are in axon cytoplasm (yellow with gray arrow) or in nearby neurons (red), and not in axon membrane. In (**c**) nodes identified by Na_V_ (or Caspr) and paranodal taper, and located at branch points (bp, 1^st^ to 2^nd^ bp in DH). Raw images for bottom left 3D image of (**c**) detailed in Extended Data Fig 1f. In (**d**) ventral terminal boutons identified by vesicular glutamate transporter 1 (VGLUT1) and adjacent to motoneurons. GABAergic contacts identified by vesicular inhibitory amino acid transporter (VGAT). **e**, Receptor densities on axon branches of varying order in dorsal (dorsal and intermediate laminae) and ventral regions. Box plots: median, thin line; mean, thick line; interquartile range, box; error bars detailed in Methods. Dashed lines: lower confidence interval, 1 SD below mean maximum density. *significantly more than ventral terminal (3^rd^ order) receptor density, + ventral terminal receptor density significantly more than 1^st^ and 2^nd^ order branch densities, *P <* 0.05, *n* = 5 rats each, with 11, 17 and 12 independently filled and reconstructed axons for α1 GABA_A_, α5 GABA_A_ and GABA_B_ receptors, respectively. **f**, Distances from GABA receptor clusters in the membrane to nodes (d_RN_, Na_V_), branch points (d_RB_) or ventral terminals at motoneurons (d_RT_). Distances to 1^st^ and 2^nd^ order dorsal and ventral nodes similar and pooled, as were branch points. *significantly less than d_RT_. + significantly less than d_RN_ and d_RB_; *n* = 5 rats, with 89, 36 and 70 clusters for α1 GABA_A_, α5 GABA_A_ and GABA_B_, respectively. **g**, Distances between branch points (d_BB_), nodes (Na_V_ clusters, d_NN_), and branch points and their nearest node (d_NB_) for 1^st^ and 2^nd^ order branches. On dorsal columns d_NN_ = 243 ± 117 µm (not shown). *significantly larger than d_NB_; *n* = 5 rats, same axons as (**e**), with 95 nodes and 57 bp. **h**, Proportion of nodes with GABA receptors, with and without (hashed) nearby branch points; n = 5 rats each, with 86, 75, 91, and 103 nodes for α5, α1, α2 GABA_A_ and GABA_B_ receptors, respectively.

## Results

### Nodal GABA_A_ and terminal GABA_B_ receptors

To confirm and extend previous observations that GABA_A_ receptors are near nodes of proprioceptive sensory axons (group Ia) rather than at ventral terminals^15, 24^, we immunolabelled the most common subunits of synaptic and extrasynaptic GABA_A_ receptors expressed in these axons^33^, both in rats with neurobiotin filled axons (Fig. 1) and VGLUT1^Cre/+^ mice with axons labelled by a reporter gene (Extended Data Fig. 1). GABA_A_ receptors containing α5, α1, α2 and γ2 subunits were expressed on these axons, especially near sodium channels (< 6 µm away; Fig. 1c-f, Extended Data Fig. 1). Specifically, GABA_A_ receptors were in the plasma membrane on large myelinated 1^st^ and 2^nd^ order branches in the spinal cord at their nodes (identified by large sodium channel clusters, nearby paranodal Caspr, and/or axonal tapers; Fig. 1a-c,e; Extended Data Fig. 1a,c,f), and on short unmyelinated terminal branches in the dorsal and intermediate laminae (3^rd^ order; Figs. 1a,e). The latter were near the nodes on 1^st^ order branches (< 100 µm away) where they can influence these nodes, consistent with previous observations of axonal GABAergic contacts^25, 34^. In contrast, GABA_A_ receptors were mostly absent from the long unmyelinated ventral terminal branches, where the axon boutons synapse onto motoneurons in the ventral horn (3^rd^ order; Figs. 1a,d,e; Extended Data Fig. 1b,d,g,h), which also generally lacked sodium channels^15^. This left GABA_A_ receptors on average far from the terminal boutons contacting motoneurons (∼500 µm; Fig. 1f) relative to the axon space constant (λ_S_ ∼90 µm), with the majority of receptors in dorsal and intermediate laminae. Nodes were widely spaced, as were branch points (∼50 µm separation, Fig. 1g), but branch points were always near nodes (Na_V_; Fig. 1c,g), the latter providing additional evidence for nodal GABA_A_ receptors, since these receptors were near branch points (Fig 1f). Nodes sometimes occurred without branching (49%, as in Fig. 1b, rat). Overall, synaptic α1 and α2 and extrasynaptic α5 GABA_A_ receptors were each expressed in about 30% of nodes, roughly equally distributed among branched and unbranched nodes (Fig 1h). Importantly, the majority of nodes were electrotonically close (< 90 µm, λ_S_) to GABA_A_ receptors on a neighboring node (96%) or bouton (80%) on the same axon, thus leaving little doubt that nodes are somehow influenced by GABA_A_ receptors. While synaptic α1 and α2 GABA_A_ receptors were in single large clusters, extrasynaptic α5 GABA_A_ receptors were in multiple smaller clusters, some in the membrane at the node (yellow arrows), and others in the cytoplasm near the node (grey arrows, Fig 1c and Extended Data Fig 1a), though all membrane receptors were at nodes. In contrast to GABA_A_ receptors, GABA_B_ receptors were found mostly on terminal branches in the ventral horn where boutons had dense receptor expression (Fig. 1d-f, Extended Data Fig. 1e,g,h), and not usually on larger myelinated ventral or dorsal branches, and thus absent from nodes (Figs. 1c,e,h; Extended Data Fig. 1e).

### Propagation failure in dorsal horn axon branches

Considering that GABA_A_ receptors are expressed in large myelinated dorsal branches of proprioceptive axons, we next directly recorded from these branches in the dorsal horn of rat and mouse spinal cords (Figs. 2 and 3) to examine whether spike propagation depends on these receptors. When we stimulated the dorsal root (DR) containing the axon branch, an all-or-nothing spike was recorded in many branches (Figs. 2b, 3d) at the latency of the fastest afferent volley that arrived at the spinal cord (group Ia afferents; EC in Fig. 2b). However, in other axon branches this spike did not occur (∼20%), but at the same latency there was a small all-or-nothing residual spike (*failure potential, FP;* Ia afferents). This FP was indicative of a spike activating a distant node, but failing to propagate further to the recording site, leaving only its passively attenuated potential, with smaller FPs reflecting more distal failure points in the spinal cord (Figs. 2c-g, 3e-f; typically a few nodes away). Failure never occurred in the DR itself (Fig. 2f). The failing branches with FPs were otherwise indistinguishable from non-failing axon branches, exhibiting full spikes (> 60 mV) with current injection pulses (directly evoking spike; or aiding DR spike, Fig. 2c*ii*, g), and low conductances and resting potentials (∼ −65 mV, Fig. 2h), ruling out penetration injury. With high repetitive DR stimulation rates all branches (100%) exhibited propagation failure and an associated FP (Fig. 2e-g), again with the FP implying that the spike is reliably initiated in the DR, but incompletely propagates within the spinal cord.

**Fig. 2.**
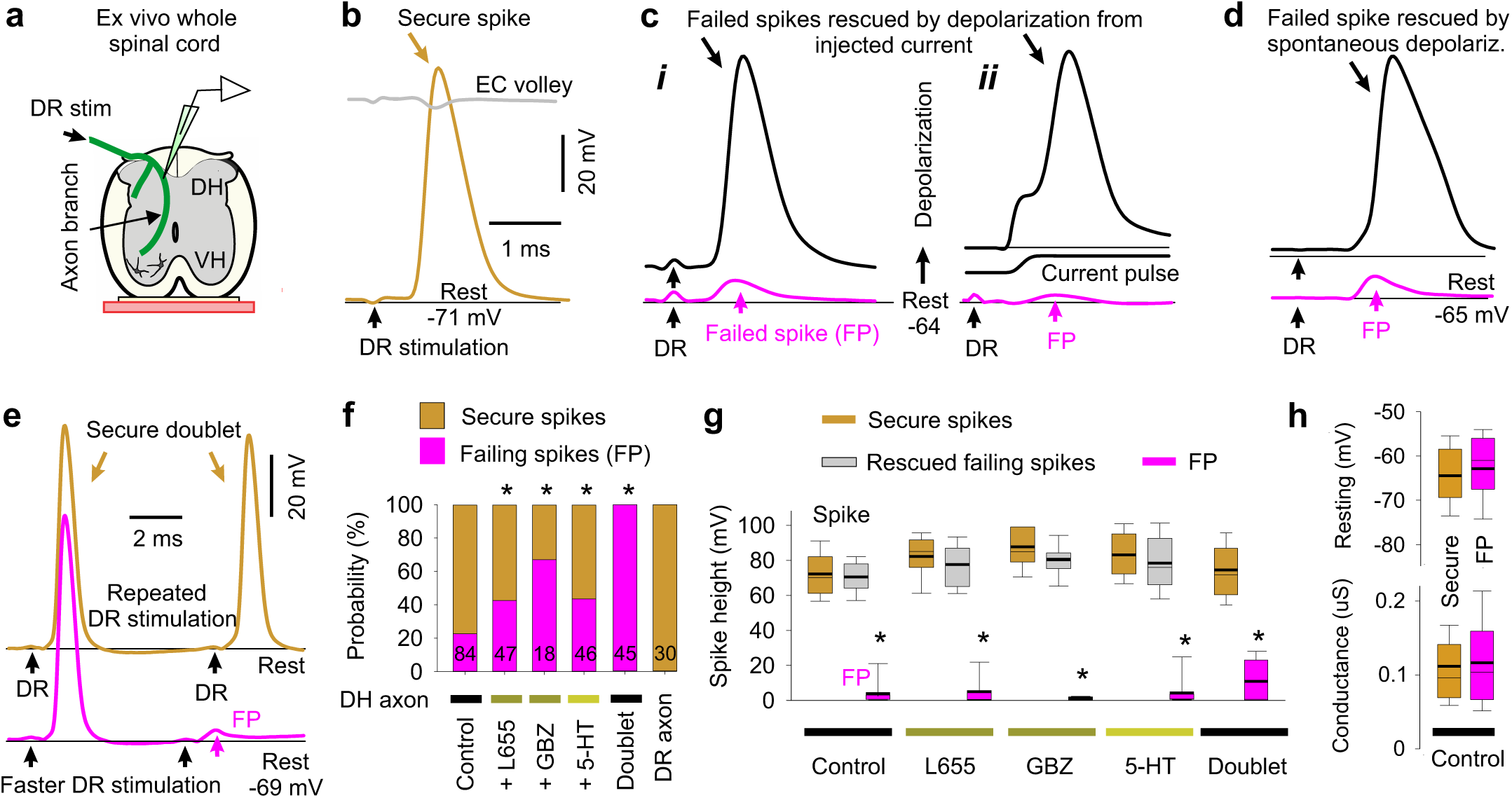
Spike failure. **a,** Recording from ex vivo whole adult rat spinal cord. **b-d**, Intracellular recordings from proprioceptive axon branches in the dorsal horn (DH), with dorsal root (DR) stimulation (1.1x T, 0.1 ms; T: afferent volley spike threshold) evoking a spike in some branches (secure, **b**) and only a failed spike in others (failure potential, FPs; **c**, **d**), but depolarization restoring full spikes (black, **c** and **d**). Averages of 10 trials at 3 s intervals. Resting potential: thin line. EC: extracellular afferent volley. Axons from S4 sacral DR. **e**, Fast repeated DR stimulation induced failure in secure spikes (not failing on first stimulation, 1.1xT). Threshold interval (longest) for failure (FP, pink), and just prior to threshold (gold). **f**, Proportions of DH axon branches (or DR axons) failing to spike with DR stimulation under control resting conditions and with L655708 (0.3 µM), gabazine (50 µM; GBZ), 5-HT (10 µM) or fast repetition (doublet, **e**). *significantly more than control, χ-squared test, number of axons indicated in bars (*n* = 84, 47, 18, 46, 45 and 30, from 11 rats each condition). **g**, Summary of spike and FP heights in secure and failing branches. Box plots. *significantly smaller than secure spike, *P <* 0.05, for spikes in (**f)**. **h**, Resting membrane potential and conductance for secure and failing branches, not significantly different, *P* > 0.05, for control axons in (**f**).

**Fig. 3.**
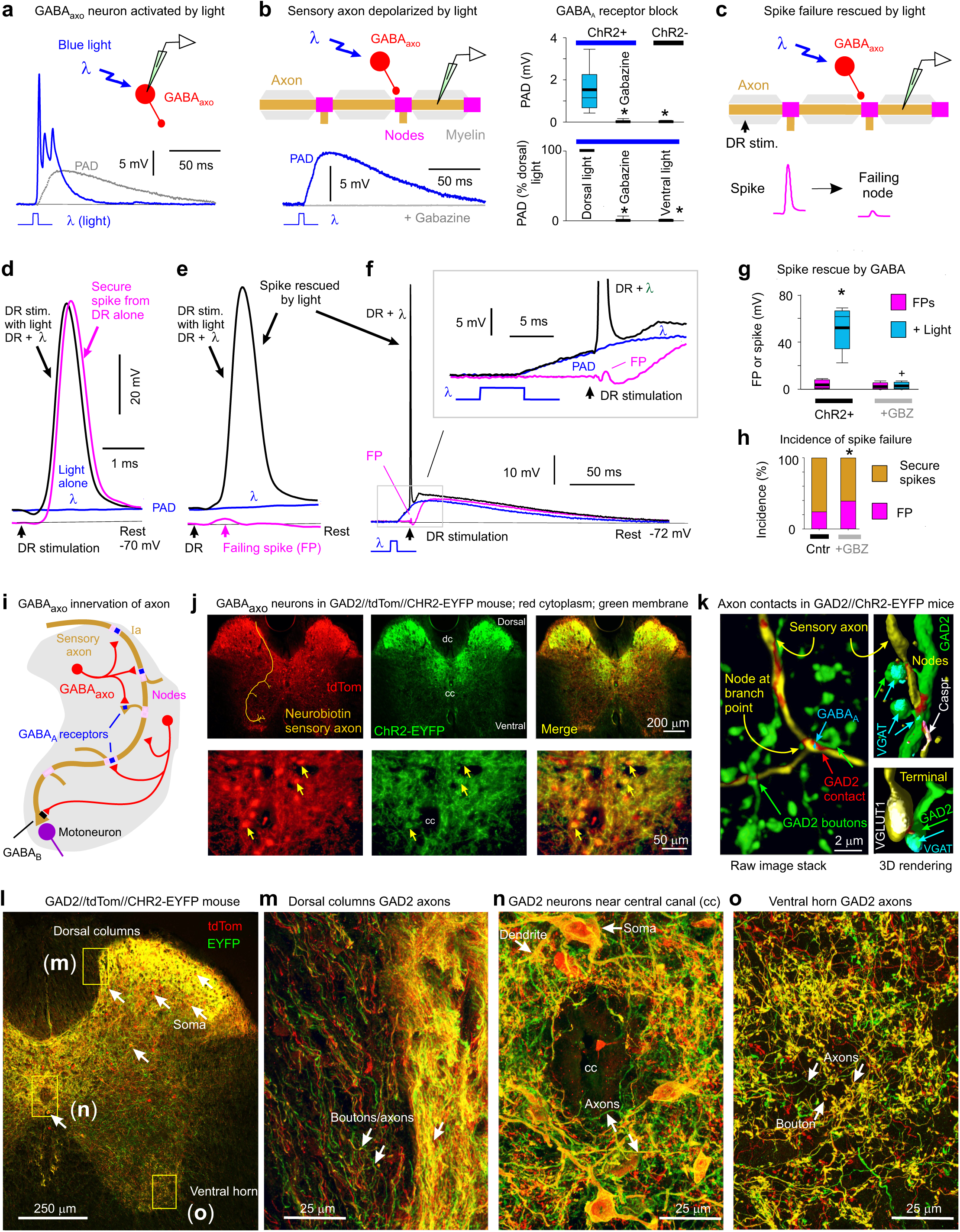
Nodal facilitation by GABA_axo_ neurons. **a,** Intracellular recording from GABA_axo_ neuron in ex vivo spinal cord of GAD2//ChR2-EYFP mouse, with ChR2 activated with a light pulse focused on the dorsal horn (5 ms, λ = 447nm laser, 0.7 mW/mm^2^, 1.5x light threshold to evoke PAD, T) causing a long depolarization and asynchronous spiking (cell isolated in 50 µM gabazine). Average of 10 trials at 0.3 Hz, blue. Cell resting at −61mV. PAD from (b) also shown, grey. b, Intracellular recording from proprioceptive axon branch (in DH, sacral S3, resting at −71 mV; average of 10 trials at 0.3 Hz) with same dorsal light pulse (1.5xT) producing a long depolarization (PAD). Box plots summary. *significantly less with gabazine or omitting ChR2 (control mice) or focusing light on the ventral rather than dorsal horn (ventral light), *n =* 14 axons, from 5 mice each. c-g, DR stimulation at rest (1.1xT) evoked a secure spike in some axon branches (d) and not others (e, f, FPs; DH S3 axons). Light evoked PAD (λ, 1.5xT, 10 ms prior) rescued most failed spikes (e, f) and sped up conduction in secure spikes (d). Box plots of FPs and spikes (g); *significant increase with light, *n =* 11 axons from (b); + significant reduction in light effect with 50 µM gabazine, *n =*11 axons from (b). h, Incidence of branches with failed DR-evoked spikes. *significant change with gabazine, χ-squared test, *P <* 0.05, for *n* = 45 control axons (Cntr) and *n* = 27 axons treated with gabazine. i-o, GABA_axo_ neurons imaged in S3 sacral spinal cord of GAD2//ChR2-EYFP//tdTom mice (j, l-o; green/red, merge yellow; *n* = 3 mice) or GAD2//ChR2-EYFP mice (k, green, dorsal horn, *n* = 5 mice), shown with raw images (maximum projection). Innervation of reconstructed neurobiotin filled sensory axons (gold in j and k, as in Fig. 1) by GABA_axo_ neurons (green; axon contacts labelled red in k) in dorsal horn. Nodes identified by Caspr and paranodal taper, sensory terminals by VGLUT1, GABA_axo_ terminals by VGAT, and axonal GABA_A_ receptors by the α5GABA_A_ subunit. ChR2-EYFP is mainly expressed on plasma membranes^120^, whereas tdTom is cytoplasmic, and so EYFP rings the tdTom labelling, especially evident on the soma and boutons (j, l-o). Regions in (l) expanded in (m-o).

Axon spike failure was voltage dependent: in branches with failing spikes (FPs) depolarizations that brought the axon closer to threshold enabled full DR-evoked spikes (via current injection, Fig. 2c*i*; or spontaneous depolarization, Fig. 2d). Also, in branches without spike failure at rest (secure spikes) a steady hyperpolarizing current induced spike failure (FP), with more branches failing with increasing hyperpolarization (Extended Data Fig. 2). With increasing hyperpolarization, nodes failed progressively more distal to the electrode, causing abrupt drops in the overall spike amplitude with each failure and a characteristic delay in the nodal spike prior to failure, and spike attenuation consistent with λ_S_ being about two internodal distances (∼90 µm; Extended Data Fig. 2a-d). Simulating spike propagation by applying a brief current pulse to mimic the current arriving from an upstream node (and FP) yielded similar results, with full spikes evoked at rest, but hyperpolarization leading to a spike delay and then failure (Extended Data Fig. 3). Large depolarizations inactivated spikes, though outside of the physiological range (> − 50 mV, Extended Data Fig. 3b-c).

### Nodal spike facilitation by GABA

Since sensory axons are tonically depolarized by spontaneous GABA activity^15^, we wondered whether this GABA aids propagation. Blocking extrasynaptic α5 GABA_A_ receptors (with L655708) or all GABA_A_ receptors (with gabazine) increased the incidence of spike failure (to ∼45% and 65%, respectively; Fig. 2f) and sensitivity to hyperpolarization (Extended Data Fig. 2e-h), without altering overall spike properties (Fig. 2g), implying that spike propagation is highly dependent on GABA receptors near enough to nodes to influence sodium channels (within λ_S_). Application of 5-HT to mimic natural brainstem-derived 5-HT also increased failure (Fig. 2f), likely via its indirect inhibition of GABA_A_ receptor activity^35^.

### Nodal spike facilitation by GABA_axo_ neuron activation

To examine whether GABA_axo_ neurons facilitate spike propagation, we expressed light-sensitive channelrhodopsin-2 (ChR2) in GAD2^+^ neurons in adult GAD2^CreER/+^;R26^LSL-ChR2-EYFP^ mice (termed GAD2//ChR2-EYFP mice, Fig. 3). A brief light pulse (5 - 10 ms) produced a long-lasting depolarization and spiking in these GABA_axo_ neurons (Fig. 3a), followed by a longer lasting GABA_A_-mediated depolarization (PAD) of proprioceptive axons at a monosynaptic latency that was blocked by gabazine (Fig. 3a-b). In these mice, spikes in proprioceptive axons failed with a similar incidence as observed in rats (Figs. 3c-h), but the light-evoked PAD prevented this failure (Fig. 3e-g), similar to direct depolarization. Occasionally, spikes were only partially rescued by PAD (< −60 mV spikes; Fig 3g), suggestive of PAD restoring conduction in some, but not all, nodes. In branches with secure non-failing spikes, light had minor effects (Fig. 3d), but blocking GABA_A_ receptors again increased the incidence of spike failure (Fig. 3h).

In GAD2//ChR2-EYFP or GAD2//ChR2-EYFP//tdTom mice the EYFP and tdTom reporters labelled GABAergic neurons (Fig. 3k; VGAT^+^, GAD2^+^ and VGLUT1^-^) residing near the central canal and throughout much of the dorsal horn (Fig. 3i-o, Extended Data Fig 4). These neurons densely innervated the dorsal horn (Fig. 3j,l,n; Extended Data Fig 4a), and less densely innervated both the ventral horn and dorsal columns with terminal boutons (Fig. 3l,m,o), allowing GABAergic innervation of sensory axons along their entire length. They made both synaptic and perisynaptic contacts along proprioceptive Ia sensory axons labelled either intracellularly with neurobiotin or peripherally with a viral vector, both at nodes and sensory axon terminals (Figs. 3k, and 1e, Extended Data Fig 4), confirming their identity as GABA_axo_ neurons. Consistent with the GABA_A_ receptor distribution, not all nodes were innervated by GABA_axo_ neurons (∼20%; Extended Data Fig. 4). However, nodes without GABA innervation often were connected to nearby (< λ_S_) short unmyelinated terminal branches and boutons innervated by GABA_axo_ neurons (Extended Data Fig 4c), like we report above for GABA_A_ receptors in the dorsal and intermediate regions of the spinal cord, enabling PAD to readily affect nodes. Overall, the majority of nodes were electrotonically close (< 90 µm, λ_S_) to GABA_axo_ contacts on a neighboring node or the node itself (98%) or on a nearby unmyelinated branch (77%) on the same axon (Extended Data Fig 4h), as for receptors. Furthermore, most nodes (95%) had nearby GABA_axo_ terminal boutons not contacting the axon (extrasynaptic contact; within 5 µm, GAD2^+^ or VGAT^+^; Extended Data Fig 4), providing extrasynaptic GABA.

Consistent with the predominantly dorsal GAD2^+^ innervation of nodes (Fig 3) and lack of terminal GABA_A_ receptors (Fig 1), PAD was evoked by light focused on the dorsal horn, but not on the ventral horn (Fig 3b, bottom). Furthermore, light-evoked PAD improved axon conductance even after silencing neuronal circuits with CNQX (50 µM, n = 11/11 axons in two mice, not shown, as in Fig 3e-g), indicating that nodal facilitation is caused directly by GABA_A_ receptors on the axon near the intra-axonal recording site and its nearby nodes (within ∼λ), and not via an indirect circuit. GABAergic neurons contacting ventral sensory terminals (previously termed GABApre neurons^6^) are likely a subpopulation of the GAD2^+^ neurons that contact nodes and terminals (GABA_axo_ neurons, Fig 3), since many GAD2^+^ neurons are located in superficial dorsal laminae (Fig 4l), unlike GABApre neurons located near the central canal^6^.

**Fig. 4.**
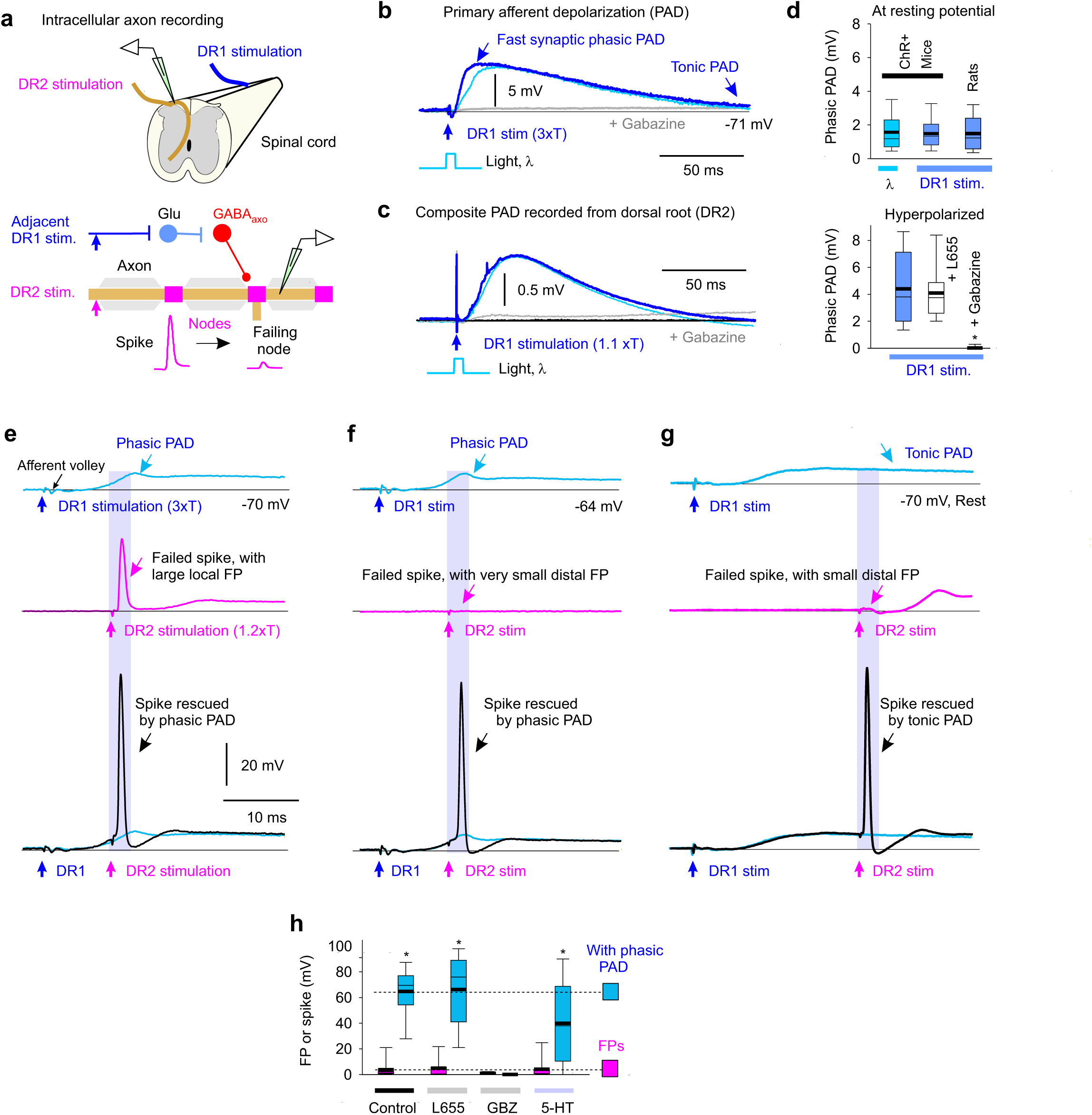
Sensory driven nodal facilitation. **a,** Experimental setup to indirectly activate GABA_axo_ neurons by DR stimulation (DR1) by trisynaptic PAD circuits (detailed in Extended Data Figs. 6-7) in ex vivo spinal cords of rats or GAD2//ChR2-EYFP mice. **b-c**, Depolarization (PAD) of a proprioceptive axon branch (**b**, intracellular in sacral S3 DH) or multiple axons in a DR (**c**, grease gap recording; sacral S3 DR; DR2) from stimulating the adjacent S4 DR (1.1 −3xT, 0.1 ms pulse; DR1; T: afferent volley spike threshold) or applying a light pulse to activate GABA_axo_ neurons (5 ms, 447nm, 0.7 mW/mm^2^, as in Fig. 3b), both blocked by gabazine (50 µM), in GAD2//ChR2 mouse. Thin line resting potential. **d**, Summary box plots of peak phasic PAD evoked in axons by adjacent DR stimulation (DR1) or light, at rest (top, n = 16 axons from 6 rats or n = 14 axons from 5 mice, the latter from Fig. 3b) and with hyperpolarization (−10 mV, bottom, same rats and axons), and effects of applied gabazine (50 µM; *n =* 14 axons from 5 rats) or L655708 (0.3 µM; *n =* 14 axons from 5 rats). *significant difference from pre-drug (blue, lower plot), *P <* 0.05. **e-g**, DR axon branches (sacral S3 DH) exhibiting spike failure (FPs, magenta) following stimulating their DR (S3 DR, 1.2xT, 0.1 ms; DR2) in rats at rest. Spikes rescued by PAD evoked by prior conditioning of adjacent DR (S4 or contralateral S3 DR, at 3xT; DR1). Rescue occurred with fast synaptic depolarizations (phasic PAD; **e-f**) and tonic depolarizations (tonic PAD, **g**), both for local FPs (large, **e**) or distal FPs (small, **f-g**). **h**, FP or spike heights before and during DR evoked phasic PAD (*n =* 17 axons from 11 rats) as in **e-f**, and actions of L655708 (*n* = 12 axons from 5 rats, 0.3 µM), gabazine (*n* = 12 axons from 5 rats, 50 µM) and 5-HT (*n* = 12 axons from 5 rats, 10 µM). *significant increase in spike with PAD, *P <* 0.05.

### Computer simulations of branch point failure and rescue by GABA

To establish that spike failure arises at the branch points where GABA can influence them, we generated a computer simulation of a proprioceptive sensory axon arbour in the spinal cord (Extended Data Fig. 5)^25^. With simulated DR stimulation, spike failure occurred distal to complex branch points (at nodes N2 and N3 in Extended Data Fig. 5a-b) that had associated increases in net conductance, which shunted the nodal currents. Simulated nodal GABA_A_ receptor activation rescued these failed spikes, with increasing GABA_A_ activation (g_GABA_) preventing more branch point failures (Extended Data Fig. 5c). Importantly, a single well placed GABA contact (at either N2 or N3, or on a nearby bouton) rescued conduction in an entire branch. In contrast, when we moved all these GABA_A_ receptors to the terminals, then their activation did not rescue failed spikes (Extended Data Fig. 5d). This is because GABA_A_-induced depolarizations (PAD) were attenuated sharply with distance (λ_S_ ∼90 µm); so only nodal, and not terminal, induced PAD was visible at the dorsal columns (Extended Data Fig. 5a,g-h), in agreement with previous terminal recordings^15^.

### Spike facilitation by sensory evoked GABA_axo_ activity

We next examined whether natural activation of GABA_axo_ neurons affects proprioceptive axon conduction (Fig 4). GABA_axo_ neurons are indirectly activated by sensory activity via two variants of a trisynaptic circuit, where sensory axons drive excitatory neurons that activate GABA_axo_ neurons and cause PAD: one driven by cutaneous afferents and the other by proprioceptive afferents (Extended Data Figs. 6 and 7)^15^. As expected, following DR stimulation these circuits caused fast synaptic and slower extrasynaptic GABA_A_ receptor mediated depolarizations of proprioceptive axons (termed sensory-evoked phasic PAD and tonic PAD, respectively^15^) that were blocked by GABA_A_ antagonists, and mimicked by optogenetic activation of GABA_axo_ neurons (Fig. 4a-d).

Like with direct GABA_axo_ activation, spike propagation failure was prevented by sensory-evoked phasic PAD, regardless of whether the failure was spontaneous (Figs. 4e-f,h), 5-HT-induced (Fig. 4h), or repetition-induced (Extended Data Fig. 7b-f). The latter is particularly important because sensory axons naturally fire at high rates, where they are vulnerable to spike failure (Fig. 2e-f). This action of phasic PAD was abolished by gabazine but not L655708, supporting a synaptic origin (Fig. 4h). Slow extrasynaptic GABAergic depolarization (tonic PAD; L655708-sensitive^15^) further facilitated spike propagation (Fig. 4g), especially as it built up with repeated DR stimulation (at 1 Hz; Extended Data Fig. 5b). Cutaneous (Extended Data Fig. 6), proprioceptive (Extended Data Fig. 7) or mixed afferent (Fig. 4e-h) -evoked PAD all helped prevent spike failure.

In secure non-failing axon branches sensory-evoked PAD (or optogenetic GABA_axo_ activation) sped up the spikes and lowered their threshold (rheobase current; Fig. 3d and Extended Data Fig. 8a-d), as predicted from computer simulations (Extended Data Fig. 5e). Importantly, spike height was only slightly reduced during PAD (∼1% or 1 mV) indicating that GABA_A_ receptor conductances have minimal shunting action on nearby spikes (Fig. 3 and Extend Data Fig. 8a-d).

### Failure of axon conduction to motoneurons and rescue by PAD

To quantify the overall failure of spikes to conduct from the DR to the sensory axon terminals we measured whether axon branches not conducting during failure were *not* refractory to subsequent stimulation with a microelectrode in the ventral horn (Extended Data Fig. 9). This method indicated that about 50 – 80% of sensory axons failed to conduct to their ventral terminals under resting conditions, especially in long axons, whereas sensory-evoked PAD decreased failure to < 30%. Similar conclusions were reached by directly recording the extracellular afferent volley in the ventral horn produced by the spikes propagating from a DR stimulation to the motoneurons, which was consistently increased by PAD (Extended Data Fig. 10).

### Facilitation of sensory feedback to motoneurons by GABA_A_ receptors

To examine the functional role of GABA in regulating sensory feedback to motoneurons, we recorded monosynaptic excitatory postsynaptic potentials (EPSPs) from motoneurons in response to proprioceptive sensory axon stimulation (Fig. 5). This EPSP was inhibited by optogenetically silencing GABA_axo_ neurons with light in mice expressing archaerhodopsin-3 (Arch3, induced in GAD2^CreER/+^;R26^LSL-Arch3-GFP^ mice; abbreviated GAD2//Arch3, Fig. 5a-b,d), consistent with a tonic GABA_A_ receptor tone facilitating spike transmission in axons. Likewise, the EPSP was reduced when sensory axon conduction was reduced by blocking endogenous GABA_A_ receptor tone with antagonists, despite increasing motoneuron and polysynaptic reflex excitability (the latter minimized with APV, Fig. 5c,d). GABA_B_ antagonists slightly increased the EPSP, suggesting a tonic GABA_B_-mediated presynaptic inhibition (Fig. 5d), though much smaller than the tonic GABA_A_-mediated nodal facilitation that dominates when all GABA was reduced (in GAD2//Arch3 mice).

**Fig. 5.**
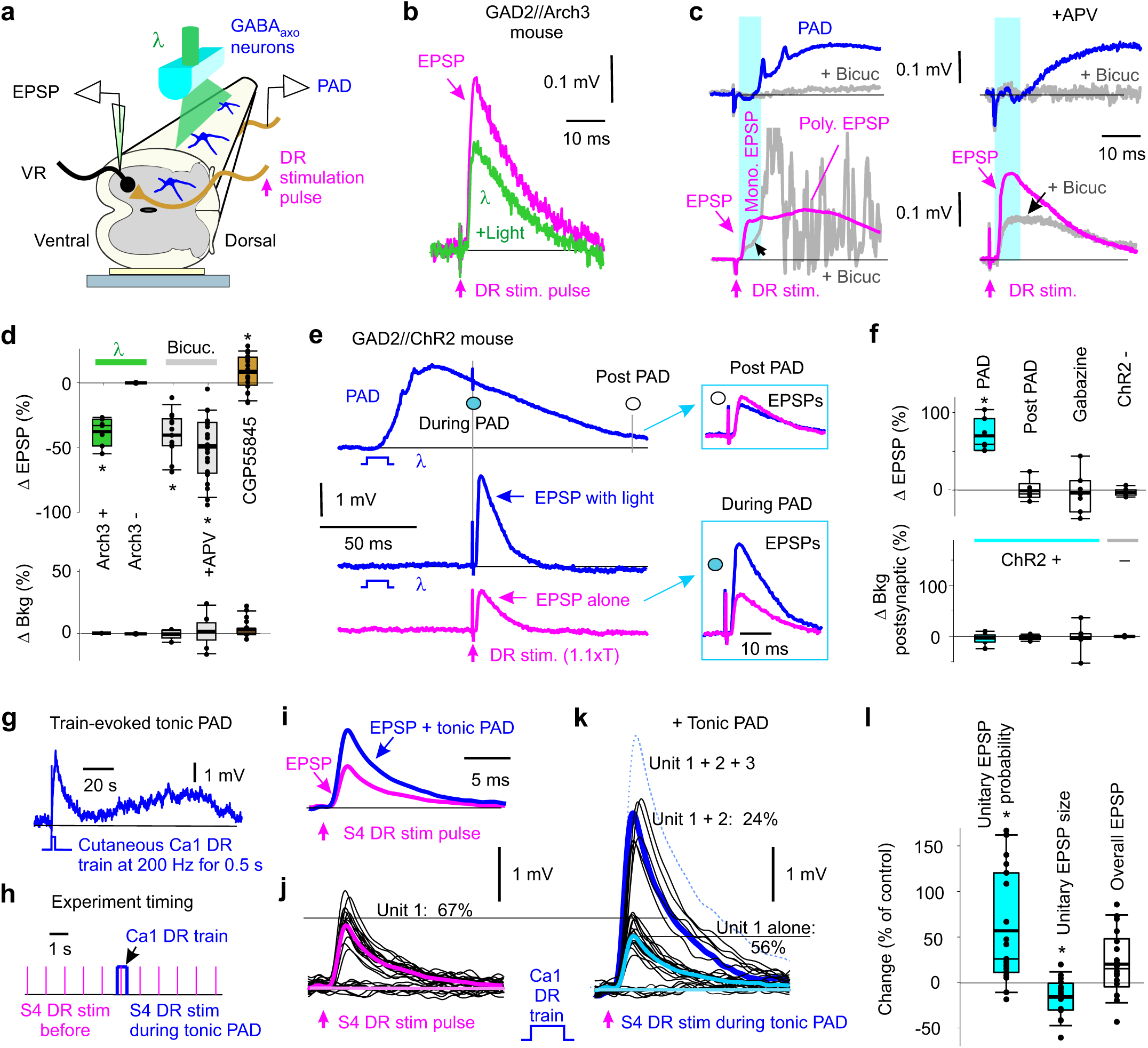
Facilitation of monosynaptic sensory transmission by GABA. **a,** Ex vivo recording from motoneurons while illuminating GABA_axo_ neurons with light λ. **b-d**, Composite monosynaptic EPSP from motoneuron pool (recorded in sacral S4 VR) evoked by a DR stimulation pulse alone (S4 DR, 0.1 ms, 1.1xT, magenta; T: EPSP threshold). Actions of optogenetic silencing GABA_axo_ neurons with light (**a-b**, 532nm, 5 mW/mm^2^, 80 ms, in *n =* 7 composite EPSPs from 4 GAD2//Arch3 mice), blocking GABA_A_ receptors (**c**, with bicuculline or gabazine, 50 µM, with and without NMDA antagonist APV, 50 µM; *n =* 23 from 10 mice and 13 EPSPs from 5 mice, respectively; rats similar, not shown), or blocking GABA_B_ receptors (**d**, CGP55845, 0.3 µM, *n =* 20 composite EPSPs in 20 mice). PAD shown for reference, recorded on S3 DR (**c,** top). Summary box plots of changes in EPSP and background postsynaptic activity (Bkg, over 10 ms prior to EPSP) with light or drugs, and with Arch3+ and Arch3-mice (**d**). * significant change, Δ, *P <* 0.05. GAD2//Arch3 mice are VGAT+, not shown. **e-f**, Composite EPSP (evoked in S4 or S3 motoneurons, as in **a-b**) before, during and post PAD (recorded simultaneously on S3 DR) evoked by light activation of GABA_axo_ neurons (10 ms, 1.1xT, 447nm, 0.5 mW/mm^2^, 60 ms and 140 ms pre EPSP, ISI) in GAD2//ChR2 mice. Box plots of changes in EPSP and Bkg (10 ms prior) with light in ChR2^+^ mice without and with gabazine (50 µM, during PAD), and in ChR2^-^ mice (60 ms ISI). * significant change, *P <* 0.05, for each *n =* 7 composite EPSPs from 7 mice each. **g**, Tonic PAD (L655708 sensitive) recorded in sacral S4 proprioceptive Ia axon in response to 0.5 s, 200 Hz DR stimulation train applied to the largely cutaneous Ca1 DR of caudal cord (3xT, DR2) in rat. **h-i**, Average EPSP in S4 motoneuron (intracellular recording, EPSP evoked by S4 DR stimulation at 3 s intervals used for average; DR1) before and during tonic PAD (**i**) evoked by the brief DR train of (**h)**, at matched postsynaptic potentials. **j-k**, Individual trials used to make EPSP averages in (**i**) (at 1 s intervals, **h**), with large all or nothing unitary EPSPs (thick lines unitary averages; dotted single occurrence of Unit 3). Lowpass filtered at 3 kHz. **l**, Changes in unitary EPSP probability and size, and overall EPSP with tonic PAD. Box plots. * significant change, *P <* 0.05, *n =* 18 motoneurons from 5 rats.

Consistent with GABA_A_ receptors and PAD facilitating axon conduction, the monosynaptic EPSP was facilitated during, but not after, depolarizing proprioceptive axons (evoking PAD) with an optogenetic activation of GABA_axo_ neurons in GAD2//ChR2 mice (10 ms *light conditioning stimulation*; Fig. 5e-f). The EPSP was also facilitated by naturally activating GABA_axo_ neurons by a *sensory conditioning stimulation* (Extended Data Fig. 11), including with a conditioning stimulation of cutaneous and/or proprioceptive afferents (Extended Data Fig. 11a,b,e). The latter indicates that proprioceptive activity primes subsequent proprioceptive reflex transmission (*self-facilitation*). GABA_A_ receptor antagonists (gabazine), but not GABA_B_ antagonists (CGP55845), blocked the EPSP facilitation with sensory (Extended Data Fig. 11e) or light (Fig. 5f) conditioning.

The facilitation of the EPSP by conditioning-evoked PAD arose from axonal GABA_A_ receptors, rather than from postsynaptic actions on the motoneurons, since it occurred with weak conditioning stimuli that produced only a transient background postsynaptic depolarization that terminated before the EPSP testing (at 60 ms; Figs. 5e, Extended Data Fig. 11b,g), followed by a slight hyperpolarization that if anything would reduce the EPSP (shunting the synaptic current, Extended Data Fig. 11h). Increasing the DR conditioning intensity produced large background depolarizing conductances in the motoneurons during the EPSP testing, which led to postsynaptic inhibition of the EPSP (shunting inhibition; Extended Data Fig. 11d,g) and post activation depression, masking the effect of nodal facilitation. Importantly, sometimes PAD itself induced afferent spikes (Extended Data Fig. 8e; termed DRR spikes), and following these spikes, the EPSP was always smaller than when these spikes were not present (*n =* 8/8 mice, not shown). This is because these DRR spikes themselves triggered EPSPs, leading to a post activation depression, as noted by Eccles^8^, and thus we minimized DRR activity by keeping the conditioning-evoked PAD small.

Sensory conditioning was particularly effective when it was repeated to mimic natural firing, which increased tonic PAD for minutes (Fig. 5g). This facilitated the EPSP for ∼3 min after a brief fast DR repetition (200 Hz, 0.5 s conditioning, Fig. 5i, Extended Data Fig. 11e, *Tonic*), and ∼1 min after slower repetition (0.1 Hz, 2 min conditioning, Extended Data Fig. 11e, *After effect*), both long outlasting postsynaptic effects from each conditioning pulse (< 1 s). This was blocked by L655708 or gabazine (Extended Data Fig. 11e). Interestingly, optogenetic activation of GABA_axo_ neurons did not produce a similar after effect, consistent with this tonic PAD and associated nodal facilitation being mediated by extrasynaptic GABA spillover from other sources of GABA^15^.

### Increases in the probability of unitary EPSPs by nodal facilitation

We often noticed large all-or-nothing EPSPs (unitary EPSPs) spontaneously fluctuating on and off during repeated EPSP testing, leading to discrete large changes in the total EPSP size and time course (Fig. 5j-k). We thought this might be due to spontaneous branch point failures, rather than quantal changes in transmitter release that produce much smaller fluctuations^36^, as previously suggested^29^. Indeed, when we increased the axon conduction by activating the GABA_axo_ neurons and PAD (via a cutaneous conditioning train) the *probability* of unitary EPSPs occurring increased (Fig. 5k-l), and this sometimes recruited further large unitary EPSPs (Fig. 5k). In contrast, the *size* of the underlying unitary EPSP was not increased by this conditioning (Fig. 5j-l), ruling out decreases in terminal presynaptic inhibition or postsynaptic inhibition contributing to the increased overall EPSP (Fig. 5i,l). Likely the smaller unitary EPSP arose from GABA_B_ receptors causing presynaptic inhibition and the increased unitary EPSP probability arose from GABA_A_ causing nodal facilitation.

### Facilitation of sensory feedback to motoneurons by GABA_axo_ neurons in awake mice

To determine whether GABA_axo_ neurons increase sensory feedback to motoneurons in *awake* mice we activated these neurons with light applied through a window chronically implanted over the spinal cord of GAD2//ChR2 mice (Fig. 6), and assessed the monosynaptic reflex (MSR) recorded in tail muscles in response to nerve stimulation (counterpart of EPSPs; Fig 6a-c). As expected, the MSR was facilitated by a conditioning light pulse, but only during, and not after, the expected time of phasic PAD induced on sensory axons (Fig. 6b-d,j). This light-induced *facilitation* occurred both at rest and when there was a background voluntary contraction, with the latter matched with and without light, again ruling out postsynaptic depolarization related differences in MSR (Fig. 6d). Light alone caused a brief pause in ongoing EMG (∼30 ms post-light; Fig. 6b), indicative of postsynaptic inhibition, which masked nodal facilitation at short intervals.

**Fig. 6.**
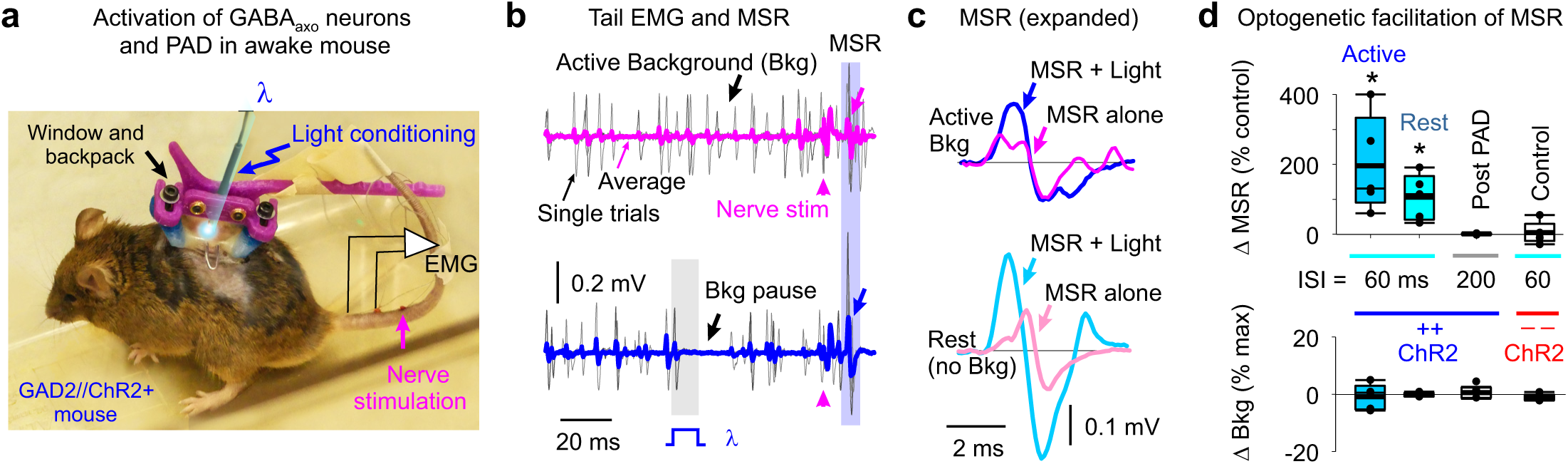
Facilitation of reflexes in awake mice. **a,** Recording tail muscle EMG and evoking monosynaptic reflexes (MSR) with tail nerve stimulation (1.1xT, 0.2 Hz; T: reflex threshold), while activating GABA_axo_ neurons (PAD) with light (λ = 447 nm, 10 ms pulse, 1.5xT, 5 mW/mm^2^) in GAD2//ChR2 mice. **b-c**, Effect of light pulse λ on active background EMG (Active Bkg condition in **b**) and the MSR evoked 60 ms later, the latter expanded in (**c**). MSR tested with (**c**, top; Active Bkg, 30% max) and without (**c**, bottom; Rest) background EMG. Thin black lines in (**b)** are individual trial examples at 10 s intervals (0.1 Hz); thick lines: averages. **d**, Changes in MSR with light activation of GABA_axo_ neurons at matched postsynaptic background (Bkg measured over 20 ms prior to MSR; lack of change in Bkg). Measured in active and resting (no Bkg) states, in ChR2+ and ChR2-mice (rest only), and during (60 ms ISI) and post PAD (200ms ISI at rest only). ISI: interstimulus interval. Box plots. * significant change, *P <* 0.05, *n =* 5 mice each.

### Facilitation of sensory feedback to motoneurons during PAD in awake rat and humans

When we evoked PAD by cutaneous sensory stimulation in awake rats (Extended Data Fig. 12) or humans (Extended Data Fig. 13) the MSR reflex recorded in the tail or lower leg (soleus) was again increased and L655708 sensitive, consistent with the increased EPSPs seen in rats in vitro (Fig 5). This generalizes our main finding to lumbar spinal circuits that control the leg. Importantly, the probability of a single motor unit (MU) contributing to the human MSR was increased by cutaneous conditioning (Extended Data Fig. 13f*i-ii*). This occurred without an increase in the estimated EPSP amplitude or rise time (PSF; see Methods; Extended Data Fig. 13F*iii*) or motoneuron depolarization prior to the MSR testing (Extended Data Fig. 13F*iv*), consistent with an increased probability of unitary EPSPs and decreased branch point failure, as in rats (Fig. 5).

## Discussion

Following the pioneering studies of Eccles on inhibition of the monosynaptic connection from sensory axons to motoneurons^8^, the concept of presynaptic inhibition of axon terminals has stood as a cornerstone of our understanding of mammalian brain and spinal cord function^9^. While presynaptic inhibition has never been directly confirmed in these sensory axons, recordings from invertebrate sensory axons have established the idea that terminal GABA_A_ receptor-mediated depolarizations (PAD) can cause conductance increases or sodium channel inactivation that inhibit transmitter release^21, 23^. Thus, our finding that GABA_A_ receptors are located too far (relative to the short λ_S_) from the axon terminals to influence terminal depolarizations or cause presynaptic inhibition of transmitter release onto motoneurons had not been anticipated. However, in retrospect a lack of terminal GABA_A_ receptors had previously been noted^15, 24^, as had a lack of ventral terminal depolarization during GABA_axo_ neuron activity^15^ (Extended Data Table 1). Direct recordings from terminal boutons of other axon types in the mammalian brain, such as the Calyx of Held, have shown that terminal GABA_A_ or glycine receptors sometimes cause presynaptic inhibition, and at other times facilitate transmitter release, depending on their action on terminal calcium and potassium channels^20, 21, 37^. However, these terminal actions of GABA are very different from the actions of GABA near the nodes that we have uncovered. That is, GABA_A_ receptors or direct current injections near sodium channels help prevent conduction failure at nodes downstream to a branch point by depolarizing the failing nodes closer to spike threshold (nodal facilitation; summarized in Fig 7), similar to the relief from spike failure seen by depolarization in leech^31^. Neuronal circuits that control GABA_axo_ neurons help produce this PAD and associated nodal facilitation, as does tonic PAD produced by spillover of GABA from any source during repeated cutaneous stimulation^15^, ultimately inducing a tonic GABAergic facilitation of nodal conduction (ArchT and gabazine-sensitive). The profound inhibitory action of GABA_A_ receptors antagonists or optogenetic inhibition of GABA_axo_ neurons on PAD and sensory transmission demonstrates that without axonal GABA, and associated depolarizations (PAD), nodal spike transmission fails in many central branches of large myelinated proprioceptive sensory axons, leaving large silent branches, depending on the branching structure (Extended Data Fig 5) and prior history of activity (firing frequency; see also^31^). This inhibitory action is not due to GABA_A_ receptor antagonists working on neuronal circuits that somehow indirectly affect PAD, because tonic PAD and nodal facilitation are resistant to blocking all circuit activity with TTX or CNQX, and blocked by subsequent application of these GABA_A_ antagonists^15^.

**Fig. 7.**
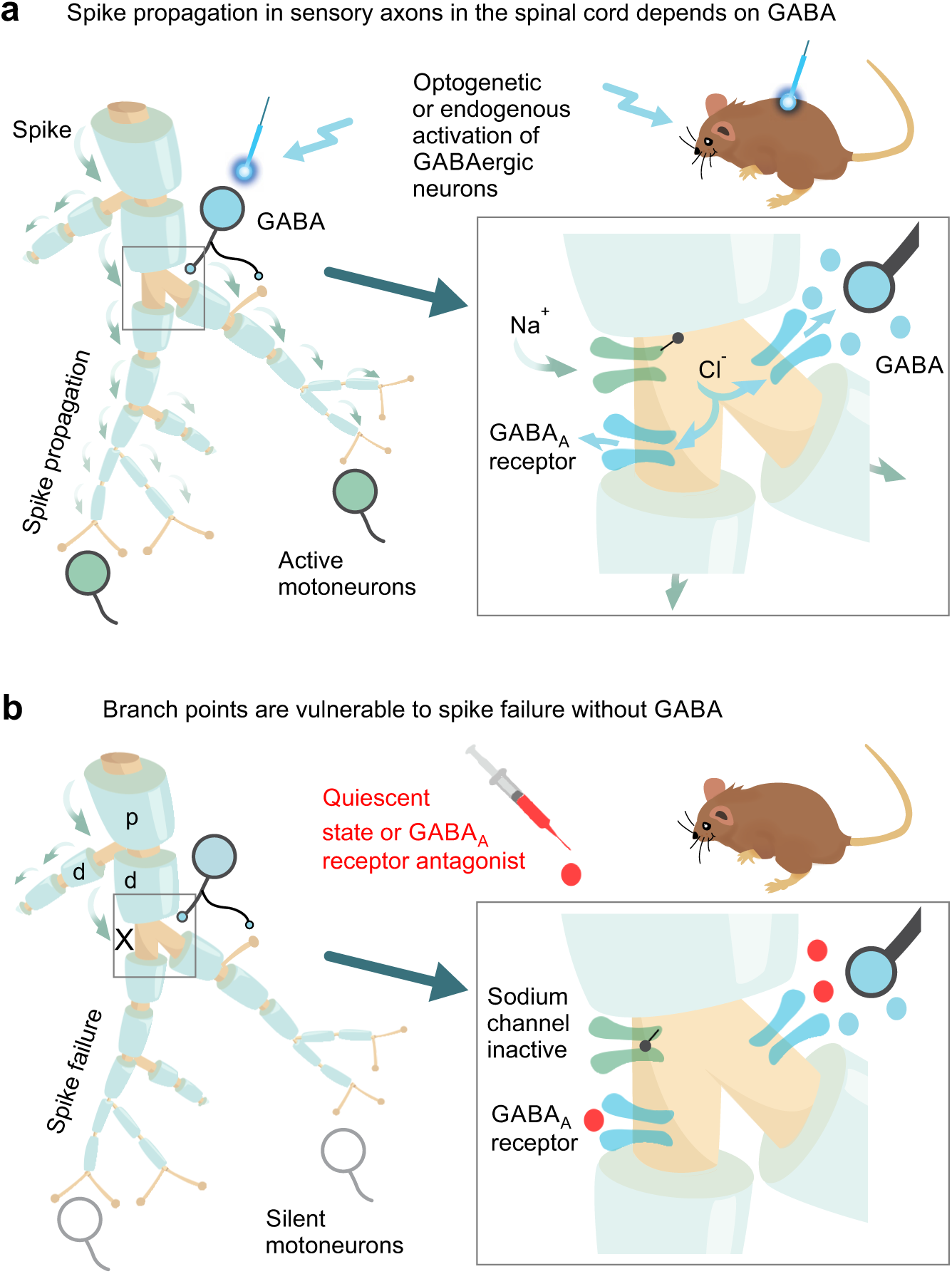
Graphical summary of nodal facilitation. **a,** Spike propagation (green arrows) in myelinated proprioceptive afferents is facilitated by GABA_A_ receptors at or near nodes, depolarizing axons closer to spike threshold via outward chloride currents. GABAergic neurons (GABA_axo_) provide synaptic or perisynaptic innervation of nodes and nearby terminal boutons (right), and optogenetic or endogenous activation of these neurons increases sensory transmission to motoneurons. **b**, Without this GABA_A_ receptor activity, spikes fail to propagate into some of the branches of proprioceptive sensory axons in the spinal cord, due to branch point failure. Failure is initiated when a parent branch, p, cannot provide enough current to drive the daughter branches, d, with at least one of the daughters failing, especially if that branch has further branch points, producing a large conductive load. Overall, nodal facilitation by GABA_axo_ neurons allows selective regulation of conduction in individual branches, increasing the computational complexity of sensory circuits.

Our computer simulations demonstrate that spike conduction failure is only initiated at particularly vulnerable branch points, as previously suggested^31^, and thus only nodes at or downstream of these failure points require GABAergic facilitation, consistent with the observation that GABA_A_ receptors and GABAergic contacts (VGAT^+^, GAD2^+^) are only at or near a faction of nodes (Fig 1)^25^. For this, GABA_A_ receptors need only be within 90 µm of the node (λ_S_), at another node or even on one of the short unmyelinated terminal branches in dorsal and intermediate regions connected to the node. Ultrastructural imaging has demonstrated that in the spinal cord GABA receptors often lack presynaptic contacts^24^, consistent with many GABA receptors being activated extrasynaptically from spill over of nearby GABA, and accounting for the fewer number of nodes with GABAergic contacts than nodes with GABA receptors. Limitations in our confocal imaging leave many open questions for future study, including the precise spatial relation of GABA receptors to sodium channels, myelin and intracellular structures. We cannot rule out the possibility that the oligodendrocytes at the paranode influence GABAergic control of the axon, since they express GABA receptors and GABA^38^, and GABA_axo_ contacts sometimes straddle the paranodal myelin and the node, maybe forming a tripartite neuron-glia-axon arrangement. Nevertheless, our observations of GABA facilitating conduction to these dorsal branches can only be accounted for by nodal facilitation, regardless of the details of how GABA innervates these nodes or nearby branches. The concept of nodal facilitation that we describe here may generalize to other large central axons, such as pyramidal cells, that are innervated by GABAergic neurons, branch extensively (and so may fail), and have depolarizing actions of GABA^19, 21, 39, 40^, allowing selective recruitment of specific axon branches and functional pathways, especially for high frequency firing.

Sensory driven GABA_axo_ circuits and associated PAD are experimentally convenient, since they allow us to estimate how sensory transmission to motoneurons is modulated with GABA_axo_ neuron activity in humans. Specifically, the expected time-course of PAD evoked by cutaneous conditioning is associated with a potent reflex facilitation in humans and awake rodents. This suggests a substantial ongoing spike failure prior to facilitation that can be alleviated by GABA_axo_ activity (PAD). Indeed, we found that during PAD the *probability* of EPSPs occurring (and MU firing) is increased without changing the EPSP amplitude, as estimated by PSFs in humans. The latter rules out changes in presynaptic inhibition with PAD that *grades* the EPSP size, including ruling out previous arguments that MSR facilitation by cutaneous conditioning is due to a removal of presynaptic inhibition^41, 42^.

A pressing question that remains is how can nearly a century of research on sensory transmission and presynaptic inhibition be reconciled with GABA-mediated nodal facilitation and reflex facilitation (summarized in Extended Data Table 1)? Sensory axon conduction failure has repeatedly been noted from indirect observations^29–31, 43^, but GABA_A_ receptors and PAD were previously thought to cause, rather than prevent, conduction failure^30^, even though computer simulations showed physiological GABA levels unlikely to block spike propagation^25^, as we confirmed. Furthermore, the fundamental assumption that GABA_A_ receptors cause presynaptic inhibition that reduces transmitter release from sensory axons was from the outset circumspect, based mainly on the observation that a conditioning stimulation on a flexor nerve caused an inhibition of the MSR evoked in extensor muscles that was somewhat correlated to the time-course of PAD caused by this conditioning in flexor afferents^8^. However, in retrospect this PAD is too brief to account for the much longer (up to 1 s) inhibition caused by this conditioning^8, 10, 44^, and GABA_B_ receptor antagonists block much of this MSR inhibition^16, 44^. This fits with GABA_B_ receptors being at the terminals (Fig. 1) and being primarily responsible for presynaptic inhibition in proprioceptive axons, as in other neurons^20, 21^, though further study of GABA_B_ receptor function is now needed. This predominant GABA_B_ action in proprioceptive axon terminals does not rule out GABA_A_-mediated presynaptic inhibition in other sensory axons that have terminal GABA_A_ receptor expression, such as cutaneous Aβ afferents^15^.

Anatomical studies suggest that GABA_axo_ neuron activation is likely accompanied by some postsynaptic inhibition, since most GABA_axo_ contacts on afferent terminals also contact motoneurons, in a triad^7, 45^. Indeed, we find that GABA_axo_ neuron activation produces an inhibition of motoneurons (Fig. 6b) and associated MU firing that masks, and at times overwhelms, the facilitation of the MSR by GABA_A_ receptors (as with muscle vibration; Extended Data Fig. 13), and thus is readily mistaken for presynaptic inhibition. The argument that presynaptic inhibition with conditioning should be evident from reductions in the EPSP without changing its time course^46^ now seems untenable, especially as unitary EPSPs differ markedly in shape and conditioning increases the number of unitary EPSPs contributing to the EPSP, as different axon branches are recruited (Fig. 5k)^29^.

Early on Barron and Matthews^22^ and later others^6, 15^ established that sensory-evoked PAD (or light-evoked) excites axons by directly inducing spiking, including spikes in the sensory axons mediating the MSR itself, raising a further contradiction with presynaptic inhibition. While these PAD-triggered spikes only sometimes fully propagate antidromically out the DR^47^, they are more likely to conduct orthodromically^15^ where they activate the motoneurons^6, 8^, making these axons and their motoneuron synapse refractory to subsequent testing^8^. This contributes to a long-lasting *post-activation depression* of the MSR pathway that is GABA_A_-mediated (sensitive to GABA_A_ antagonists, like PAD) and is thus readily mistaken for GABA_A_-mediated presynaptic inhibition^6, 16, 17^. While such PAD-triggered spikes are fundamentally excitatory, it remains an open question as to the physiological role of their induced post-activation depression.

Functionally, nodal facilitation and regulation of branch point failure by GABA_axo_-driven GABA_A_ receptors acts like a global switching system that recruits entire silent sensory or motor circuits. This works in concert to terminal presynaptic inhibition (including GABA_B_ receptor action) that locally fine tunes reflex gains, to optimize the stability and compliance of movement^3, 6^. The direct activation of GABA_axo_ neurons and associated PAD by cortical (CST) and spinal (CPG) circuits^11, 12, 48^, and inhibition by the brainstem (e.g. 5-HT)^35, 49^, suggests that nodal facilitation is under explicit central control during reaching and locomotion. The widespread action of PAD, occurring simultaneously over many spinal segments,^10, 15^ implies that nodal facilitation acts over large regions of the spinal cord to ready sensory axons for action during cortical, spinal or sensory evoked activity, reminiscent of the Jendrassik maneuver^50^, ensuring that adequate sensory feedback aids postural stability and walking. More generally, our results imply that each axonal branch point has the capacity to function separately, depending on its GABAergic innervation, increasing the complexity of sensory processing in the spinal cord.

## Methods

### Adult mice, rats and humans used

Recordings were made from large proprioceptive group Ia sensory afferents, GABAergic neurons, motoneurons and muscles in adult mice (2.5 – 6 months old, both female and male equally; strains detailed below) and rats (3 - 8 months old, female only, Sprague-Dawley). All experimental procedures were approved by the University of Alberta Animal Care and Use Committee, Health Sciences division. Recordings were also made from the soleus muscle of neurologically intact adult humans (female and male equally), aged 21 to 58, with written informed consent prior to participation. Experiments were approved by the Health Research Ethics Board of the University of Alberta (Protocols 00023530 and 00076790) and conformed to the Declaration of Helsinki. No effects of sex were noted and data from both sexes were combined for analysis.

### Mice used for optogenetics and imaging

We evaluated GABAergic neurons in a strain of mice with Cre expressed under the endogenous *Gad2* promotor region. *Gad2* encodes the Glutamate decarboxylase 2 enzyme GAD2 (also called GAD65), which is unique to axoaxonic contacting GABAergic neurons that project to the ventral horn, whereas all GABAergic neurons express GAD1^5^. These GAD2^+^ neurons were activated or inhibit optogenetically using channelrhodopsin-2 (ChR2)^51, 52^ or archaerhodopsin-3 (Ach3)^53, 54^, respectively. The following mouse strains were employed (Extended Data Table 2):

(1) *Gad2^tm1(cre/ERT2)Zjh^* mice (abbreviated Gad2^CreER^ mice; The Jackson Laboratory, Stock # 010702; CreER^T2^ fusion protein expressed under control of the endogenous *Gad2* promotor)^55^,
(2) B6;129S-*Gt(ROSA)26Sor^tm32(CAG-COP4*H134R/EYFP)Hze^* mice (abbreviated R26^LSL-ChR2-EYFP^ mice; The Jackson Laboratory, Stock # 012569; ChR2-EYFP fusion protein expressed under the R26::CAG promotor in cells that co-express Cre because a loxP-flanked STOP cassette, LSL, prevents transcription of the downstream ChR2-EYFP gene)^56^,
(3) B6.Cg-*Gt(ROSA)26Sor^tm14(CAG-tdTomato)Hze^* and B6.Cg-*Gt(ROSA)26Sor^tm9(CAG-tdTomato)Hze^* mice (abbreviated R26^LSL-tdTom^ mice; The Jackson Laboratory, Stock # 007914 and #007909; tdTomato fluorescent protein expressed under the R26::CAG promotor in cells that co-express Cre)^57^,
(4) B6;129S-*Gt(ROSA)26Sor^tm35.1(CAG-aop3/GFP)Hze^* mice (abbreviated R26^LSL-Arch3-GFP^ mice; The Jackson Laboratory Stock # 012735; Arch3-GFP fusion protein expressed under the R26::CAG promotor in cells that co-express Cre)^56^,
(5) B6;129S-*Slc17a7^tm1.1(cre)Hze^* mice (abbreviated VGLUT1^Cre^ mice; The Jackson Laboratory, Stock # 023527; Cre protein expressed under control of the endogenous *Vglut1* promotor; spinal cords kindly donated by Dr. Francisco J. Alvarez)^58^ and
(6) EIIa-cre x Gabra5-floxed in a C57BL/6 mouse background, with cre bred out to yield α5 GABA_A_ receptor knockout mice (termed Gabra5 KO mice).

Heterozygous GAD2^CreER^ mice (i.e., GAD2^CreER/+^ mice) were crossed with homozygous reporter strains to generate GAD2^CreER/+^; R26^LSL-ChR2-EYFP^, GAD2^CreER/+^; R26^LSL-tdTom^ and GAD2^CreER/+^; R26^LSL-Arch3-GFP^ mice that we abbreviate: GAD2//ChR2, GAD2//tdTom and GAD2//Arch3 mice. Offspring without the GAD2^CreER^ mutation, but with the effectors ChR2, Arch3 or tdTom were used as controls. We also used mice bred by crossing homozygous VGLUT1^Cre^ mice with R26^lsl-tdTom^ reporter mice to obtain mice with VGLUT1 labelled sensory axons^59^.

CreER is an inducible form of Cre that requires tamoxifen to activate ^60^, which we applied in adult mice to prevent developmental issues of earlier induction of Cre. Specifically, mice were injected at 4 - 6 weeks old with two doses of tamoxifen separated by two days, and studied > 1 month later, long after washout of tamoxifen. Each injection was 0.2 mg/g wt (i.p.) of tamoxifen dissolved in a corn oil delivery vehicle (Sigma C8267). These tamoxifen-treated mice were denoted GAD2//ChR2+ and GAD2//Arch3+, and non treated mice were used as controls and denoted GAD2//ChR2- and GAD2//Arch2-. For all mice, genotyping was performed according to the Jackson Laboratories protocols by PCR of ear biopsies using primers specific for the appropriate mutant and wild type alleles for each of the mouse lines (see Extended Data Table 2 for primer details).

### Ex vivo recording from axons and motoneurons in whole adult spinal cords

Mice or rats were anaesthetized with urethane (for mice 0.11 g/100 g, with a maximum dose of 0.065 g; and for rats 0.18 g/100 g, with a maximum dose of 0.45 g), a laminectomy was performed, and then the entire sacrocaudal spinal cord was rapidly removed and immersed in oxygenated modified artificial cerebrospinal fluid (mACSF), as detailed previously ^61–63^. This preparation is particularly useful as the small sacrocaudal spinal cord is the only portion of the adult spinal cord that survives whole ex vivo, allowing axon conduction to be assessed along large distances. Furthermore, this segment of cord innervates the axial muscles of the tail that are readily assessable for reflex recording in awake animals, and has proven to be a useful model of motor function in normal and injured spinal cords, with very similar spinal circuitry, reflex and motoneuron properties to those seen in the hindlimb of other preparations, including having reciprocal inhibition, Ia afferent innervation of muscle spindles and monosynaptic reflexes^62, 64–66^. Interestingly, the rat motoneuron firing rates in the sacral region are more similar to those in human hindlimb motoneurons than the much higher firing rats seen in rat hindlimb motoneurons, and thus the sacral cord has proven to be a useful model of lumbar motoneuron function in humans^67^. Spinal roots were removed, except the sacral S3, S4 and caudal Ca1 ventral and dorsal roots on both sides of the cord. After 1.5 hours in the dissection chamber (at 20° C), the cord was transferred to a recording chamber containing normal ACSF (nACSF) maintained at 23 - 32°C, with a flow rate > 3 ml/min. A one-hour period in nACSF was given to wash out the residual anaesthetic prior to recording, at which time the nACSF was recycled in a closed system. The cord was secured onto tissue paper at the bottom of a rubber (Silguard) chamber by insect pins in connective tissue and cut root fragments. The dorsal surface of the cord was usually oriented upwards when making intracellular recording from afferents in the dorsal horn, whereas the cord was oriented with its left side upwards when making recordings from motoneurons or afferent terminals in the ventral horn. The laser beam used for optogenetics was focused vertically downward on the GAD2 neurons, as detailed below.

### Optogenetic regulation of GABA_axo_ neurons

The GAD2//ChR2 or GAD2//Arch3 mice were used to optogenetically excite or inhibit GAD2+ neurons (with 447 nm D442001FX and 532 nM LRS-0532-GFM-00200-01 lasers from Laserglow Technologies, Toronto), respectively, using methods we previously described ^68^. Light was derived from the laser passed through a fibre optic cable (MFP_200/220/900-0.22_2m_FC-ZF1.25 and MFP_200/240/3000-0.22_2m_FC-FC, Doric Lenses, Quebec City) and then a half cylindrical prism the length of about two spinal segments (8 mm; 3.9 mm focal length, Thor Labs, Newton, USA,), which collimated the light into a narrow long beam (200 µm wide and 8 mm long). This narrow beam was usually focused longitudinally on the left side of the spinal cord roughly at the level of the dorsal horn, to target the epicentre of GABA_axo_ neurons, which are dorsally located (Fig. 3). ChR2 rapidly depolarizes neurons ^52^, and thus we used 5 – 10 ms light pulses to activate GABA_axo_ neurons, as confirmed by direct recordings from these neuron (see below). Light was always kept at a minimal intensity, 1.1x T, where T is the threshold to evoke a light response in sensory axons, which made local heating from light unlikely. Arch3 is a proton pump that is activated by green light, leading to a hyperpolarization and slowly increased pH (over seconds), both of which inhibit the neurons ^52, 69^. Thus, we used longer light pulses (∼200 ms) to inhibit GABA_axo_ neurons.

To directly confirm the presence of functional ChR2 expression in GABA_axo_ neurons of GAD2//ChR2 mice we recorded from them with similar methods and intracellular electrodes used to record from motoneurons (see below). Electrodes were advanced into these cells through the dorsal horn (with the dorsal surface oriented upwards), and their identity established by a direct response to light activation of the ChR2 construct (5 – 10 ms light pulse, 447 nm), without a synaptic delay (<1 ms) and continued light response after blocking synaptic transmission.

### Dorsal and ventral root stimulation

Dorsal and ventral roots (DR and VR) were mounted on silver-silver chloride wires above the nASCF of the recording chamber and covered with grease (a 3:1 mixture of petroleum jelly and mineral oil) for monopolar stimulation ^15, 64, 70^. This grease was surrounded by a more viscous synthetic high vacuum grease to prevent oil leaking into the bath flow. Bipolar stimulation was also used at times to reduce the stimulus artifact during recording from ventral roots (detailed below). Roots were stimulated with a constant current stimulator (Isoflex, Israel) with short pulses (0.1 ms). Note that proprioceptive afferents are selectively activated by low intensity DR stimulation (1.1 – 1.5 x afferent volley threshold, T) and cutaneous afferents are additionally activated by higher intensity DR stimulation (2 – 3xT). DRs were dissected to be as long as possible, and the distal end of this root was stimulated, so it was ∼20 mm way from the spinal cord. In this way the DR stimulation site itself (at wire, and threshold for stimulation) could not be affected by axonal depolarizations in the spinal cord, since dorsal root potentials from spinal events (PAD) are only observed very close to the cord (within a few mm, see below), and drop exponentially in size with distance^15^.

### Intracellular recording from sensory axon branches in the dorsal horn

#### Electrode preparation and amplifier

Recording from fine afferent collaterals in the spinal cord without damaging them or disturbing their intracellular milieu required specialized ultra-sharp intracellular electrodes modified from those we developed for motoneuron recording^61^. That is, glass capillary tubes (1.5 mm and 0.86 mm outer and inner diameters, respectively; with filament; 603000 A-M Systems; Sequim, USA) were pulled with a Sutter P-87 puller (Flaming-Brown; Sutter Instrument, Novato, USA) set to make bee-stinger shaped electrodes with a short relatively wide final shaft (∼1 mm) that tapered slowly from 30 to 3 µm over its length, and then abruptly tapered to a final tip over the final 20 µm length. The tip was subsequently bevelled to a < 100 nm hypodermic-shaped point, as verified with electron microscope images (Harvey et al. 2006). This very small tip and wide shaft gave a combination of ease of penetrating axons in dense adult connective tissue, and good current-passing capabilities to both control the potential and fill the axons with neurobiotin. Prior to beveling, electrodes were filled through their tips with 2 M K-acetate mixed with varying proportions of 2 M KCl (to make KCl concentrations ranging of 0, 100, 500, and 1000 mM) or 500 mM KCl in 0.1 Trizma buffer with 5 - 10% neurobiotin (Vector Labs, Birmingame, USA). Electrodes were then beveled from an initial resistance of 40 - 150 MΩ to 30 - 40 MΩ using a rotary beveller (Sutter BV-10). GABAergic chloride-mediated potentials (PAD) and their reversal potentials were the same with different concentrations of KCl, without passing large amounts of negative current, as we have previously detailed^15^, indicating that the ultra-sharp tips impeded passive fluid exchange between the electrode and intracellular milieu, with in particular electrode Cl^-^ not affecting the axon; thus, recordings were mostly made with electrodes with 1 M K-acetate and 1 M KCl, when not filling cells with neurobiotin.

Intracellular recording and current injection were performed with an Axoclamp2B amplifier (Axon Inst. and Molecular Devices, San Jose, USA). Recordings were low pass filtered at 10 kHz and sampled at 30 kHz (Clampex and Clampfit; Molecular Devices, San Jose, USA). Sometimes recordings were made in discontinuous-single-electrode voltage-clamp (gain 0.8 –2.5nA/mV; for Ca PICs) or discontinuous- current-clamp modes (switching rate 7 kHz), as indicated (the latter only when injecting current, for example during recording of input resistance or the voltage dependence of spikes).

#### Axon penetration

Electrodes were advanced into myelinated afferents of the sacrocaudal spinal cord with a stepper motor (Model 2662, Kopf, USA, 10 µm steps at maximal speed, 4 mm/s), usually at the boundary between the dorsal columns and dorsal horn gray matter, where axons bundle together densely, as they branch and descend to the ventral horn (Extended Data Fig 4A). Extracellular tissue (especially myelin in the white matter) often impeded and blocked the electrode tip following a forward step, as determined by an increase in resistance to small current pulses passed from the tip of the electrode (20 ms, -0.3 nA, 1 Hz), and this was cleared with a brief high frequency current (from capacitance overcompensation buzz) and moving backwards slowly, the latter which helped prevent tissue dimpling. Prior to penetrating afferents, we recorded the extracellular (EC) afferent volley following dorsal root (DR) stimulation (0.1 ms pulses, 3xT, T: afferent volley threshold, where T = ∼3 uA, repeated at 1 Hz), to determine the minimum latency and threshold of afferents entering the spinal cord. The group Ia afferent volley occurs first with a latency of 0.5 - 1.0 ms, depending on the root length (which were kept as long as possible, 10 - 20 mm), corresponding to a conduction velocity of about 16 - 24 m/s, as previously described for in vitro conduction at 23 C ^15, 71^. When a forward step penetrated an axon, small slow movements were made to stabilize the recordings. Penetrations were usually in the myelinated portion of the axon between nodes, rather than at nodes, because the chance of penetrating a node is low since they only make up a small fraction of the total axon length (Fig. 1). The spikes from the two nodes adjacent to the electrode were readily detected separately when testing for the spike threshold with current injection pulses (20 ms; rheobase test), because just at threshold the current sometimes evoked a spike from just one node and not the other, which usually halved the total spike height, consistent with the penetration being about halfway between the two nodes separated by about a space constant distance.

#### Proprioceptive afferent identification

Upon penetration, afferents were identified with direct orthodromic spikes evoked from DR stimulation. We focused on the lowest threshold proprioceptive group Ia afferents, identified by their direct response to DR stimulation, very low threshold (< 1.5 x T, T: afferent volley threshold), short latency (group Ia latency, coincident with onset of afferent volley), and antidromic response to ventral horn afferent terminal microstimulation (∼ 10 µA stimulation via tungsten microelectrode to activate Ia afferent terminals; tested in some afferents, detailed below)^15^. Post hoc these were confirmed to be large proprioceptive Ia afferents by their unique extensive terminal branching around motoneurons, unlike large cutaneous Aβ afferents that do not project to the ventral horn. Clean axon penetrations without injury occurred abruptly with a sharp pop detected on speakers attached to the recorded signal, the membrane potential settling rapidly to near – 70 mV, and > 70 mV spikes usually readily evoked by DR stimulation or brief current injection pulses (1 – 3 nA, 20 ms, 1 Hz). Sensory axons also had a characteristic >100 ms long depolarization following stimulation of a dorsal root (primary afferent depolarization, PAD, at 4 - 5 ms latency, detailed below) and short spike afterhyperpolarization (AHP ∼ 10 ms), which further distinguished them from other axons or neurons. Injured axons had higher resting potentials (> - 60 mV), poor spikes (< 60 mV) and low resistance (to current pulse; R_m_ < 10 MΩ), and were discarded.

#### Quantification of spike conduction failure in the dorsal horn: failure potentials (FPs)

Sometimes healthy intracellular penetrations were made into a sensory axon branch (e.g. < -60 mV rest, large PAD), but dorsal root stimulation did not evoke a full spike, even though a full > 60 mV spike could be readily evoked by intracellular current injection. Instead, DR stimulation evoked a partial spike at the latency and threshold of group Ia afferents, indicating that this was a branch of a Ia afferent that failed to fully conduct spikes to the electrode, with only the passively attenuated spike from the last node to spike prior to conduction failure recorded at the electrode (failure potential, FP; also referred to as electronic residue by Luscher^72^). The size of the FP reflected how far away the spike failure occurred, with spatial attenuation corresponding to a space constant of about 90 µm (see Results), and so FPs became exponentially smaller with distance from failure and undetectable when many mm away (nodes separated by about 50 µm). Occasionally axons were penetrated with undetectable DR evoked spikes or FPs, but otherwise they had characteristics of a Ia afferent (PAD, R_m_ similar). These were likely afferents with FPs too distal to detect, but were usually excluded from the main analysis to avoid ambiguity, though this underestimates the incidence of failure. However, some of these axons exhibited short latency, low threshold DR spikes when depolarized by a prior DR stimulation (PAD) of an adjacent DR, in which case they were unequivocally Ia afferents and included in the analysis (Fig. 4f).

Both during extracellular and intracellular recording the group Ia afferent volley (small negative field) was observed as the first event after DR stimulation (the latter subthreshold to a spike), though this was usually small in relation to intracellular events and ignored. However, this was sometimes removed from the intracellular record by subtracting the extracellular potential recorded just outside the same axon to determine the actual transmembrane potential ^15^. This was necessary to see the very smallest FPs following DR stimulation in some afferents, as the negative volley from other nearby afferents obscured the FPs.

After quantifying the axons spikes and conduction failures (FPs) under resting conditions, we then examined the changes in spike conduction with changes in membrane potential induced by either directly injecting current into axons or inducing GABA-mediated changes in membrane potential by pharmacological methods, optogenetic methods (activating ChR2 on GABA_axo_ neurons to induce PAD) or more naturally evoking PAD with a DR stimulation.

#### Neurobiotin filling of axons

Some of the proprioceptive afferents that we recorded intracellularly were subsequently filled with neurobiotin by passing a very large positive 2 - 4 nA current with 90% duty cycle (900 ms on, 100 ms off) for 10 - 20 min. The identity of group Ia proprioceptive afferents were then confirmed anatomically by their unique extensive innervation of motoneurons^15^. Prior to penetrating and filling axons with neurobiotin filled electrodes, a small negative holding current was maintained on the electrodes to avoid spilling neurobiotin outside axons.

### Quantification of spike conduction failure in the ventral horn

#### Wall’s method

To measure whether spikes fail during propagation to their fine terminals in the ventral horn we examined whether failed axon segments were relatively less refractory to activation after spike conduction failure, using a double pulse method adapted from Wall^30, 73^. The essence of the method is that after DR activation all nodes that generate spikes become relatively refractory for a few ms, whereas nodes that fail to spike are not refractory to activation. Thus, a microelectrode placed near these failing nodes more readily activates them if they fail rather than generate spikes with DR stimulation and orthodromic conduction. For this we placed a tungston microelectrode (12 MΩ, #575400, A-M Systems, Sequim, USA) in the ventral horn near the axons terminals on motoneurons, to activate the branches/nodes of the axon projecting to the motoneuron that may have failed (VH stimulation).

Spikes from VH or DR stimulation were recorded intracellularly in a proprioceptive Ia axon penetrated in the dorsal columns directly above the VH stimulation site or in an adjacent segment, with two combinations of double axon stimulations. First, we applied two rapidly repeated VH stimuli (VH doublet; two 0.1 ms pulses) at a ∼4 ms interval to make the axon relatively refractory to stimulation and determine both the threshold current to activate the first spike (*T_VH1_*, with VH1 stimulation) and the higher threshold current to overcome this the inactivation and generate a second spike (*T_VH2_*, with VH2 stimulation). Second, we repeated this double spike activation, but with the first activation from a supra- threshold DR stimulation (at 1.5x DR threshold) and the second from a VH stimulation at the *T_VH2_* intensity from B (DR-VH pair). In this case the VH stimulation readily activates the axon spike if the orthodromic DR evoked spike does not propagate to the ventral horn, leaving the silent portion of the axon non refractory. Accordingly, we also determined the threshold current to activate the VH after the DH in this arrangement (termed *T_DR,VH_*), which was lower than T_VH2_. For comparison to the spike inactivation with VH doublets, we adjusted the DR-VH pair timing slightly so that the pairs of spikes (or expected spikes, at vertical lines) are separated by the same interval (∼ 4 ms) when they reach the recording site, to compensate for DR conduction delays. The putative spike failure with DR stimulation happens at a node somewhere between the recording site and the VH, because we only studied axons that securely conducted single DR pulses to the recording site, and thus failure was not directly visible.

We quantified the spike failure based on the following considerations: If the DR-evoked spike entirely fails to propagate to the VH, then the threshold for subsequently activating the ventral horn (*T_DR,VH_*) should be the same as the threshold without any prior activation (*T_VH1_ = T_DR,VH_*), whereas if it does not fail, then the threshold for activating the ventral horn should be the same as with a VH doublet (*T_VH2_ = T_DR,VH_*). In between these two extreme scenarios, the DR evoked spike may only partially fail to propagate spikes to the ventral horn (by only some of its branches failing or conducting only partially to the VH); in this case *T_DR,VH_* should be between *T_VH1_* and *T_VH2_*, with the difference *T_VH2_ -T_VH1_* representing the range of possible thresholds between full failure and full conduction. Thus, overall the failure was quantified as: *Conduction failure = (T_VH2_ - T_DR,VH_) / (T_VH2_ - T_VH1_) x 100%,* which is 100% at full failure and 0% with no failure. This estimate is predicated on the assumption that the failed spikes are only relatively refractory to conduction and increased stimulation can overcome this failure, which is reasonable for the interspike intervals we used, and means that the computed % failure reflects the number of nodes that failed to spike, with more dorsal branch point failures giving more failed nodes. On the other hand, we used interspike intervals that were short enough for the DR stimulation not to evoke PAD that affected the subsequent spike threshold (∼ 4 ms), in contrast to the longer intervals where PAD can help DR doublet firing (DR-DR in Extended Data Fig. 7, ∼ 5 - 10 ms).

#### Extracellular recording from sensory axon terminals

To directly record spike conduction in proprioceptive afferent terminal branches in the VH we used our intracellular glass pipette electrode (∼30 MΩ) positioned just outside these axons (extracellular, EC), to avoid penetration injury in these fine axon branches. The DR was stimulated near threshold for spikes (1.1xT, T: afferent volley threshold) to evoke the EC response in a few single axons near the electrode, and many trials were averaged to remove noise from these small signals (20 – 50 trials at 3 s intervals). The EC field was multiphasic as previously described for other axons^74–76^, with a small initial positive field resulting from passively conducted axial current from sodium spikes at distant nodes (closer to the DR; outward current at electrode), some of which fail to propagate spikes to the VH recording site, making this field a measure of conduction failure^74, 76^. Following this, a larger negative field arises, resulting from spikes arising at nodes near the electrode (inward current), making this negative field a measure of secure conduction. A relatively large stimulus artifact is present prior to these fields, due to the small size of the EC fields themselves, and we truncated this.

We conducted three control experiments to confirm the relation of these EC fields to spike conduction. First, in the dorsal horn where we can readily intracellularly record from large proprioceptive axon branches, we compared intracellular (IC) recordings from axons to EC recordings just outside the same axon, to confirm that the DR evoked spike (IC) arrives at about the time of the negative EC field. Second, we locally applied TTX to the DR near the recording site (10 µl bolus of 100 µM TTX over DR) which eliminated the negative field and left only the initial positive field, confirming that the positive field is from distal nodes upstream of the TTX block, and generated by passive axial current conduction. This is important, since some investigators have argued on theoretical grounds that the positive field can instead result from the closed-end electrical properties of axons at their terminals^77^, rather than spike failure, though others have refuted this^76^. Finally, we improved nodal spike conduction by reducing the divalent cations Mg^++^ and Ca^++^ in the bath medium, since divalent cations normally cause a gating or guarding action on the sodium channel, the latter by one charge binding to the membrane and the other raising the local extracellular positive charge, and overall raising the local voltage drop across the channel and its spike threshold^78^. This decreased the failure-related initial positive field and increased the main EC negative field, indicating improved conduction, and again confirming the use of these fields as measures of conduction, similar to previous conclusions for the motor endplate^74^ and mathematical consideration of axon cable properties ^79^.

To quantify the EC fields we estimated the overall conduction to the recording site as: *Conduction Index* = *nf / (nf + pf) x 100%*, where pf and nf are the positive and negative EC field amplitudes. This conduction index approaches 100% for full conduction (pf ∼=0) and 0% for no conduction (*nf =* 0). The absolute EC field potential amplitudes are highly variable between different recordings sites, and thus are difficult to quantify across animals and sites, whereas this ratio of field amplitudes (nf / (nf + pf)) eliminates the variability, and can effectively be viewed as a normalization of the negative field (nf) by the total field peak-to-peak size (nf + pf).

### Intracellular recording from motoneurons

The same intracellular glass electrode, stepper motor and amplifier used for recording sensory axons were used for intracellular recording from motoneurons, except that the electrodes were bevelled to a lower resistance (30 MΩ). The electrode was advanced into motoneurons with fast 2 µm steps and brief high frequency currents (capacitance overcompensation) guided by audio feedback from a speaker. After penetration, motoneuron identification was made with antidromic ventral root stimulation, and noting ventral horn location, input resistance and time constant (> 6 ms for motoneurons)^62^. The monosynaptic excitatory postsynaptic potentials (EPSPs) and associated currents (EPSCs) were measured in motoneurons following stimulation of dorsal roots (at 1.1- 1.5 xT, 0.1 ms, 3 – 10 s trial intervals). These were identified as monosynaptic by their rapid onset (first component), lack of variability in latency (< 1 ms jitter), persistence at high rates (10 Hz) and appearance in isolation at the threshold for DR stimulation (< 1.1xT; T, Threshold for EPSP, which also equals afferent volley threshold), unlike polysynaptic EPSPs which varying in latency, disappear at high rates, and mostly need stronger DR stimulation to activate.

### Dorsal and ventral root grease gap recording

In addition to recording directly from single proprioceptive axons and motoneurons, we employed a grease gap method to record the composite intracellular response of many sensory axons or motoneurons by recording from dorsal and ventral roots, respectively, as previously detailed for similar sucrose and grease gap methods, where a high impedance seal on the axon reduces extracellular currents, allowing the recording to reflect intracellular potentials^15, 79–81^. We mounted the freshly cut roots onto silver-silver chloride wires just above the bath, and covered them in grease over about a 2 mm length, as detailed above for monopolar recordings. Return and ground wires were in the bath and likewise made of silver-silver chloride. Specifically for sensory axons, we recorded from the central ends of dorsal roots cut within about 2 - 4 mm of their entry into the spinal cord, to give the compound potential from all afferents in the root (dorsal roots potential, DRP), which has previously been shown to correspond to PAD, though it is attenuated compared to the intracellular recordings of PAD^15^. The signal attenuation has two reasons. First the voltage PAD is attenuated along the length of nerve in the bath, as detailed in the next paragraph. Second, the grease does not completely remove the extracellular fluid around the nerve, even though we deliberately allowed the nerve to dry for a few seconds before greasing, and this causes a conductance that shunts or short circuits the recorded signal, reducing it by about half^74, 81^. For optogenetic experiments we additionally added silicon carbide powder (9 % wt, Tech-Met, Markham) to the grease to make it opaque to light and minimize light induced artifactual current in the silver-silver chloride recording wire during optogenetic activation of ChR2 (detailed below). Likewise, we covered our bath ground and recording return wires with a plastic shield to prevent stray light artifacts. The dorsal root recordings were amplified (2,000 times), high-pass filtered at 0.1 Hz to remove drift, low-pass filtered at 10 kHz, and sampled at 30 kHz (Axoscope 8; Axon Instruments/Molecular Devices, Burlingame, CA).

These grease gap recordings of PAD on sensory afferents reflect only the response of largest diameter axons in the dorsal root, mainly group I proprioceptive afferents, because of the following considerations. First, the largest axons in peripheral nerves have a nodal spacing of about 1 mm^82, 83^, and length constants λ_S_ are estimated to be similar, at about 1 – 2 times the nodal spacing^84^, Further, in our recordings we were only able to get the grease to within about 2 mm of the spinal cord. Thus, the centrally generated signal (PAD) is attenuated exponentially with distance x along the axon length in the bath (x = 2 mm). This is proportional to exp(– x / λ_S_) (see ^79^), which is 1 / e^2^ = 0.11 for x = 2 λ_S_, as is approximately the case here. This makes a central PAD of about 4 mV appear as a ∼0.4 mV potential on the root recording (DRP, 10 times smaller), as we previously reported^15^. Furthermore, the nodal spacing and λ_S_ decrease linearly with smaller axon diameters^79, 82^, making the voltages recorded on the smaller afferents contribute to much less of the compound root potential (halving the diameter attenuates PAD instead by 1/e^4^ or 0.012, which is 99% attenuation). Finally, unmyelinated sensory axons attenuate voltages over a much shorter distance than myelinated axons, since that membrane resistance (*Rm*) drops markedly without myelin and λ_S_ is proportional to 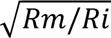 (where *Ri* is axial resistance; Stein 1980). Thus, any centrally generated change in potential in these small axons is unlikely to contribute to the recorded signal 2 mm away.

The composite EPSPs in many motoneurons were likewise recorded from the central cut end of ventral roots mounted in grease (grease gap), which has also previously been shown to yield reliable estimates of the EPSPs, though again attenuated by the distance from the motoneurons^85^. The monosynaptic EPSPs were again identified as monosynaptic by their rapid onset (first component, ∼1 ms after afferent volley arrives in the ventral horn; see below), lack of variability in latency (< 1 ms jitter), persistence at high rates (10 Hz) and appearance in isolation at the threshold (T) for evoking EPSPs with DR stimulation (< 1.1xT, T ∼ afferent volley threshold), unlike polysynaptic reflexes which varying in latency, disappear at high rates, and mostly need stronger DR stimulation to activate.

### Analysis of synaptic responses in sensory axons (PAD) and motoneurons (EPSPs)

When we recorded from sensory axons of an associated dorsal root (directly or via the dorsal roots) stimulation of an adjacent dorsal root (not containing the recorded axon; 0.1 ms, 1 – 3xT; T: threshold to evoke PAD or EPSPs, same as afferent volley threshold) evoked a characteristic large and long depolarization of the afferents, previously demonstrated to be mediated by GABAergic input onto the sensory axons^15^. This depolarization is termed primary afferent depolarization (PAD). PAD occurs at a minimal latency of 4 – 5 ms following the afferent volley, consistent with its minimally trisynaptic origin^13, 15^, making it readily distinguishable from earlier events on the axon. PAD has a fast synaptic component evoked by a single DR stimulation (rising within 30 ms and decaying exponentially over < 100 ms; termed phasic PAD) and a slower longer lasting extrasynaptic component (starting at about 30 ms and lasting many seconds) that is enhanced by repeated DR stimulation (tonic PAD, especially with cutaneous stimulation)^15^. We used this sensory activation of PAD or direct optogenetic activation of PAD to examine the action of GABA on sensory axon spike transmission to motoneurons, usually evoking phasic PAD about 10 – 60 ms prior to spikes or associated EPSPs on motoneurons (during phasic PAD), though we also examined longer lasting effects of tonic PAD evoked by repeated DR stimulation. Sometimes PAD is so large that it directly evokes spikes on the afferents, and these travel out the dorsal root, and thus they have been termed dorsal root reflexes (DRRs)^15, 22^. We usually minimized these DRRs by keeping the DR stimulus that evokes PAD low (1.1 - 3.0 xT), though there were inevitably some DRRs, as they even occur in vivo in cats and humans8^,47, 86^.

When we recorded from motoneurons (directly or via ventral roots) stimulation of proprioceptive afferents in a dorsal root (0.1 ms, 1.1-1.5xT, T: EPSP threshold, which is similar to afferent volley threshold) evoked a monosynaptic EPSP, and associated monosynaptic reflex (MSR, spikes from EPSP). This EPSP is depressed by fast repetition (rate depended depression, RDD) ^87^, and thus to study the EPSP we evoked it at long intervals (10 s, 0.1 Hz rate) where RDD was less. However, even with this slow repetition rate (0.1 Hz), at the start of testing the first EPSP was often not similar to the steady state EPSP after repeated testing. Thus, to avoid RDD we usually ran the 0.1 Hz EPSP testing continuously throughout the experiment, at least until a steady state response was reached (after 10 minutes). We then examined the action of activating (or inhibiting) GABA_axo_ neurons on this steady state EPSP, by introducing light or sensory conditioning that activated these neurons at varying intervals (inter-stimulus intervals, ISIs) prior to each EPSP stimulation (control, GAD2//ChR2 mice and GAD2//Arch3 mice). We averaged the EPSP from ∼10 trials (over 100 s) just before conditioning and then 10 trials during conditioning, and then computed the change in the peak size of the monosynaptic EPSP with conditioning from these averages. After conditioning was completed EPSP testing continued and any residual changes in the EPSP was computed from the 10 trials following conditioning (after- effect). Finally, EPSP testing continued over many minutes after which the original steady state EPSP was established. The background motoneuron potential, membrane resistance (Rm) and time constant just prior to the EPSP was also assessed before and after conditioning to examine whether there were any postsynaptic changes that might contribute to changes in the EPSP with conditioning. Along with the VR recordings, we simultaneously recorded PAD from DRs by similar averaging methods (10 trials of conditioning), to establish the relation of changes in EPSPs with associated sensory axon depolarization PAD.

### Drugs and solutions

Two kinds of artificial cerebrospinal fluid (ACSF) were used in these experiments: a modified ACSF (mACSF) in the dissection chamber prior to recording and a normal ACSF (nACSF) in the recording chamber. The mACSF was composed of (in mM) 118 NaCl, 24 NaHCO3, 1.5 CaCl2, 3 KCl, 5 MgCl2, 1.4 NaH2PO4, 1.3 MgSO4, 25 D-glucose, and 1 kynurenic acid. Normal ACSF was composed of (in mM) 122 NaCl, 24 NaHCO3, 2.5 CaCl2, 3 KCl, 1 MgCl2, and 12 D-glucose. Both types of ACSF were saturated with 95% O2-5% CO2 and maintained at pH 7.4. The drugs sometimes added to the ACSF were APV (NMDA receptor antagonist), CNQX (AMPA antagonist), gabazine (GABA_A_ antagonist), bicuculline (GABA_A_, antagonist), L655708 (α5 GABA_A_, antagonist), CGP55845 (GABA_B_ antagonist; all from Tocris, USA), 5-HT, kynurenic acid (all from Sigma-Aldrich, USA), and TTX (TTX-citrate; Toronto Research Chemicals, Toronto). Drugs were first dissolved as a 10 - 50 mM stock in water or DMSO before final dilution in ACSF. DMSO was necessary for dissolving gabazine, L655708, bicuculline and CGP55845, but was kept at a minimum (final DMSO concentration in ACSF < 0.04%), which by itself had no effect on reflexes or sensory axons in vehicle controls (not shown). L655708 was particularly difficult to dissolve and precipitated easily, especially after it had been exposed a few times to air; so immediately after purchase we dissolved the entire bottle and froze it at -40°C in single use 5 - 20 µl aliquots, and upon use it was first diluted in 100 µl distilled water before dispersing it into ACSF.

### Recording monosynaptic reflexes in awake mice and rats, and PAD activation

#### Window implant over spinal cord

In GAD2//ChR2+ mice and control GAD2//ChR- mice a glass window was implanted over the exposed spinal cord to gain optical access to the sacrocaudal spinal cord, as described previously ^68^. Briefly, mice were given Meloxicam (1 mg/kg, s.c.) and then anesthetized using ketamine hydrochloride (100 mg/kg, i.p.) and xylazine (10 mg/kg, i.p.). Using aseptic technique, a dorsal midline incision was made over the L2 to L5 vertebrae. Approximately 0.1 ml of Xylocaine (1%) was applied to the surgical area and then rinsed. The animals were suspended from a spinal-fork stereotaxic apparatus (Harvard Apparatus) and the muscles between the spinous and transverse processes were resected to expose the L2 to L5 vertebrae. The tips of modified staples were inserted along the lateral edge of the pedicles and below the lateral processes of L2 and L5, and glued in place using cyanoacrylate. A layer of cyanoacrylate was applied to all of the exposed tissue surrounding the exposed vertebrae followed by a layer of dental cement to cover the cyanoacrylate and to form a rigid ring around the exposed vertebrae. A modified paperclip was implanted in the layer of dental cement to serve as a holding point for surgery. A laminectomy was performed at L3 and L4 to expose the spinal cord caudal to the transection site. Approximately 0.1 ml of Xylocaine (1%) was applied directly to the spinal cord for 2 – 3 s, and then rinsed. A line of Kwik-Sil (World-Precision Instruments) was applied to the dura mater surface along the midline of the spinal cord and a glass window was immediately placed over the exposed spinal cord. The window was glued in place along the outer edges using cyanoacrylate followed by a ring of dental cement. Small nuts were mounted onto this ring to later bolt on a backpack to apply the laser light (on the day of experimentation). Saline (1 ml, s.c.) and buprenorphine (0.03 mg/kg, s.c.) was administered post-operatively, and analgesia was maintained with buprenorphine (0.03 mg/kg, s.c.) every 12 hours for two days. Experimentation started 1 week after the window implant when the mouse was fully recovered.

#### Percutaneous EMG wire implant and fibre optic cable attachment

On the day of experimentation, the mouse was briefly sedated with isoflurane (1.5 %) and fine stainless steel wires (AS 613, Cooner Wire, Chatsworth, USA) were percutaneously implanted in the tail for recording EMG and stimulating the caudal tail trunk nerve, as we previously detailed (wires de-insulated by 2 mm at their tip and inserted in the core of 27 gauge needle that was removed after insertion)^62^. A pair of wires separated by 8 mm were inserted at base of the tail for recording EMG the tail muscles, and second pair of wires was inserted nearby for bipolar activation of the caudal trunk nerve to evoke reflexes. A fifth ground wire was implanted between the EMG and stimulation wires. Following this a backpack was bolted into the nuts imbedded in the dental cement ring around the window. This backpack held and aligned a light fibre optic cable that was focused on the centre of the S3 – S4 sacral spinal cord. The Cooner wires were secured to the skin with drops of cyanoacrylate and taped onto the backpack so that the mouse could not chew them. The isoflurane was removed, and the mouse quickly recovered from the anesthesia and was allowed to roam freely around an empty cage during recording, or was sometimes lightly restrained by hand or by a sling. The fibre optic cable was attached to a laser (447 nM, same above) and the Cooner wires attached to the same models of amplifiers and stimulators used for ex vivo monosynaptic testing detailed above.

#### MSR testing

The monosynaptic reflex (MSR) was recorded in the tail EMG at ∼6 ms latency after stimulating the caudal tail trunk nerve at a low intensity that just activated proprioceptive afferents (0.2 ms current pulses, 1.1 xT, T: Threshold to evoke MSR, which is near afferent threshold), usually near the threshold to activate motor axons and an associated M-wave (that arrived earlier). We studied the tail MSR reflex because our ex vivo recordings were made in the corresponding sacral spinal cord of adult mice and rats, which is the only portion of the spinal cord that survives whole ex vivo, due to its small diameter^64^. This reflex was verified to be of monosynaptic latency because it was the first reflex to arrive, had little onset jitter, and had the same latency as the F wave (not shown; the F wave is evoked by a strong stimulation of all motor axons, at 5xT, which causes a direct motoneuron response on the same axons, while the monosynaptic EPSP is blocked by collision at this intensity) ^88^. The MSR also underwent rate dependent depression (RDD) with fast repeated stimulation and so was synaptic and not a direct muscle response (M-wave), which occurred earlier at sufficient intensity to recruit the motor axons (not shown).

#### Conditioning of the MSR by optogenetic activation of GABA_axo_ neurons

As with in vitro EPSP testing, the MSR was tested repeatedly at long 5 – 10 s intervals until a steady state MSR was achieved. Then testing continued but with a conditioning light pulse applied just prior to the MSR stimulation (40 – 120 ms), to examine the effect of PAD evoked during this time frame on sensory transmission to motoneurons. Background EMG just prior to MSR testing was assessed to estimate the postsynaptic activity on the motoneurons. The changes in MSR and background EMG with light were quantified by comparing the average response before and during the light application, computed from the mean rectified EMG at 6 – 11 ms after the nerve stimulation (MSR) and over 20 ms prior to the nerve stimulation (background just prior to the MSR, Bkg). Because awake mice spontaneously varied their EMG, we plotted the relation between the MSR and the background EMG, with as expected a positive linear relation between these two variables^89^, computed by fitting a regression line. In trials with conditioning light applied the same plot of EMG vs background EMG was made and a second regression line computed. The change in the MSR with conditioning at a fixed matched background EMG level was then computed for each mouse by measuring the difference between the regression line responses at the fixed background EMG. This ruled out changes in MSRs being due to postsynaptic changes. Two background levels were assessed: rest (0%) and 30% of maximum EMG, expressed as a percentage of the control pre-conditioning MSR. The change in background EMG with light was computed by comparing the EMG just prior to the light application (over 20 ms prior) to the EMG just prior to the MSR (over 20 ms prior, Bkg), and expressed as a percentage of the maximum EMG.

#### Cutaneous conditioning of the MSR in rats

A similar examination of how PAD affected the MSR was performed in rats with percutaneous tail EMG recording. However, in this case PAD was evoked by a cutaneous conditioning stimulation of the tip of the tail (0.2 ms pulses, 3xT, 40 – 120 ms prior to MSR testing) using an additional pair of fine Cooner wires implanted at the tip of the tail (separated by 8 mm). In rats the MSR latency is later than in mice due to the larger peripheral conduction time, ∼12 ms (as again confirmed by a similar latency to the F wave). This MSR was thus quantified by averaging rectified EMG over a 12 – 20 ms window. Also, to confirm the GABA_A_ receptor involvement in regulating the MSR, the antagonist L655708 was injected systemically (1 mg/kg i.p., dissolved in 50 µl DMSO and diluted in 900 µl saline). Again, the MSR was tested at matched background EMG levels before and after conditioning (or L655708 application) to rule out changes in postsynaptic inhibition.

### Conditioning of the MSRs in humans

#### H-reflex as an estimate of the MSR

Participants were seated in a reclined, supine position on a padded table. The right leg was bent slightly to access the popliteal fossa and padded supports were added to facilitate complete relaxation of all leg muscles. A pair of Ag-AgCl electrodes (Kendall; Chicopee, MA, USA, 3.2 cm by 2.2 cm) was used to record surface EMG from the soleus muscle. The EMG signals were amplified by 1000 and band-pass filtered from 10 to 1000 Hz (Octopus, Bortec Technologies; Calgary, AB, Canada) and then digitized at a rate of 5000 Hz using Axoscope 10 hardware and software (Digidata 1400 Series, Axon Instruments, Union City, CA) ^62^. The tibial nerve was stimulated with an Ag-AgCl electrode (Kendall; Chicopee, MA, USA, 2.2 cm by 2.2 cm) in the popliteal fossa using a constant current stimulator (1 ms rectangular pulse, Digitimer DS7A, Hertfordshire, UK) to evoke an H-reflex in the soleus muscle, an estimate of the MSR ^90^. Stimulation intensity was set to evoke a test (unconditioned) MSR below half maximum. MSRs recorded at rest were evoked every 5 seconds to minimize RDD ^91^ and at least 20 test MSRs were evoked before conditioning to establish a steady baseline because the tibial nerve stimulation itself can presumably also activate spinal GABAergic networks, as in rats. All MSR were recorded at rest, except when the motor unit firing probabilities were measured (see below).

#### Conditioning of the MSR

To condition the soleus MSR by cutaneous stimulation, the cutaneous medial branch of the deep peroneal (cDP) nerve was stimulated on the dorsal surface of the ankle using a bipolar arrangement (Ag-AgCl electrodes, Kendall; Chicopee, MA, USA, 2.2 cm by 2.2 cm), set at 1.0xT, where T is the threshold for cutaneous sensation. A brief burst (3 pulses, 200 Hz for 10 ms) of cDP stimuli was applied before evoking a MSR at various inter-stimulus intervals (ISIs; interval between tibial and cDP nerve stimuli) within the window expected for phasic PAD evoked by cutaneous stimuli, presented in random order at 0, 30, 60, 80, 100, 150 and 200 ms ISIs. Seven conditioned MSR at each ISI were measured consecutively and the average of these MSR (peak-to-peak) was used as an estimate of the conditioned MSR. This was compared to the average MSR without conditioning, computed from the 7 trials just prior to conditioning.

The cDP nerve was also stimulated with a 500 ms long train at 200 Hz to condition the MSR, and examine the effect of tonic PAD evoked by such long trains, as in rats. Following the application of at least 20 test MSRs (every 5 s), a single cDP train was applied 700 ms before the next MSR and following this the MSR continued to be evoked for another 90 to 120 s (time frame of tonic PAD). We also conditioned the soleus MSR with tibialis anterior (TA; antagonist muscle, flexor) tendon vibration (brief burst of 3 cycles of vibration at 200Hz) to preferentially activate Ia afferents, as has been done previously ^90^.

#### Motor unit recording to examine postsynaptic actions of conditioning

Surface electrodes were used to record single motor units in the soleus muscle during low level contractions by placing electrodes on or near the tendon or laterally on the border of the muscle as detailed previously ^92^. Alternatively, single motor unit activity from the soleus muscle was also recorded using a high density surface EMG electrode (OT Bioelettronica, Torino, Italy, Semi-disposable adhesive matrix, 64 electrodes, 5×13, 8 mm inter-electrode distance) with 3 ground straps wrapped around the ankle, above and below the knee. Signals were amplified (150 times), filtered (10 to 900 Hz) and digitized (16 bit at 5120 Hz) using the Quattrocento Bioelectrical signal amplifier and OTBioLab+ v.1.2.3.0 software (OT Bioelettronica, Torino, Italy). The EMG signal was decomposed into single motor units using custom MatLab software as per ^93^. Intramuscular EMG was used to record MUs in one participant as detailed previously ^94^ to verify single motor unit identification from surface EMG.

To determine if there were any postsynaptic effects from the conditioning stimulation on the motoneurons activated during the MSR, we examined whether the cDP nerve stimulation produced any changes in the tonic firing rate of single motor units, which gives a more accurate estimate of membrane potential changes in motoneurons compared to compound EMG. Single motor units were activated in the soleus muscle by the participant holding a small voluntary contraction of around 5% of maximum. Both auditory and visual feedback were used to keep the firing rates of the units steady while the conditioning cutaneous was applied every 3 to 5 seconds. The instantaneous firing frequency profiles from many stimulation trials were superimposed and time-locked to the onset of the conditioning stimulation to produce a peri-stimulus frequencygram (PSF, dots in Extended Data Fig 13b*iii*), as previously detailed ^94, 95^. A mean firing profile resulting from the conditioning stimulation (PSF) was produced by averaging the frequency values in 20 ms bins across time post conditioning (thick lines in Extended Data Fig 13b*iii* and c*iii*). To quantify if the conditioning stimulation changed the mean firing rate of the tonically firing motor units, the % change in the mean PSF rate was computed at the time when the H reflex was tested (vertical line in Extended Data Fig 13b*ii-iii*).

#### Unitary EPSP estimates from PSF

To more directly examine if the facilitation in MSR resulted from changes in transmission in Ia afferents after cutaneous afferent conditioning, we measured changes in the firing probability of single motor units (MUs) during the brief MSR time-course (typically 30 to 45 ms post tibial nerve stimulation) with and without cDP nerve conditioning. Soleus MSRs were as usual evoked by stimulating the tibial nerve, but while the participant held a small voluntary plantarflexion to activate tonic firing of a few single motor units. The size of the MSR was set to just above reflex threshold (when the M-wave was < 5% of maximum) so that single motor units at the time of the MSR could be distinguished from the compound potential from many units that make up the MSR ^96^. For a given trial run, test MSRs were evoked every 3-5 s for the first 100 s and then MSR testing continued for a further 100s, but with a cDP-conditioning train (50 ms, 200 Hz) applied 500 ms prior to each MSR testing stimulation. These repeated high frequency trains evoke a tonic PAD in rats that facilitates sensory conduction. A 500 ms ISI was used to ensure the firing rate of the motor unit returned to baseline before the MSR was evoked, and this is also outside of the range of phasic PAD. Approximately 40-50 usable test and conditioned firing rate profiles were produced for a single session where the motor units had a steady discharge rate before the cDP nerve stimulation. Sessions were repeated 3-6 times to obtain a sufficient number of frequency points to construct the PSF (∼ 200 trials).

To estimate the EPSP profile and prior background motoneuron activity, motor unit (MU) firing was again used to construct a PSF, as detailed above, but this time locked to the tibial nerve stimulation used to evoke the MSR, so that we could estimate the motoneuron behaviour during the MSR (EPSP). When more than one MU was visible in the recordings firing from these units (usually 2 – 3) were combined into a single PSF. Overall this gave about of 100 – 600 MU MSR test sweeps to generate each PSF. Firing frequency values were initially averaged in consecutive 20 ms bins to produce a mean PSF profile over time before the tibial nerve stimulation, for both unconditioned and conditioned MSR reflex trials. From this, the mean background firing rate within the 100 ms window immediately preceding the tibial stimulation was compared between the test and conditioned MSR trials to determine if the conditioning cDP nerve stimulation produced a change in firing rate, and thus post-synaptic effect, just before the conditioned MSR was evoked. Next, as an estimate of EPSP size, the mean firing rate during the MSR window was also measured, but computed with smaller PSF bins of 0.5 ms during the MSR. Finally, for each PSF generated with or without conditioning, the probability that a motor unit discharged during the MSR window (30 to 45 ms after the TN stimulation) was measured as the number of discharges during the time of the MSR window divided by the total number of tibial nerve test stimuli.

### Temperature, latency and PAD considerations

Large proprioceptive group Ia sensory afferents conduct in the peripheral tail nerve with a velocity of about 33 m/s (33 mm/ms) in mice ^97^. Motor axons are similar, though slightly slower (30 m/s)^98^. Thus, in the awake mouse stimulation of Ia afferents in the mouse tail evokes spikes that take ∼ 2 ms to conduct to the motoneurons in the spinal cord ∼70 mm away. Following ∼1 ms synaptic and spike initiation delay in motoneurons, spikes in the motor axons take a further ∼2 ms to reach the muscles, after which the EMG is generated with a further 1 ms synaptic and spike initiation delay at the motor endplate to produce EMG. All told this gives a monosynaptic reflex latency of ∼6 ms. The motor unit potentials within the EMG signal have a duration of about 3 – 5 ms, and thus we averaged rectified EMG over 6 – 11 ms to quantify the MSR. We have shown that similar considerations hold for the rat where tail nerve conduction velocities are similar, except the distance from the tail stimulation to the spinal cord is larger (150 mm), yielding a peripheral nerve conduction delay of ∼10 ms and total MSR delay of ∼12 ms ^99^. In humans the MSR latency is dominated by the nerve conduction latency (50 – 60 m/s) over a large distance (∼800 mm), yielding MSR latencies of ∼30 ms.

In our ex vivo whole adult spinal cord preparation the bath temperature was varied between 23 and 32°C. All data displayed is from 23 – 24°C, though we confirmed the main results (facilitation of sensory axon transmission to motoneuron by PAD) at 32°C. The Q10 for peripheral nerve conduction (ratio of conduction velocities with a 10 °C temperature rise) is about 1.3 ^100^, yielding a conduction in dorsal roots of about 20 m/s at 23 – 24 °C, as we directly confirmed (not shown). Thus, when the DR is stimulated 20 mm from the cord the latency of spike arrival at the cord should be about 1 ms, which is consistent with the time of arrival of afferent volleys that were seen in the intracellular and extracellular recordings from sensory axons (e.g. Figs. 2b and 4e).

When we found that PAD evoked in sensory axons can prevent failure of spikes to propagate in the cord after DR stimulation, we worried that PAD somehow influenced the initiation of the spike by the dorsal root stimulation at the silver wire. However, we ruled this out by stimulating dorsal roots as far away from the spinal cord as possible (20 mm), where PAD has no effect, due to the exponential attenuation of its dorsal root potential with distance (see above), and found that PAD still facilitated sensory axon spike transmission to motoneurons. The added advantage of these long roots is that there is a clean 1 ms separation between the stimulus artifact and the afferent volley arriving at the spinal cord, allowing us to quantify small FPs and afferent volleys that are otherwise obscured by the artifact.

We did not consistently use high temperature ex vivo baths (32°C) because the VR and DR responses to activation of DRs or PAD neurons are irreversibly reduced by prolonged periods at these temperatures, suggesting that the increased metabolic load and insufficient oxygen penetration deep in the tissue damages the cord at these temperatures. Importantly, others have reported that in sensory axons PAD-evoked spikes (DRRs) are eliminated in a warm bath and argued that this means they are not present in vivo, and not able to evoke a motoneuron response ^6^, despite evidence to the contrary ^8, 47^. However, we find that PAD itself is reduced in a warm bath by the above irreversible damage, and it is thus not big enough to evoke spikes in sensory axons; thus, this does not tell us whether these spikes should be present or not in vivo. Actually, in vivo we sometimes observed that with optogenetic activation of GABA_axo_ neurons and associated PAD there was a direct excitation of the motoneurons (seen in the EMG) at the latency expected for PAD evoked spikes (not shown). However, this was also at the latency of the postsynaptic inhibition produced by this same optogenetic stimulation, which often masked the excitation (Fig. 6). In retrospect, examining the GABA_axo_ evoked motoneuron responses during optogenetic-evoked PAD (Fink et al.)^6, 16^, or sensory-evoked PAD^16, 17^, there is either outright excitation or an excitation riding on the postsynaptic IPSPs resulting from the activation of there GABA_axo_ neurons. This is consistent with the PAD-evoked spike activating the monosynaptic pathway, which inhibits subsequently tested monosynaptic responses by post activation depression (see Discussion).

The latency of a single synapse in our ex vivo preparation at 23 – 24°C was estimated from the difference between the time arrival of the sensory afferent volley at the motoneurons (terminal potential seen in intracellular and extracellular recordings) and the onset of the monosynaptic EPSP in motoneurons. This was consistently 1 – 1.2 ms (Fig. 5b and e). This is consistent with a Q10 of about 1.8 – 2.4 for synaptic transmission latency ^101, 102^, and 0.4 ms monsynaptic latency at body temperature ^103, 104^. Based on these considerations we confirm that the PAD evoked in sensory axons is monosynaptically produced by optogenetic activation of GABA_axo_ neurons with light, since it follows ∼1 ms after the first spike evoked in GABAaxo neurons by light (Fig. 3a). This first spike in GABA_axo_ neurons itself takes 1 – 2 ms to arise and so the overall latency from light activation to PAD production can be 2 - 3 ms (Fig. 3f), as seen for IPSCs at this temperature in other preparations ^105^. With DRs stimulation PAD arises with a minimally 4 – 5 ms latency, which is consistent with a trisynaptic activation of the sensory axon, after taking into account time for spikes to arise in the interneurons involved (Fig. 4a,e).

### Viral labelling of sensory afferents

Large diameter peripheral afferents were labelled by viral injections, as previously detailed^43, 106^, providing an additional method of examining central afferent projections in the spinal cord. Adeno-associated viral vectors (AAVs) with the transgene encoding the cytoplasmic fluorophore tdTom under the CAG promoter were injected IP into anesthetized P1-2 mice (AAV9-CAG-tdTom, 5.9 x 10^12^ vg/ml; 2-4 µl per injection; UNC Vector Core). To improve transduction efficiencies, viral vectors were incubated with LAH4 peptide (200 µM) at 37 °C for ∼45 min immediately prior to injection. Mice were perfused for immunolabelling > 60 days post injection (adult mice). This injection method yields a sparce Golgi-like labelling of about 5% of afferents in each spinal segment, without central labelling of other neurons (with the exception of one or two motoneurons labelled per spinal segment), allowing afferents to be traced to the motoneurons for morphological identification as Ia afferents.

### Immunohistochemisty

#### Tissue fixation and sectioning

After sensory axons were injected with neurobiotin ex vivo in mouse and rat sacrocaudal spinal cords, the cords were left in the recording chamber in oxygenated nACSF for an additional 4 – 6 hr to allow time for diffusion of the neurobiotin throughout the axon. Then the spinal cord was immersed in 4% paraformaldehyde (PFA; in phosphate buffer) for 20-22 hours at 4°C, cryoprotected in 30% sucrose in phosphate buffer for 24-48 hours. Alternatively, afferents were labelled genetically in VGLUT1^Cre/+^; R26^lsl-tdTom^ mice or by a AAV9-CAG-tdTom viral injection, which were euthanized with Euthanyl (BimedaMTC; 700 mg/kg) and perfused intracardially with 10 ml of saline for 3 – 4 min, followed by 40 ml of 4% paraformaldehyde (PFA; in 0.1 M phosphate buffer at room temperature), over 15 min (Gabra5-KO mice also fixed similarly). Then spinal cords of these mice were post-fixed in PFA for 1 hr at 4°C, and then cryoprotected in 30% sucrose in phosphate buffer (∼48 hrs). Following cryoprotection all cords were embedded in OCT (Sakura Finetek, Torrance, CA, USA), frozen at -60C with 2-methylbutane, cut on a cryostat NX70 (Fisher Scientific) in sagittal or transverse 25 µm sections, and mounted on slides. Slides were frozen until further use.

#### Immunolabelling

The tissue sections on slides were first rinsed with phosphate buffered saline (PBS, 100 mM, 10 min) and then again with PBS containing 0.3% Triton X-100 (PBS-TX, 10 min rinses used for all PBS-TX rinses). For the sodium channel antibody, we additionally performed antigen retrieval by incubating slides three times for 10 min each with a solution of 0.2% sodium borohydride (NaBH4, Fisher, S678-10) in PB, followed by a PBS rinse (4x 5 min), because this antibody is sensitive to over-fixation. We verified that this sodium channel antibody labels axon nodes just as well in our tissue treated with the antigen retrieval, compared to in control tissue that was only lightly fixed (PFA perfusion, followed by no postfixation, not shown). Next, for all tissue, nonspecific binding was blocked with a 1 h incubation in PBS-TX with 10% normal goat serum (NGS; S-1000, Vector Laboratories, Burlingame, USA) or normal donkey serum (NDS; ab7475, Abcam, Cambridge, UK). Sections were then incubated for at least 20 hours at room temperature with a combination of the following primary antibodies in PBS-TX with 2% NGS or NDS: rabbit anti-α_5_ GABA_A_ receptor subunit (1:200; TA338505, OriGene Tech., Rockville, USA; same antibody as SAB2100878, Sigma-Aldrich, St. Louis, USA; verified with Western blot and IHC^107, 108^, and knockout detailed below), rabbit anti-α1 GABA_A_ receptor subunit (1:300; 06-868, Sigma-Aldrich, St. Louis, USA; verified by Western blot, IHC, and α1 GABA_A_ knockout^109, 110^), guinea pig anti-α2 GABA_A_ receptor subunit (1:500; 224 104, Synaptic Systems, Goettingen, Germany; verified with Western blot, IHC, lack of labelling with loss of GABRA2 quantified with RT-qPCR^111^, and labelling in HEK239 cells transfected with GABRA2 cDNA^112^), chicken anti-γ2 GABA_A_ receptor subunit (1:500; 224 006, Synaptic Systems, Goettingen, Germany; verified with Western blot, IHC and receptor colocalization with gephrine^113^), rabbit anti- GABA_B1_ receptor subunit (1:500; 322 102, Synaptic Systems, Goettingen, Germany), mouse anti-Neurofilament 200 (NF200) (1:2000; N0142, Sigma-Aldrich, St. Louis, USA), guinea pig anti-Neurofilament M (NFM, 1:500; 171 204, Synaptic Systems), guinea pig anti-VGLUT1 (1:1000;AB5905, Sigma-Aldrich, St. Louis, USA), rabbit anti-Caspr (1:500; ab34151, Abcam, Cambridge, UK), mouse anti-Caspr (1:500; K65/35, NeuroMab, Davis, USA), chicken anti-Myelin Basic Protein (MBP) (1:200; ab106583, Abcam, Cambridge, UK), guinea pig anti-GAD2/GAD65 (1:500; 198 104; Synaptic Systems); chicken anti-VGAT (1:500; 131 006, Synaptic Systems, Goettingen, Germany), rabbit anti-VGAT (1:500; AB5062P, Sigma-Aldrich, St. Louis, USA), rabbit anti-EYFP (1:500; orb256069, Biorbyt, Riverside, UK), goat anti-RFP (1:500; orb334992, Biorbyt, Riverside, UK), rabbit anti-RFP (1:500; PM005, MBL International, Woburn, USA), rabbit anti-GFP (1:500, A11122, ThermoFisher Scientific, Waltham, USA), and mouse anti-Pan Sodium Channel (1:500; S8809, Sigma-Aldrich, St. Louis, USA). The latter is a pan-sodium antibody, labelling an intracellular peptide sequence common to all known vertebrate sodium channels. Genetically expressed EYFP, tdTom (RFP) and GFP were amplified with the above antibodies, rather than rely on the endogenous fluorescence. When anti-mouse antibodies were applied in mice tissue, the M.O.M (Mouse on Mouse) immunodetection kit was used (M.O.M; BMK-2201, Vector Laboratories, Burlingame, USA) prior to applying antibodies. This process included 1h incubation with a mouse Ig blocking reagent. Primary and secondary antibody solutions were diluted in a specific M.O.M diluent.

The following day, tissue was rinsed with PBS-TX (3x 10 min) and incubated with fluorescent secondary antibodies. The secondary antibodies used included: goat anti-rabbit Alexa Fluor 555 (1:200; A32732, ThermoFisher Scientific, Waltham, USA), goat anti-rabbit Alexa Fluor 647 (1:500, ab150079, Abcam, Cambridge, UK), goat ant-rabbit Pacific orange (1:500; P31584, ThermoFisher Scientific, Waltham, USA), goat anti-mouse Alexa Fluor 647 (1:500; A21235, ThermoFisher Scientific, Waltham, USA), goat anti-mouse Alexa Fluor 488 (1:500; A11001, ThermoFisher Scientific, Waltham, USA), goat anti-mouse Alexa Fluor 555 (1:500; A28180, ThermoFisher Scientific, Waltham, USA), goat anti-guinea pig Alexa Fluor 647 (1:500; A21450, ThermoFisher Scientific, Waltham, USA), goat anti-chicken Alexa Fluor 405 (1:200; ab175674, Abcam, Cambridge, UK), goat anti-chicken Alexa Fluor 647 (1:500; A21449, ThermoFisher Scientific, Waltham, USA), donkey anti-goat Alexa Fluor 555 (1:500; ab150130, Abcam, Cambridge, UK), donkey anti-rabbit Alexa Fluor 488 (1:500; A21206, ThermoFisher Scientific, Waltham, USA), Streptavidin-conjugated Alexa Fluor 488 (1:200; 016-540-084, Jackson immunoResearch, West Grove, USA) or Streptavidin-conjugated Cyanine Cy5 (1:200; 016-170-084, Jackson immunoResearch, West Grove, USA) in PBS-TX with 2% NGS or NDS, applied on slides for 2 h at room temperature. The latter streptavidin antibodies were used to label neurobiotin filled afferents. After rinsing with PBS-TX (2 times x 10 min/each) and PBS (2 times x 10 min/each), the slides were covered with Fluoromount-G (00-4958-02, ThermoFisher Scientific, Waltham, USA) and coverslips (#1.5, 0.175 mm, 12-544-E; Fisher Scientific, Pittsburg, USA).

Standard negative controls in which the primary antibody was either 1) omitted or 2) blocked with its antigen (quenching) were used to confirm the selectivity of the antibody staining, and no specific staining was observed in these controls. Previous tests detailed by the manufactures further demonstrate the antibody specificity, including quenching, immunoblots (Western blots), co-immunoprecipitation, and/or receptor knockout. Most antibodies had been previously tested with quenching for selectivity, as detailed in the manufacture’s literature and other publications ^15^, but we verified this for the GABA receptors with quenching. For antibody quenching, the peptides used to generate the antibodies, including anti-α5 GABA_A_ receptor subunit (AAP34984, Aviva Systems Biology, San Diego, USA), anti-α1 GABA_A_ receptor subunit (224-2P, Synaptic Systems, Goettingen, Germany) and anti-γ2 GABA_A_ receptor subunit (224-1P, Synaptic Systems, Goettingen, Germany), were mixed with the antibodies at a 10:1 ratio and incubated for 20 h and 4°C. This mixture was then used instead of the antibody in the above staining procedure. Control receptor knockout experiments were also preformed on for the anti-α5 GABA_A_ antibody, with this antibody producing no receptor labelling in brain tissue from α5 GABA_A_ knockout mice (Gabra5 KO mice; Extended Data Fig. 14).

### Confocal and epifluorescence microscopy

Image acquisition was performed by confocal (Leica TCS SP8 Confocal System) and epifluorescence (Leica DM 6000 B) microscopy for high magnification 3D reconstruction and low magnification imaging, respectively. All the confocal images were taken with a 63x (1.4 NA) oil immersion objective lens and 0.1 µm optical sections that were collected into a z-stack over 10–20 µm. Excitation and recording wavelengths were set to optimize the selectivity of imaging the fluorescent secondary antibodies. The same parameters of laser intensity, gain and pinhole size was used to take pictures for each animal, including the negative controls. Complete sagittal sections were imaged with an epifluorescence 10x objective lens using the Tilescan option in Leica Application Suite X software (Leica Microsystems CMS GmbH, Germany). Sequential low power images were used to reconstruct the afferent extent over the whole spinal cord, using CorelDraw (Ottawa, Canada), and to identify locations where confocal images were taken.

### 3D reconstruction of afferents and localization of GABA receptors

The fluorescently labelled afferents (neurobiotin, tdTom), GABA receptors, VGLUT1, VGAT, NF200, Caspr, MBP and sodium channels were analyzed by 3D confocal reconstruction software in the Leica Application Suite X (Leica Microsystems CMS GmbH) ^15^. To be very conservative in avoiding non-specific antibody staining, a threshold was set for each fluorescence signal at a minimal level where no background staining was observed in control tissue with the primary antibody omitted, less 10%. Signals above this threshold were rendered in 3D for each antibody. Any GABA receptor, Caspr or Na_V_ expression within the volume of the neurobiotin filled axon (binary mask set by threshold) was labelled in 3D reconstructions (yellow, pink and white respectively in Fig. 1). Receptor density within the axon membrane surface area was quantified using the same Leica software. Receptors are usually cycled in and out of the membrane^62^ and so receptors within the axon cytoplasm provide additional evidence of the presence of axonal GABA receptors, distinct from receptors that may be in the postsynaptic contacts of the afferents. Thus, we also computed the receptor densities in the axon volume and found qualitatively the same distribution (at nodes and terminals) as with surface density calculations and thus only reported the surface membrane density. Receptor densities were measured for all orders of branch sizes (1^st^, 2^nd^, 3^rd^ etc.; see below), for both branches dorsal to the central canal (dorsal) and ventral to the central canal (ventral). Nodes were identified with dense bands of Caspr or Na channel labelling (and lack of MBP). Branch points were also identified. We also examined raw image stacks of the neurobiotin afferents and receptors, to confirm that the automatically 3D reconstructed and identified receptors labelled within the afferent (yellow) corresponded to manually identified receptors colocalized with neurobiotin (Fig. 1). This was repeated for a minimum of 10 examples for each condition, and in all cases the 3D identified and manually identified receptors and channels were identical. Many receptors and channels lay outside the afferent, and near the afferent these were difficult to manually identify without the 3D reconstruction software, making the 3D reconstruction the only practical method to fully quantify the receptors over the entire afferent. We also optimized the reconstruction of the neurobiotin filled afferents following the methods of Fenrich^114^, including brightening and widening the image edges slightly (1 voxel, 0.001 µm^3^) when necessary to join broken segments of the afferent in the final afferent reconstruction, to account for the a priori knowledge that afferents are continuous and neurobiotin signals tend to be weaker at the membrane (image edges) and in fine processes. Finally, we also counted the proportion of nodes and ventral boutons innervating motoneurons that contain GABA receptors clusters, as an additional quantification of the receptor distribution. Limitations in the sensitivity of receptor antibody labeling leave open the possibility that we missed small quantities of receptors. Thus, while we find that most proprioceptive afferent terminal boutons lack GABA_A_ receptors^15^ (see Results) there may well still be small quantities of receptors. However, these receptors are unlikely to have much functional impact, since previous direct recording from ventral terminals boutons show little PAD at the time when PAD is observed in more dorsal portions of the same axons^15^.

GABA receptors usually occurred in the axons in distinct clusters. The distances from these receptor clusters to nodes or branch points was measured and average distances computed (from centers of clusters to centre of nodes), from high power confocal images evenly sampled across the axon arbour. Some nodes did not branch, so receptors at these nodes were fairly far from the nearest branch (∼20 µm), making the average receptor to branch point distance larger than the receptor to node distance, the latter which were small because GABA_A_ receptors were mainly only at nodes (see Results). The average distance between the receptor clusters and the nearest axon terminals on the motoneurons was also computed, but this was complicated by the very large distances often involved, forcing us to compute the distances from low power images and relate these to the high power images of receptors sampled relatively evenly along the axon arbour. For this distance calculation, to avoid sampling bias in the high power images, we only admitted images from axon branch segments (1^st^, 2^nd^ and 3^rd^ order, detailed below) that had a receptor density within one standard deviation (SD) of the mean density in branch types with the highest density (1^st^ or 2^nd^ order ventral branches for GABA_A_ receptors and 3^rd^ order ventral terminal branches for GABA_B_ receptors; i.e. images from axons branches with density above the dashed confidence interval lines in Fig. 1e were included; this SD computed from densities of pooled axons from all rats, rather than single rat averages, to better reflect the axon density variability). This eliminated very large distances being included from branch segments with relatively insignificant receptor densities. We also confirmed these calculations by computing the weighted sum of all the receptor distances weighted by the sum of the receptor density for each branch type (and divided by the sum of all receptor densities), which further eliminated sampling bias. This gave similar average distance results to the above simpler analysis (not shown).

#### Sensory axon branch order terminology

The branches of proprioceptive axons were denoted as follows: dorsal column branches, 1^st^ order branches that arose of the dorsal column and project toward the motoneurons, 2^nd^ order branches that arose from the 1^st^ order branches, and 3^rd^ order branches that arose from the 2^nd^ order branches. Higher order branches occasionally arose from the 3^rd^ order branches, but these were collectively denoted 3^rd^ order branches. First and second order branches were myelinated with large dense clusters of sodium channels at the nodes in the myelin gaps, which were characteristically widely spaced. As the second order branches thinned near the transition to 3^rd^ order branches, they became unmyelinated, and at this point sodium channel clusters were smaller and more closely spaced (∼6 µm apart, not shown). These thinned branches gave off 3^rd^ order (and higher) unmyelinated terminal branches with chains of characteristic terminal boutons that terminated on motoneurons. The 1^st^ order branches gave off 2^nd^ order branches along most of their length as they traversed the cord from the dorsal columns to the motoneurons, but we separately quantified 1^st^, 2^nd^ and 3^rd^ order branches in more dorsal (including dorsal and intermediated laminae) and ventral (ventral laminae) regions of the cord.

#### Node identification

Nodes in myelinated axon segments nodes were identified either directly via direct Na channel clusters and paranodal Caspr, or indirectly by their characteristic paranodal taper. That is, in the paranodal region the neurobiotin filled portion of the axon tapered to a smaller diameter, likely because the Caspr and presumably other proteins displaced the cytoplasmic neurobiotin, which also made the intracellular neurobiotin label less dense (Fig. 1b, black regions in taper). Regardless of the details, this taper made nodes readily identifiable. This taper forces the axial current densities to increase at the nodes, presumably assisting spike initiation, and consistent with previous reconstructions of myelinated proprioceptive afferents ^115^.

#### GAD2 neuron labelling

GABA_axo_ neurons that express GAD2 were visualized by genetically tagging them with Cre-ER driven fluorescent reporters. Usually we used the ChR2-EYFP reporter to both insert ChR2 and label with EYFP. This ChR2 construct is membrane bound and so does not fill soma or large processes making cells sometimes hard to visualize. Thus, in some animals we additionally included the Cre driven tdTom reporter, which is a cytoplasmic reporter that fills the entire cell to help visualize the complete anatomy of the entire GABA_axo_ neuron (Fig 3). In this case, GAD2 neurons should have both EYFP (green in Fig 3) and tdTom (red) reporter labelling. However, the balance of green and red expression intensity was variable, with some processes with more EYFP and others with more tdTom, leading to some axons more one color than the other. This was likely due to a number of factors. First, membrane bound fluorophores are easier to see in small diameter axons or dendrites, because of a higher membrane-to-cytoplasm ratio, making red more visible in small axons. Second, variability in tissue penetration of the antibodies we used to amplify the reporter signals and more intense ChR2-EYFP labelling (green) in axons may have led to variable red and green intensity. Finally, genetic variability in the Cre-ER driven reporter expression, which only occurs transiently after the tamoxifen administration, may explain why a small proportion of neurons are either just green or just red, with one reporter not expressed by this transient Cre expression. Expression of only one reporter happened in only a small proportion of neurons, but when it did our double reporter method is an advantage in visualizing these neurons.

### Computer simulations

All computer models and simulations were implemented in NEURON ver7.5 ^116^. The geometry and myelination pattern of the model were extracted from a previous study that used serial-section electron microscopy to generate about 15,000 photomicrographs to reconstruct a large myelinated proprioceptive Ia afferent collateral in the cat (Nicol and Walmsley, 1991)^115^. This structure was used in a prior modeling study ^25^. Four classes of segment were defined in the model: myelinated internodes, nodes, unmyelinated bridges, and terminal boutons. Data from 18 of the 83 segments were missing from the original study. The missing data were estimated using mean values of the same segment class. The cable properties of the model were determined from diameter-dependent equations previously used for models of myelinated axons^117^ and included explicit representation of myelinated segments using the double cable approach^117–119^. Hodgkin-Huxley style models of voltage gated sodium (transient and persistent) and potassium channels were adopted from a previous study, at 37°C^117^. All three voltage-gated conductances were colocalized to unmyelinated nodes and segments throughout the modelled axon collateral. The density of sodium and potassium conductances was adjusted to match the size and shape of experimentally recorded action potentials. To be conservative, sodium channels were placed at each node and bouton (gNa = 1 S/cm^2^), even though bouton immunolabelling for these channels was not common in our terminal bouton imaging (Fig. 1), since disperse weak sodium channel labelling may have been missed. Removing these bouton sodium channels did not qualitatively change our computer simulation results (not shown). Current clamp stimulation was applied to the middle of the first myelinated segment (pulse width 0.1 ms, amplitude 2 nA; near dorsal root) to initiate propagating action potentials in the model. Voltage at multiple sites of interest along the collateral was measured to assess propagation of action and graded potentials through branch points. Transient chloride conductance (i.e. GABA_A_ receptors) was modeled using a double-exponential point process (Eq. 1); parameters were manually fit to experimental data. GABA_A_ receptors were localized to nodes at branch points to match experimental data. The amplitude and time course of the modeled PAD (also termed PAD) was measured from the first myelinated internode segment, similar to the location of our intra-axonal recordings.

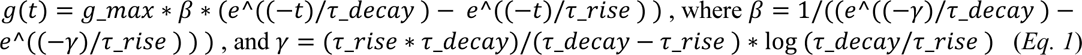

The parameters at all synapses were the same: time constant of rise (𝜏_rise_) = 6ms, time constant of decay (𝜏_decay_) = 50ms, default maximum conductance (*g_max_*) = 1.5nS (varied depending on simulation, see figure legends), and chloride reversal potential (*E*_Cl_ ^-^) = -25mV (i.e. 55 mV positive to the resting potential to match our experimental data) ^15^. Space constants (λ_S_) were computed for each segment of the afferent, from subthreshold current injections (100 ms) on the distal end of each branch segment and fitting an exponential decay (with space constant λ_S_) to the passive depolarization along its length, and then repeating this with current injected in the proximal end to get a second λS, and finally averaging these two space constants.

## QUANTIFICATION AND STATISTICAL ANALYSIS

Data were analyzed in Clampfit 8.0 (Axon Instruments, USA) and Sigmaplot (Systat Software, USA). A Student’s *t*-test or ANOVA (as appropriate) was used to test for statistical differences between variables, with a significance level of *P <* 0.05 (two tailed). Power of tests was computed with α = 0.05 to design experiments. A Kolmogorov-Smirnov test for normality was applied to the data set, with a *P <* 0.05 level set for significance. Most data sets were found to be normally distributed, as is required for a *t*-test. For those that were not normal a Wilcoxon Signed Rank Test was instead used with *P <* 0.05. Categorical data was instead analyzed using Chi-squared tests, with Yate’s continuity correction used for 2 × 2 contingency tables and again significant difference set at *P <* 0.05. Effects in male and female animals were similar and grouped together in analysis. For in vivo experiments, a single data point is taken from the average response in each subject/animal and *n* values indicate subject number. Axons and motoneurons were recorded ex vivo from widely separated locations (one segment apart or contralateral) within the whole spinal cord, and are considered independent; so statistics were performed across all neurons (*n*) from all animals, though all main effects were confirmed to occur in each animal, and comparing across animal averages also showed significant changes (*n* animal numbers). Data are indicated as box plots representing the interquartile range and median (thin line) and error bars representing the 90^th^ and 10^th^ percentile, interpolated between nearest points (Cleveland method). Mean also shown as thick line in boxes.

## Data availability

All data are available in the manuscript or the supplementary materials. Raw data are available upon request to the corresponding authors. This study did not generate data sets or new unique reagents.

## Code availability

The computer code used to perform the axon simulations (Extended Data Fig. 5) are publicly available on the github repository: https://github.com/kelvinejones/noah-axon.git

## Acknowledgements

We thank Leo Sanelli, Jennifer Duchcherer, Babak Afsharipour and Christopher K. Thompson for technical assistance, and Shawn Hochman, CJ Heckman, FJ Alvarez and Tia Bennett for discussions and editing the manuscript. VGLUT1^Cre^ mice cords were kindly donated by Dr. Francisco J. Alvarez. We thank Prof Uwe Rudolph (McLean Hospital, currently University of Illinois Urbana-Champaign) for providing Gabra5-floxed mice. This research was supported by the Canadian Institutes of Health Research (MOP 14697 and PJT 165823 D.J.B.) and the US National Institutes of Health (NIH, R01NS47567, D.J.B. and K.F.; R01GM118801, R.A.P.).

## Author information

Contributions. K.H, and A.M.L-O. designed the study, carried out the animal experiments and analyzed data. K.M. and M.A.G. designed and performed the human experiments. N.P. and K.E.J. designed and performed the computer simulations. S.L., S.B., A.M., M.J.S. and R.S. assisted with animal electrophysiology. K.F. and A.M.L-O provided confocal microscopy. K.K.F, S.L., D.A.R. and K.H. developed the transgenic mice and performed the optogenetic experiments. R.A.P. developed transgenic mice used in immunohistochemical studies. Y.L. and D.J.B. conceived and designed the study, carried out experiments and analyzed data. D.J.B, K.H., and Y.L. wrote the paper, with editing from other authors. These authors contributed equally: Krishnapriya (Veni) Hari and Ana M. Lucas-Osma. These authors jointly supervised this work as senior authors: Yaqing Li, Keith K. Fenrich and David J. Bennett.

Corresponding author. David J. Bennett

## Ethical declarations

All authors declare no competing interests.

## Supplementary information

**Extended Data Fig. 1.**
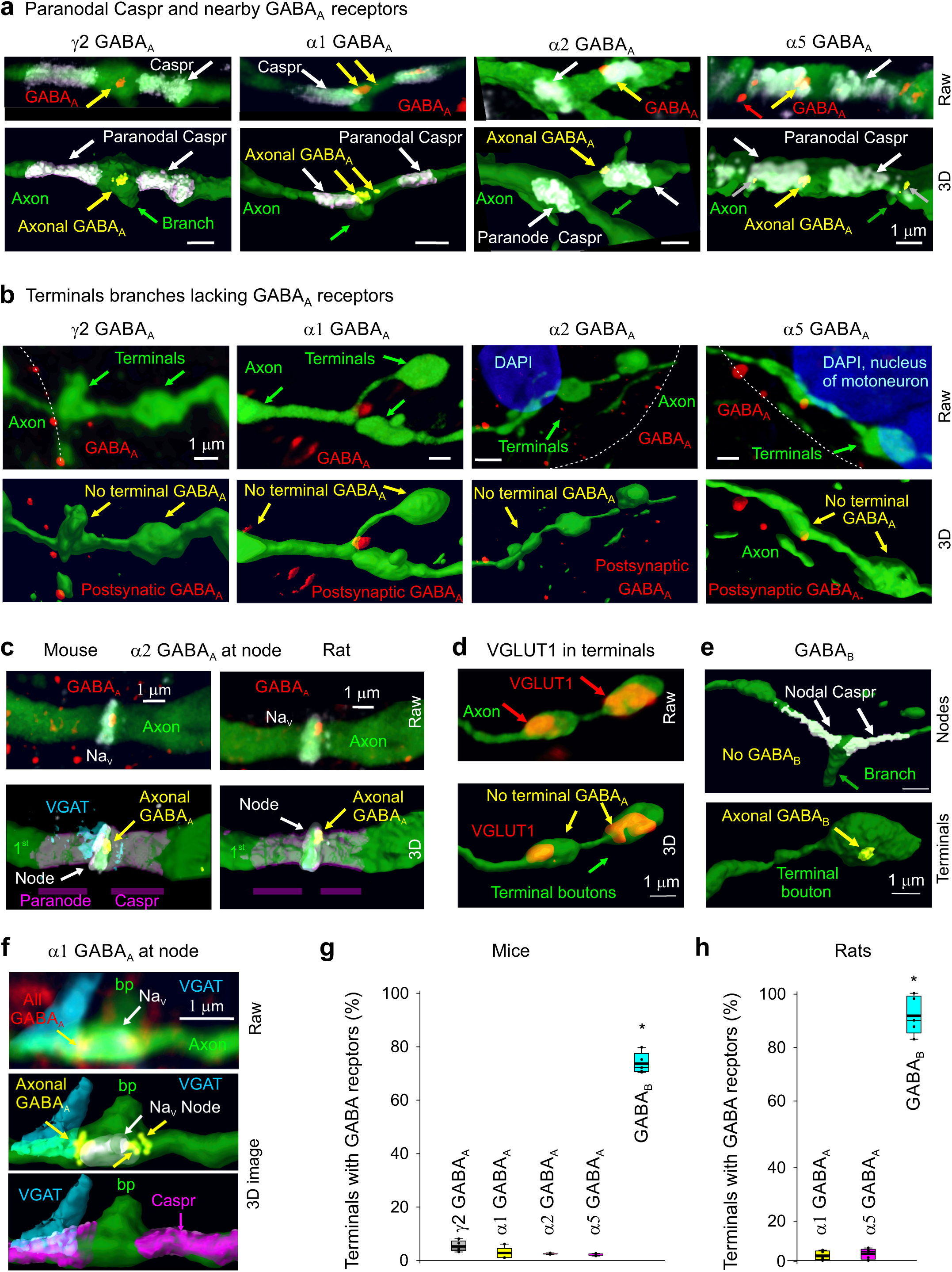
Nodal and not terminal GABA_A_ receptors in mice and rats. **a-b**, In the sacrocaudal spinal cord of mice we examined the distribution of GABA_A_ receptor subunits on nodes and terminals of sensory axons, including extrasynaptic α5 subunits, synaptic α1 and α2 subunits, and ubiquitous γ2 subunits (e.g. forming the common α1βγ2 or α5βγ2 receptors, though less common extrasynaptic α1βδ have been reported)^121-124^. We genetically labelled primary sensory axons by their expression of the vesicular glutamate transporter VGLUT1 with a reporter in VGLUT1^Cre/+^; R26^LSL-tdTom^ mice (tdTom reporter displayed as green, for consistency with Fig. 1). VGLUT1 is mainly only expressed in sensory axons^59^, especially ventral proprioceptive afferents, as other afferents do not reach the ventral horn^15^. Axons were reconstructed in 3D as detailed in Fig. 1. 3D reconstructed nodes of Ranvier on myelinated 2^nd^ order ventral branches are shown identified by paranodal Capsr immunolabelling (**a**, bottom), along with raw confocal image stacks (maximum projection from z-stack) prior to 3D reconstruction (**a**, top; receptors red). As in rats, nodes were often near branch points (green arrow). Terminal boutons from the ventral horn are likewise shown with raw and 3D rendered images (**b**). GABA receptors colocalized with the axon are labelled yellow in the 3D reconstructions. Receptor clusters specifically in the plasma membrane are indicated by yellow arrows in (**a**). Similar to rats, the α5, α1, and γ2 GABA_A_ receptor subunits were found on large axon branches (1^st^ and 2^nd^ order) in the dorsal, intermediate, and ventral cord, near nodes (**a**), but not on ventral horn terminal boutons (**b**, 3^rd^ order). As also seen in rats (Fig 1e), synaptic GABA_A_ receptors were usually in single large membrane bound clusters at nodes, whereas extrasynaptic α5 receptors were often broken up into multiple clusters, with the largest clusters in the membrane (yellow arrows), and smaller cytoplasmic clusters several µm from the edge of the node under the paranodal Caspr (grey arrows; Fig 1f). Cytoplasmic α5 receptors have been reported previously^109^. Many receptors were in neighboring neurons (red arrows), and in our previous publication^15^ these and cytoplasmic axon receptors may have been mistaken for nodal receptors in the axon membrane, though this is corrected here with evaluation of higher resolution confocal images. The presence of these α5, α1, and γ2 GABA_A_ subunits is consistent with their mRNA previously reported in the dorsal root ganglion^125^. Also, the finding of α1 subunits on these axons is consistent with the recent observation that α1 is only on myelinated sensory axons, rather than unmyelinated C fibres^126^. **c**, Synaptic α2 GABA_A_ receptor labelling in mouse and rat axons (rat axon labelled as in Fig 1) with nodes labelled with both antibodies to sodium channels Na_V_ and Caspr, to confirm the relation of receptors to the node and paranodal region. Receptors are often at the transition between the node and paranodal Caspr, as here. Putative GABAergic synaptic contact labelled with VGAT. **d**, Ventral terminal boutons in VGLUT1^Cre/+^; R26^LSL-tdTom^ mouse (with reporter labelling the complete axon, green) immunolabelled with VGLUT1 to verify that these are afferent terminals, which have vesicular VGLUT1^+^ protein expression. Immunolabelling for γ2 GABA_A_ receptors again showed that terminals lacked this ubiquitous subunit that makes up most GABA_A_ receptors. **e**, Immunolabelling for GABA_B_ receptors on 3D reconstructed sensory axons, with same format and mice of (**a-b**). GABA_B_ receptors were generally absent from nodes identified by paranodal Caspr, but present on ventral terminal boutons, as in rats (Fig. 1). Similar results (**a- e**) were obtained from n = 5 mice. **f**, Synaptic α1 GABA_A_ receptor labelling at a rat Ia afferent node labelled with both antibodies to sodium channels Na_V_ and Caspr, to confirm the relation of receptors to the node and paranodal region in rat, like in mouse (**c,** same node as in left of Fig 1c, but Na_V_ and raw images included). Putative GABAergic synaptic contact labelled with VGAT, where left red arrow shows GABAA receptor contacting VGAT at edge of node. VGAT is near, but does not contact Caspr (in 3D view), but may well contact the paranodal myelin loops, since oligodendrocytes express GABA receptors^38^. Node is at dorsal 2^nd^ order axon branch point. **g-h**, Box plots of the proportion of ventral terminal boutons with GABA receptors in both mice (**g,**VGLUT1^+^) and rats (**h**), again showing that few boutons contain GABA_A_ receptors, whereas many contain GABA_B_ receptors. * significantly more GABA_B_ than GABA_A_ receptors, n = 5 animals per condition, with 70 - 120 terminals examined per animal and receptor.

**Extended Data Fig. 2.**
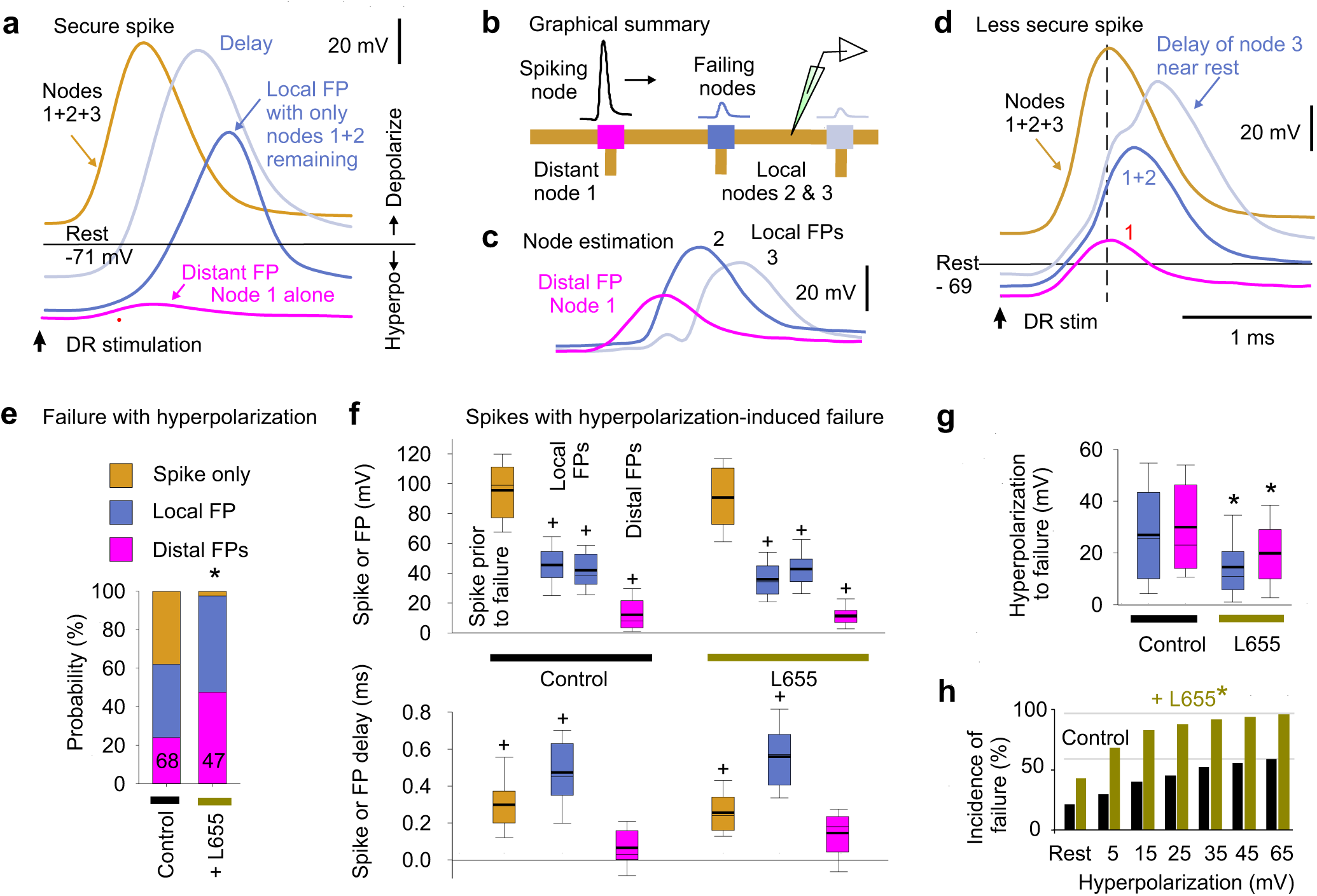
Voltage dependence of nodal spike propagation failure following dorsal root stimulation. **a-d**, Intracellular recording from proprioceptive Ia afferent branches in the rat dorsal horn with secure spikes at rest, evoked by DR stimulation (1.1xT, 0.1 ms; T: afferent volley spike threshold; sacral S4 DR; **a, d**). Spike failure was induced by increasing hyperpolarization (failure near rest in **d**, but not **a**), with a delay and then abrupt loss of height, reflecting failure of successively further away nodes (**a, d**). Of the two local nodes adjacent to the electrode, one failed first with hyperpolarization, leaving the attenuated spike from the other node (local FP, about 1 λ_S_ away), which eventually failed as well with further hyperpolarization, leaving a much smaller FP from more distal nodes (distal FP). Spike failure in a node was always proceeded by a delay in the nodal spike. Estimated contributions from local nodes (**b-c**) were computed by subtraction from traces in (**d**). Spike attenuation was down to about 1/e by the second failure, consistent with the space constant λ_S_ being two internodal distances (**c**), or about 90 µm. Note that due to attenuation of injected current with distance, larger hyperpolarizations were needed to stop spikes with more distal vulnerable nodes (secure spikes), as observed by smaller distal FPs (**a** verses **d**), and some axon spikes could not be stopped (not shown; Spike only, quantified in **e**). Data collected in discontinuous current clamp (DCC) mode so electrode rectification during the injected hyperpolarizing current did not affect potential (DCC switching rate 7 kHz, low pass filtered at 3 kHz to remove switching artifact, which also removed stimulus artifact). **e**, Distribution of spiking in branches with full spikes only, one local nodal spike (local FP), or a distal nodal spike (distant FP) remaining after maximal hyperpolarization, and * significant change when blocking α5 GABA_A_ receptors with L655708 (0.1 - 0.3 µM) using χ-squared test, *P <* 0.05; *n* = 68 control and *n* = 47 L655708 treated axon branches from 7 rats. **f**, Box plots of spike or FP heights and delays in branches and rats indicated in (**e**), measured just prior to spike failure (or at maximal hyperpolarization for secure spikes) and after failure (for local and distal FPs, as induced in **a-d**). Delay measured relative to peak of spike at rest. + FP significantly different than spike at rest, *P <* 0.05. **g**, Box plots of the hyperpolarization needed to induce failure, in axon branches and rats from (**e**). * significant change with L655708, *P <* 0.05. **h**, Incidence of failure at varying potentials (% of total spikes from **e**). * significantly more failure with L655708, χ-squared test, *P <* 0.05. Note that secure spikes in rats and mice are not overall different when measure by spike height at rest (Figs. 2g and 3g). However, the spike height shown here in (**f**) is the height while the cell is hyperpolarized far from rest, and so is much larger than at rest, since spikes generally overshoot to near the reversal potential for sodium, and so spike heights while hyperpolarizing are not comparable to the spikes at rest in mice or rats (measured from holding potential to peak). Thus, during hyperpolarization the nodes produce larger overall spikes, including FPs from local nodes (**a-f**). Also, during hyperpolarization from current injection, the two adjacent nodes to the electrode fail, unlike during natural failures, and thus when one nodes fails the spike only halves in height, leaving the spike from the second adjacent node (**a-b**). Thus, these local FPs are large in relation to the FPs from distal nodes with natural failure (Fig 2).

**Extended Data Fig. 3.**
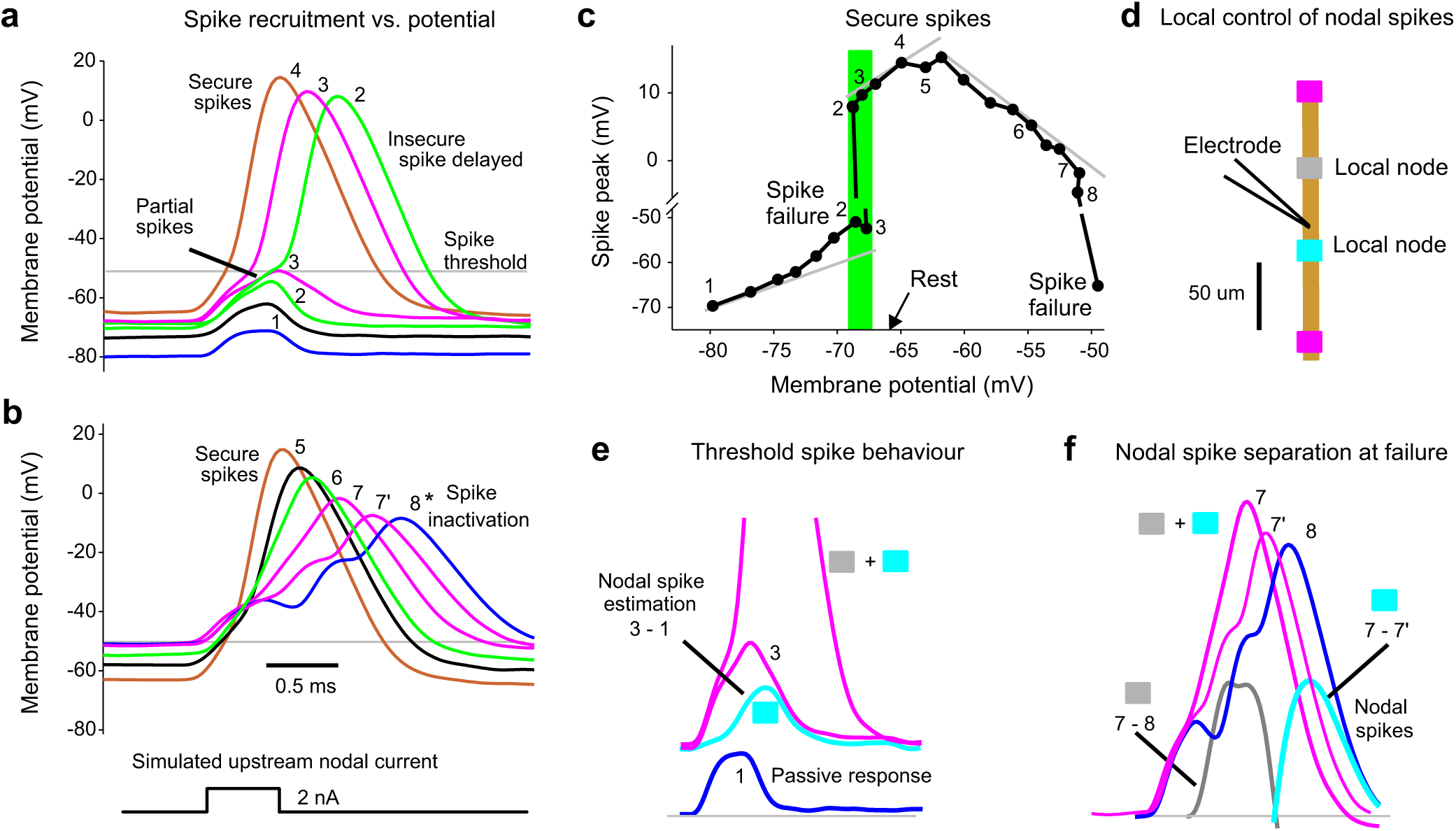
Voltage dependence of spike failure with simulated nodal currents. Simulating spike propagation failure in a proprioceptive axon by applying a brief intracellular current injection to mimic the current arriving from an upstream node (and FP), yielded full spikes evoked at rest, but nearby nodal spikes delayed and then failing as the membrane was held progressively more hyperpolarized with a steady bias current. Also, large steady depolarizations inactivated these spikes, though well outside of the physiological range (> −50 mV). **a**, Intracellular recording from proprioceptive Ia afferent branch in the rat dorsal horn (sacral S4 axon). A brief current injection pulse (0.5 ms) was applied to simulate the current arriving from distal nodes during normal axon spike conduction, and repeated at 1s intervals. During these pulses the membrane was held at varying potentials for 1 – 2 s with steady current injection, with numbers and colours denoting a given holding potential (in DCC mode). At the most hyperpolarized levels spikes failed to be evoked and only the passive response is seen, like a FP (blue, 1). As the potential was depolarized to near the axon’s resting potential (−67 mV) partial spikes occurred (green, 2 and 3), likely from a single adjacent node activating, and then delayed broad spikes occurred, as both adjacent nodes were activated. At more depolarized levels the spikes arose more rapidly and increased in height to full secure spikes (4). **b**, In the same axon as (**a**), at holding potentials well above those seen physiologically (near −50 mV, lower plots) spikes started to exhibit sodium channel inactivation and failure, with a decrease in spike height and delay (7 – 8) and eventually full failure (shown in **c**). Adjacent nodes started failing at slightly different times with different delays, broadening the spike and eventually separating into two distinct nodal spikes (8*). **c**, Spike heights plotted as a function of holding potential, including those spikes illustrated in (**a** and **b**), with spike number- labels indicated. Left grey line indicates passive leak current response, and shows deviation from passive response near rest. Shaded green region shows all or nothing failure or spikes near the resting potential. Middle grey line shows a region of secure spikes with relatively invariant spikes. Right grey line shows spike inactivation with large depolarizations and outright failure near − 50 mV. Note the split vertical axis. Similar voltage dependence of spike failure occurred for *n =* 5/5 axons showed similar results, from 4 rats. This demonstrates two modes of spike failure: 1) spikes that fail at rest or at hyperpolarized potentials and 2) spikes that fail with large depolarizations above rest. The latter is likely not physiological, since even the largest PAD that we have observed (5 – 10 mV; Fig. 4d) does not depolarize axons to −50 mV, since axons rest near −70 mV, and PAD is only large at hyperpolarized levels (Fig. 4d) and decreases steeply as the potential approaches the reversal potential for chloride (−15 mV)^15^. **d**, Schematic of recording arrangement and relation to adjacent nodes for data in (**e**) and (**f**). **e**, Expansion of responses 1 (blue) and 3 (pink) from (**a**), and difference (cyan) to show the first local nodal spike height at threshold, recorded at electrode. Active nodes from schematic in (**d**) shown with shaded boxes. **f**, Spikes near sodium inactivation from (**b**) (7 and 8), with differences indicating local nodal spikes (grey: 7 − 8, and cyan: 7 – 7’, both truncated to only show estimated nodal spike). Nodes likely arranged as in (**d**).

**Extended Data Fig. 4.**
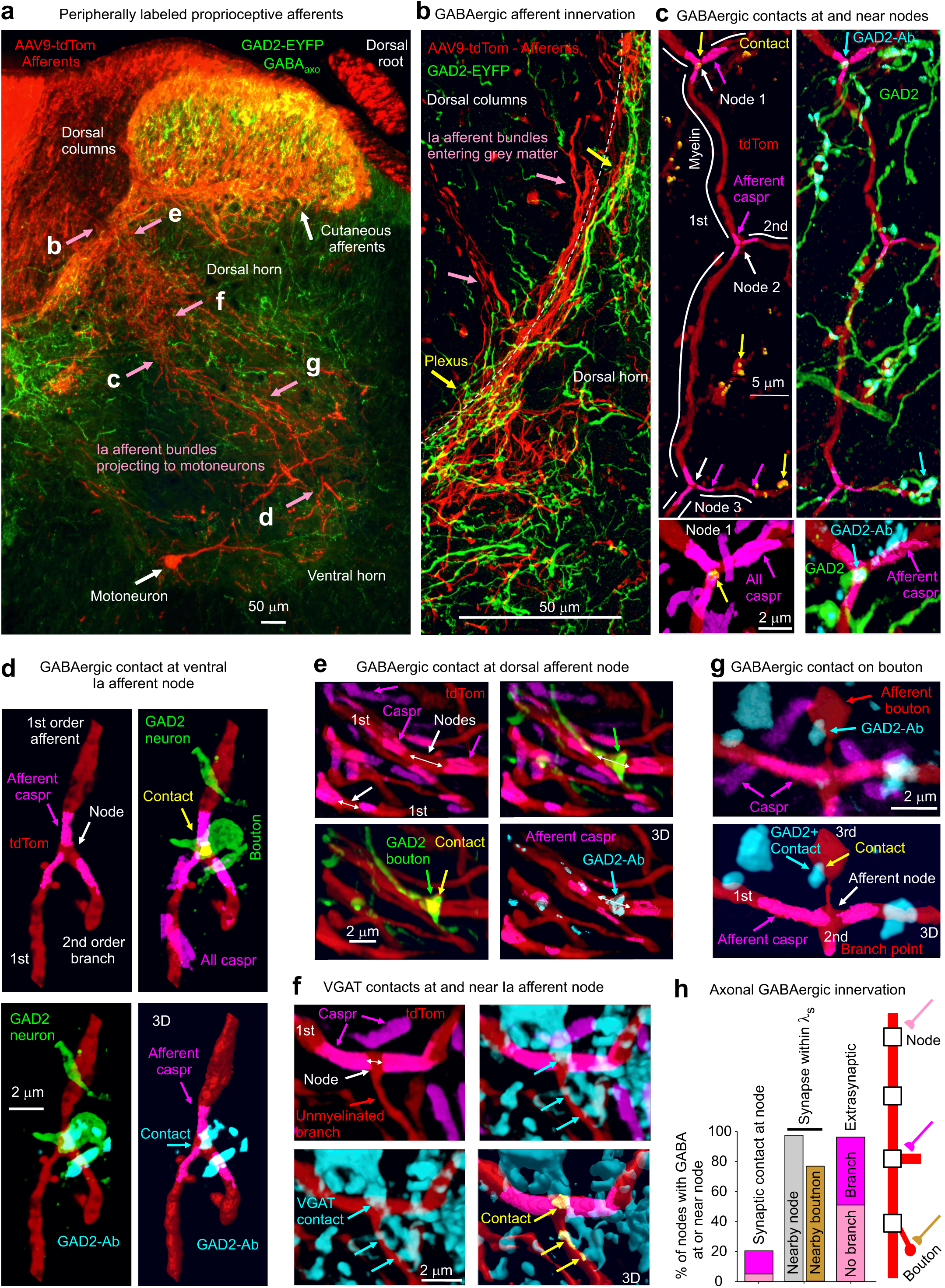
GABAergic innervation of nodes and nearby boutons in viral vector labelled proprioceptive afferents. **a**, Sensory afferents labelled in the spinal cord and dorsal root of an adult GAD2//ChR2-EYFP mouse (indicated GAD2-EYFP, green) by a peripheral viral vector injection (adeno-associated virus, AAV9-tdTom, red; dosing detailed in Methods), shown from a confocal microscope image of a transverse spinal cord section. Large myelinated proprioceptive Ia afferents were identified by their characteristic extensive ventral horn branching and unique innervation of motoneurons (as in Fig 1a). These Ia afferents left the dorsal columns in bundles that were readily traced to the ventral horn in serial sections, with a characteristic S-shaped projection path from the dorsal columns to the motoneurons (as in Fig 3j)^127^, allowing their identification from the point where they enter the dorsal horn to their terminals in the ventral horn (pink arrows). The relatively sparce afferent labelling helped facilitate this tracing of afferent innervation. The peripheral AAV9- tdTom injection occasionally also labelled one motoneuron in a transverse (lower white arrow), though this only occurred a couple times per spinal cord segment, making them possible to distinguished from afferents elsewhere, but conveniently identifying the motor nucleus. No other central neurons were labelled, indicated that the viral vector did not substantially cross the blood brain barrier and thus mainly only affected peripheral sensory and motor axons, as we previously found^43^. EYFP^+^ cells (GABA_axo_ neurons) were only in the dorsal and intermediate laminae (GAD2-EYFP^+^), with projections into the ventral horn and dorsal columns along the entire path of the Ia afferent bundles, as detailed in Fig 3. Many confocal images were stitched together to cover the entire spinal cord (tile scan), with afferents in the the ventral region shown with 50% greater intensity, to visualize the fine afferent terminals. The approximate regions where images in (**b-g**) were taken are indicated in (**a**), but in different transverse sections, oriented similarly. **b**, GABA_axo_ neurons formed a dense plexus (yellow arrows) that wound around Ia afferents as they branched out of the dorsal columns into the grey matter (at dashed line). **c**, Main 1^st^ order myelinated branch of a Ia afferent in intermediate laminae with three nodes at branch points labelled with axonal caspr, approximate myelin regions marked with white lines, and GABA_axo_ neurons seen wrapping the axon (top images). Node 1 had a GAD2-EYFP+ contact that had presynaptic GAD2 labelled with an antibody (GAD2-Ab, cyan; yellow marks contact of afferent, GAD2-EYFP and GAD2-Ab computed in 3D), whereas Nodes 2 and 3 lacked contacts, though the short branch arising off of Node 3 had an unmyelinated terminal branch with a GAD2^+^ contact. All nodes were well with λ_S_ of a GAD2^+^ contact on the same axon. Also, all nodes were close (< 5 µm) to GAD2^+^ terminals, providing the potential for extrasynaptic GABA. Expanded view of Node 1 shown at bottom left, where all caspr labelling is displayed, including in overlapping nearby axons, whereas in other images only the axonal caspr in the 3D volume of the afferent is shown for clarity, together with raw images from other antibodies. Expanded view of Node 1 contact shown on bottom right, where all GAD2-ab labelling is shown, whereas in other images only GAD2-Ab in the GAD2-EYFP neuron volume is shown. Not all GAD2-Ab was in GAD2-EYFP^+^ neurons, likely due to variability in tamoxifen induced cre or transport of EYFP in GAD2//Ch2R-EYFP mice. **d**, Nodal contact in the ventral horn, shown with similar format to (**c**), with again a GABA_axo_ neuron wrapping around the axon and specifically making a contact at the node (yellow; EYFP+, GAD2-Ab+). **e**, Two nodes on two 1^st^ order Ia afferent branches in the dorsal horn plexus of (**b**), again delineated by paranodal Caspr, one of which had a GABA_axo_ neuron contact (GAD2-EYFP, marked yellow again) with presynaptic GAD2-Ab (cyan, shown in 3D volume of GABA_axo_ neuron) and the other did not. **f-g**, Nodes from 1^st^ order Ia afferent branches, identified by paranodal Caspr, in control GAD2//ChR2-EYFP mouse without tamoxifen injection, and so lacking EYFP, but inhibitory innervation examined with VGAT or GAD2 antibodies. In (**f**) the node had a direct VGAT^+^ contact and nearby contacts on the small unmyelinated branch arising from the node (contacts computed in 3D labelled yellow). In (**f**) the node lacked a direct GABAergic contact, but had a nearby GAD2-Ab^+^ contact on a bouton of a short unmyelinated branch arising from the node, enabling nodal depolarization well within the space constant of the axon (λ_S_ = 91 µm). **h**, Quantification of the fraction of total nodes with direct synaptic GABAergic contacts (nodal contacts, GAD2+, VGAT+; ∼20%), nearby contacts at a node or unmyelinated terminal branch/bouton on the same axon (within λ_S_ = 90 µm; 98 − 77%) and putative extrasynaptic innervation (within 5 µm, ∼95%) from GABA_axo_ neurons, with presynaptic GABA inferred from GAD2 or VGAT immunolabelling, in *n* = 3 mice from 53 nodes each on 1^st^ and 2^nd^ order branches of Ia afferents. Synaptic contacts occurred at unbranched (pink) or branched (magenta) nodes, as did extrasynaptic innervation, with about half the nodes branched overall. Synaptic contacts that occurred on short unmyelinated afferent terminal branches arising from the node usually occurred at a bouton and are labelled as: Nearby bouton.

**Extended Data Fig. 5.**
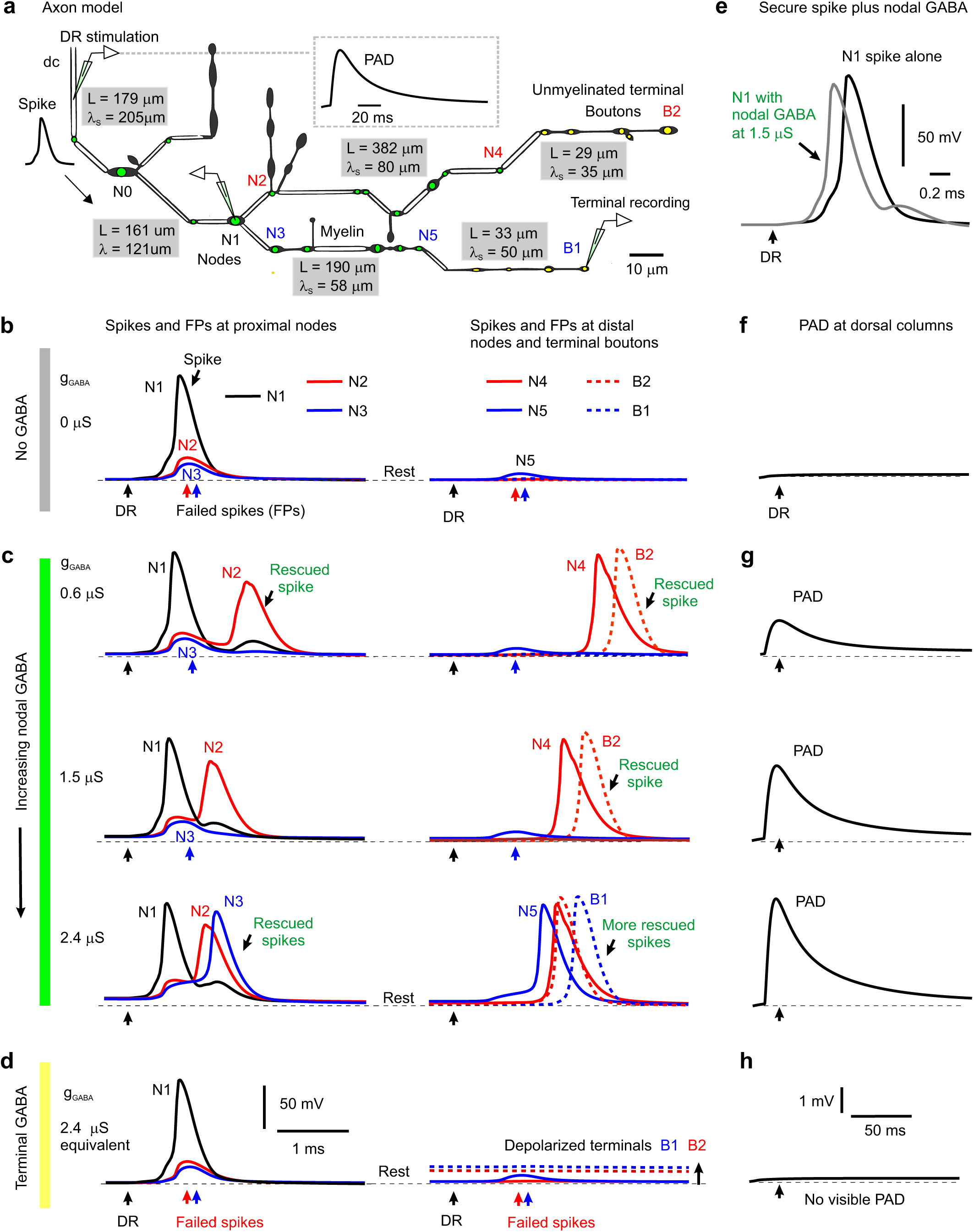
Computer simulation of branch point failure and rescue by GABA. **a**, Model of a 3D reconstructed proprioceptive afferent, drawn to scale, except myelinated branch lengths all shortened an order of magnitude. Double line segments are myelinated (white) and the rest unmyelinated. Adapted from anatomical studies of Walmsley^25^. Nodes are indicated with a green dot, and ventrally projecting terminal boutons indicated with a yellow dot. As in our axons of Fig. 1, branch points were always at nodes. GABA_A_ receptors of equal conductance (nS) were placed at each node and associated branch points, and total dorsal columns (dc) depolarization from phasically activating these receptors is shown in inset. The branch lengths (L) and computed space constants (λ_S_) are indicated in gray boxes for each segment of the afferent, the latter computed from subthreshold current injections into each segment. From left to right the gray boxes are for segments spanning from the dorsal columns (dc) to N0, N0 to N1, N1 to N5, N1 to N4, N4 to B2 and N5 to B1. Average space constant was λ_S_ = 91 µm, similar to in other axons^39^. **b**, Responses to simulated dorsal root (DR) stimulation (0.1 ms pulse, 2 nA, at black arrows) computed at various downstream branch points (nodes) and terminal boutons in the spinal cord, with resting potential indicated by thin dashed line (−82 mV). A sodium spike propagated to the branch point at node N1, but failed to invade into the downstream branches, leaving nodes N2 and N3 with only a passive depolarization from the N1 spike (failure potential, FP). Further downstream nodes and terminals experienced vary little depolarization during this failure (N5, B1 and B2). In this case the GABA conductance was set to zero (g_GABA_ = 0, control), simulating a lack of GABA tone. Note that only the nodes beyond the parent branch at node N1 failed due to the conductance increases in large daughter branches to nodes N2 and N3 that drew more current than node N1 could provide (shunting conductances). Node 3 is particularly interesting as it does not itself branch, though a neighbouring node has a small branch, both contributing to the overall conductance and related failure, and Node 2 has two adjacent branches contributing to its conductance. Other branch points with relatively smaller conductance increases (N0, simpler branching) did not fail to conduct spikes. Generally speaking, if the upstream node of a parent branch cannot provide enough current to activate the nodes of its daughter branches then spikes fail, and this is especially likely with multiple sequential branch points, like in N1 – N2. However, failure even occurs at daughter nodes than themselves lack branches (N3), so GABA receptors are useful in aiding spikes at unbranched nodes. **c**, As seen experimentally, nodal GABA_A_ receptor activation to produce PAD prior to the DR stimulation (∼ 10 ms prior; as in Fig. 4f) rescued spikes from failing to propagate. That is, with GABA_A_ receptors placed just at nodes and associated branch points a weak phasic activation of these receptors (conductance g_GABA_ = 0.6 − 1.5 µS per node shown) rescued conduction down the branch to node N2, with full nodal spikes seen at the distal node N4 and the terminal bouton B2 (DR stimulation at peak of PAD, detailed in (**g**) with black arrows indicating DR stimulation timing). A larger GABA receptor activation (2.4 nS) additionally rescued spike conduction down the branch to node N3, with full spike conduction to the distal node N5 and the terminal bouton B1. Note that increasing GABA conductance sped up the arrival of distal spikes (e.g. at N4 and B2), by up to 1 ms, suggesting substantial variation in sensory transmission times induced by GABA, as we see experimentally. Also note that this nodal GABA depolarized the nodes (N1 – N3) relative to rest (thin dashed line), thus assisting spike initiation. In contrast, nodal GABA did not depolarize the terminal boutons (B1 and B2), consistent with our recent direct recordings from terminals^15^. Sensitivity analysis revealed similar results with a wide range of sodium channel and GABA receptor conductances, though increasing sodium conductance sufficiently prevented failure all together (not shown, but like in Extended Data Fig. 10f). Interestingly, when we put GABA receptors only at node N2 (or at the unmyelinated bouton immediately above N2) then the spike propagating though N2 to the terminal bouton B2 was rescued with the same GABA conductance (0.6 µS, not shown), as in the main simulation (**c**). Likewise with GABA receptors only at node N3 and nowhere else, then the spike was rescued at that node N3 (at 2.4 µS; not shown). **d**, When instead we removed all nodal GABA receptors and instead place them on terminal boutons (near B1 and B2, yellow, with equivalent total conductance, 2.4 µS condition), then activating them did not rescue the spike propagation failure, since the associated depolarization of nodes is too attenuated at the failure point (N1-N3; no change from resting potential). The GABA receptors did depolarize the terminal boutons (B1 and B2, thick dashed lines) substantially relative to the resting potential (thin dashed lines), but this depolarization was sharply attenuated in more proximal nodes (N1-3). **e**, Reduction of spike height (shunt) and speeding of spike onset with increasing GABA conductance at a non-failing node (N1; model with nodal and not terminal bouton GABA conductances, **c**), consistent with actual recordings from axons in Fig. 3d and Extended Data Fig. 8a. **f-h**, PAD recorded at the dorsal columns (dc) during conditions in (**b**-**d**), respectively, as experimentally recorded dorsal PAD. A phasic GABA induced depolarization (PAD) was generated by changing GABA conductances, g_GABA_, as detailed in Methods, and GABA receptor location varied as in (**c-d**). DRs were stimulated at the peak of this PAD in (**c-d**). Note that nodal (**g**) but not terminal (**h**) GABA_A_ receptors caused a visible depolarization (PAD) at the dc, due to less electrotonic attenuation over a shorter distance to the dc.

**Extended Data Fig. 6.**
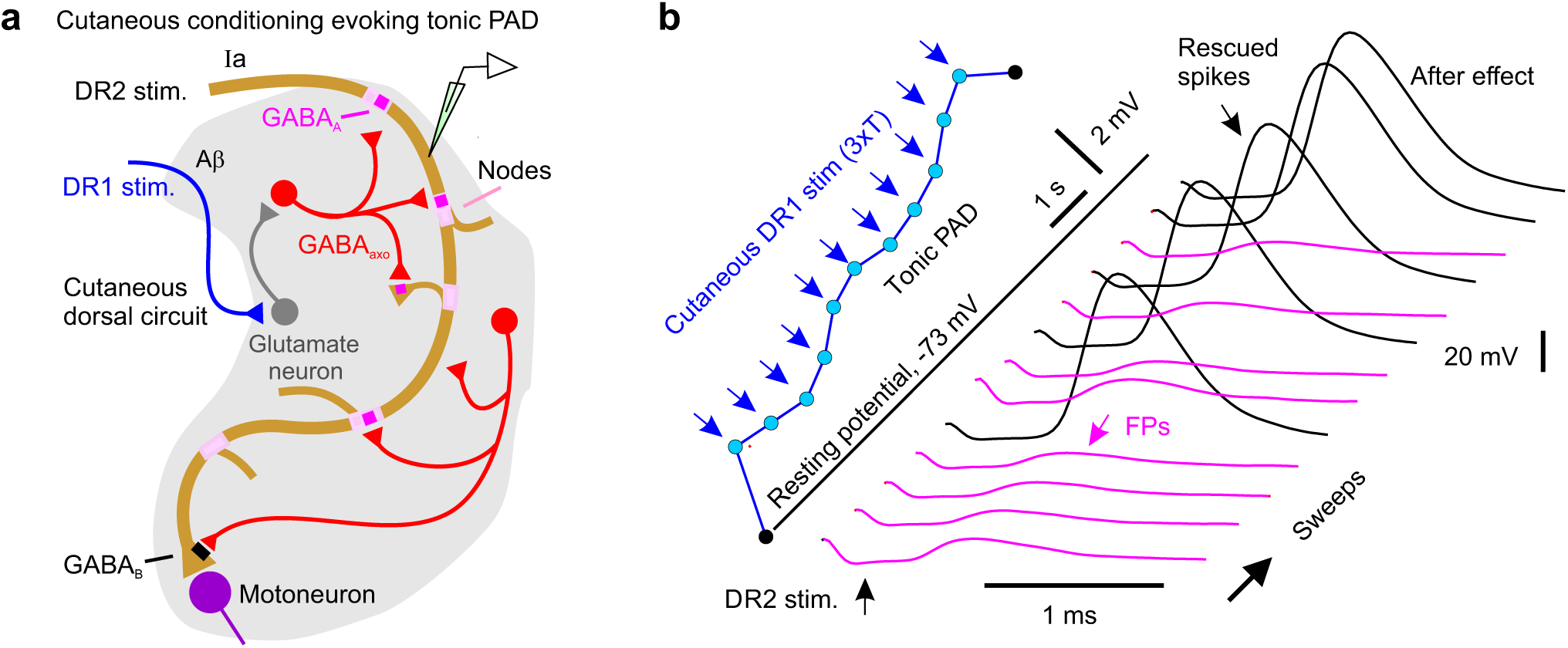
Cutaneous driven trisynaptic circuits mediating PAD and assisting repetitive firing. **a**, Cutaneous driven dorsal trisynaptic circuit mediating PAD. A minimally trisynaptic circuit is classically known to depolarize afferents via GABA_axo_ neurons. This circuit involves sensory afferents activating excitatory intermediary neurons (glutamatergic) that in turn activate GABA_axo_ neurons that return to innervate sensory axons^13, 128^. Even though GABA_axo_ neurons are small^5^ this circuit influences afferents over widespread regions of the spinal cord^15^. Specifically, the activation of a small group of sensory axons in just one DR or nerve causes this trisynaptic circuit to produce a widespread activation of many axons across the spinal cord, even many segments away and across the midline^15^. This allowed us to activate PAD from adjacent roots without directly activating an axon in a particular root, as detailed in Fig. 4 and the rest of this figure. One variant of this classic trisynaptic circuit specifically involves cutaneous stimulation activating dorsal intermediary neurons^13, 14^ that activates GABA_axo_ neurons (likely dI4 neurons^128^) that in turn innervate cutaneous afferents, which we term the cutaneous dorsal circuit. While this cutaneous dorsal circuit also synaptically innervates some proprioceptive afferents^129^ (**a**), its main action on proprioceptive afferents is to produce a pronounced extrasynaptic spillover of GABA that depolarizes these afferents tonically via α5 GABA_A_ receptors (termed: tonic PAD, L655708 sensitive), especially with repetitive cutaneous nerve stimulation (1 - 200 Hz) that leads to minutes of depolarization^15^, and we see similar tonic PAD here (detailed next). **b**, Intracellular recording from a proprioceptive axon branch in rat dorsal horn (sacral S4 axon, DR2). The axon branch spontaneously exhibited spike propagation failure when its was stimulated alone (denoted DR2 stimulation, repeated at 1 Hz, 1.1xT, 0.1 ms; T: afferent volley spike threshold), with only a small failure potential (FP) visible (lower pink traces). Activation of a largely cutaneous DR (caudal Ca1 DR, innervating the tip of the tail, stimulation at intensity for cutaneous afferents, 3xT, 0.1 ms; denoted DR1) evoked a slowly rising tonic PAD when repeated at 1 Hz (blue). When the axon stimulation (DR2 stimulation) was combined with the repeated cutaneous stimulation (DR1, 60 ms prior to each DR2 stimulation) the slowly building PAD prevented spike failure (black spikes), and this outlasted the cutaneous stimulation (after effect). Similar results obtained in *n =* 20/20 axons tested from 10 rats.

**Extended Data Fig. 7.**
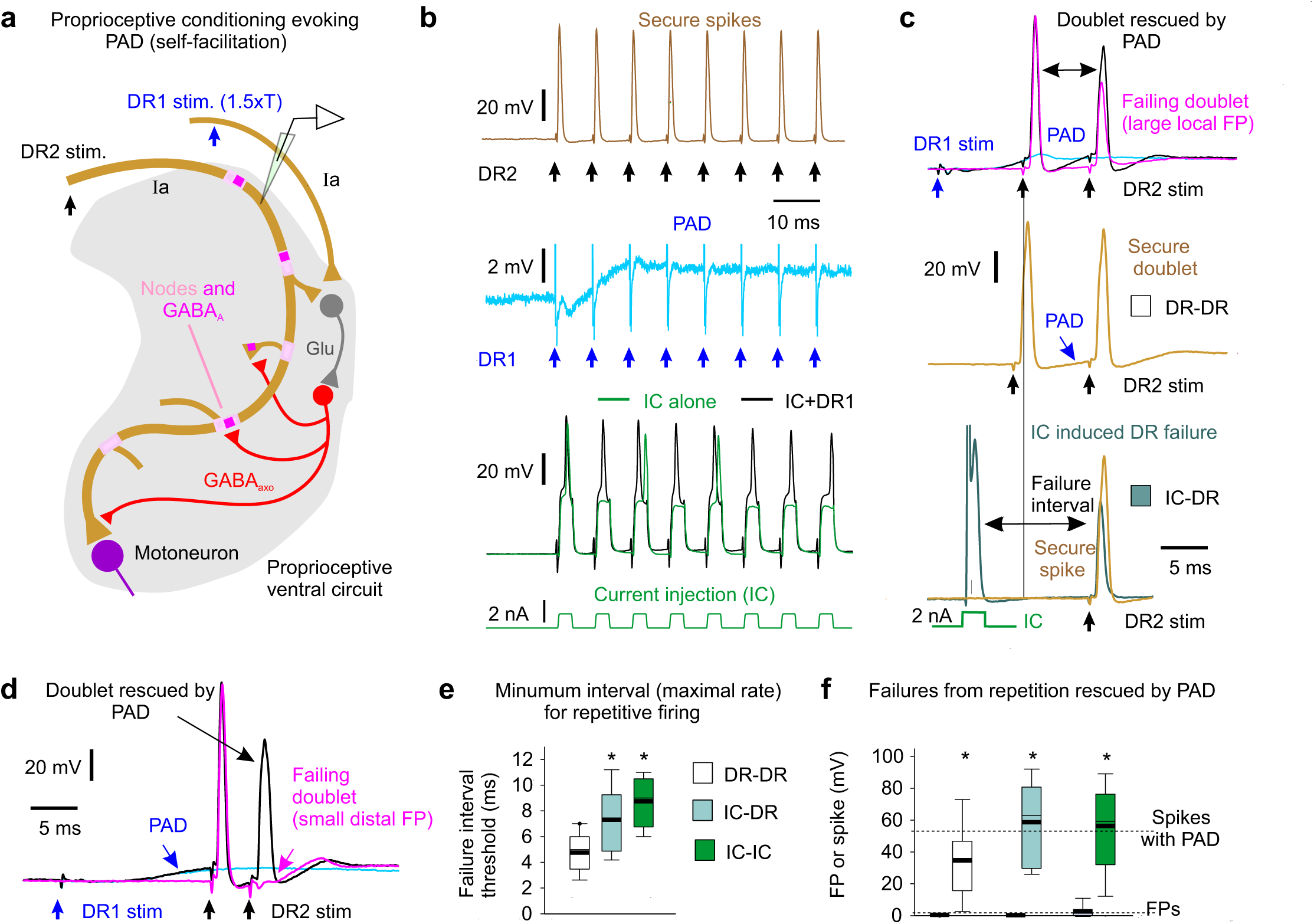
Proprioceptive driven trisynaptic circuit for PAD enabling high frequency spike transmission. **a**, Proprioceptive driven ventral trisynaptic circuit mediating PAD. Another variant of the classic trisynaptic circuit involves proprioceptive afferents activating excitatory intermediary neurons (glutamatergic) that then activate GABA_axo_ neurons that innervate these same afferents, including ventral terminal regions of the afferents^13, 15, 128^. This circuit is more ventrally located compared to the cutaneous dorsal PAD circuit^13^, and thus, we term it the proprioceptive ventral PAD circuit. It likely involves GABA_axo_ neurons that are from the dIL_A_ population^128^. Importantly, the circuit is recurrent, with proprioceptive afferents causing self-facilitation of themselves (via homonymous PAD). It produces a fast phasic axon depolarization (phasic PAD, fast synaptic; Fig. 4c), as well as a slower tonic depolarization (tonic PAD, Fig. 4c, likely from extrasynaptic GABA spillover), as detailed previously^15^. Since proprioceptive sensory axons naturally fire at high rates^130^ where they are vulnerable to spike failure (Fig. 2e-f), we examined the action of self-facilitation by GABA on this failure. **b-f**, During rapid repetitive stimulation of a DR to evoke spikes in a proprioceptive axon there was an inevitable activation of PAD from low threshold proprioceptive axons (homonymous PAD, **b**). This PAD helped spikes fire at high physiological rates of up to 200-300 Hz (5 – 3 ms intervals) before spike inactivation and failure occurred because, in absence of PAD, isolated repetitive activation of the axon with intracellular current pulses (IC) led to failure at much lower firing rates (∼100 Hz; longer spike intervals; **b, e**), even after just two stimuli (doublets; **c, e**). Additional PAD evoked by simultaneous stimulation of an adjacent DR (2xT; T, spike threshold observed from afferent volley) reduced failure from fast repeated IC stimuli (**b, f**), repeated DR stimuli (doublet, **c, d, f**) or hybrid IC-DR stimulation pairs (**f**). Legend details are as follows: **b**, Intracellular recording from proprioceptive axon in rat dorsal horn (sacral S4 axon) with spikes securely evoked by fast repeated DR stimulation (top, 1.1xT, 0.1 ms, sacral S4 DR, denoted DR2; resting potential −68 mV), but spikes failing intermittently with repeated intracellular current injection (IC) at the same rate (bottom green), due to sodium channel inactivation. The reason that spike failure does not occur with the fast DR stimulation is that it is accompanied by a build up of tonic PAD (from self-activation) that helps prevent failure, because adding to the IC stimulation a simultaneous conditioning stimulation of other proprioceptive afferents in an adjacent DR (DR1 stim at 1.5xT, 0.1 ms, S3 DR) prevents spike failure (black trace, bottom), via the proprioceptive ventral circuit (**a**). This DR1 conditioning stimulation does not directly activate spikes in the axon, but it causes a fast depolarization (phasic PAD) that rapidly helps spikes (as early as 6 - 10 ms later), and a building tonic depolarization (tonic PAD) with repetition that further helps later spikes in the stimulation train (DR1 stimulation alone blue, middle trace). Similar results obtained in *n =* 7/7 axons from 4 rats. Likely similar tonic PAD and associated increased spike conduction helps explain post-tetanic potentiation of the monosynaptic EPSP, as previously suggested^131^. **c**, Repeated DR stimulation at higher rates eventually causes spike failure in proprioceptive axons (sodium spike relatively refractory), as shown in the top panel where a double stimulation (doublet, S4 DR, denoted DR2-DR2, 1.1xT, 0.1 ms, resting at −75 mV) exhibits failure on the second spike (with large FP indicated, magenta). However, additional PAD provided by stimulating an adjacent DR (DR1; 1.5xT, group I intensity, 0.1 ms) about 10 ms earlier helps prevent this spike failure (black trace; blue trace: PAD alone). When the same axon was stimulated slightly slower (with a longer doublet interval, second plot, DR-DR) failure did not occur, which we designate the failure interval threshold, which is quantified in (**e**). The self-activated PAD caused by the first DR stimulation in this doublet helped prevent failure in second DR stimulation because replacing the first DR stimulation with an intracellular stimulation (IC, 2 nA) to activate the spike leads to failure of the second spike evoked by the DR stimulation at much longer intervals (lower trace, IC-DR). **d**, Another example of a failed doublet spike (DR2-DR2 stim, 1.1xT, 0.1 ms) that was rescued by PAD evoked by adjacent DR stimulation (DR1 1.5xT, 0.1 ms, resting at −78 mV), as in the top plots of (**c**), except that in this case the failure is at a more distal node, since the FP is small. **e**, Failure interval threshold (minimum firing interval prior to failure, or maximal firing rate) with DR doublets (DR-DR), IC doublets (IC-IC) or IC-DR pair stimulation. Note the shorter intervals possible with the PAD evoked by the first stimulation (DR-DR). * significantly longer than minimum DR doublet interval (DR-DR), *n =* 18 axons each condition from 5 rats, *P <* 0.05. **f**, Quantification of the FP heights that were induced by a fast doublet (DR-DR or IC-IC; *n =* 14 axons each from 5 rats, at failure threshold interval) or IC and DR stimulation (IC-DR; *n =* 11 axons from 5 rats), and the rescue of spikes by PAD evoked by adjacent DR stimulation for each condition. * significant increase in height with PAD, *P <* 0.05.

**Extended Data Fig. 8.**
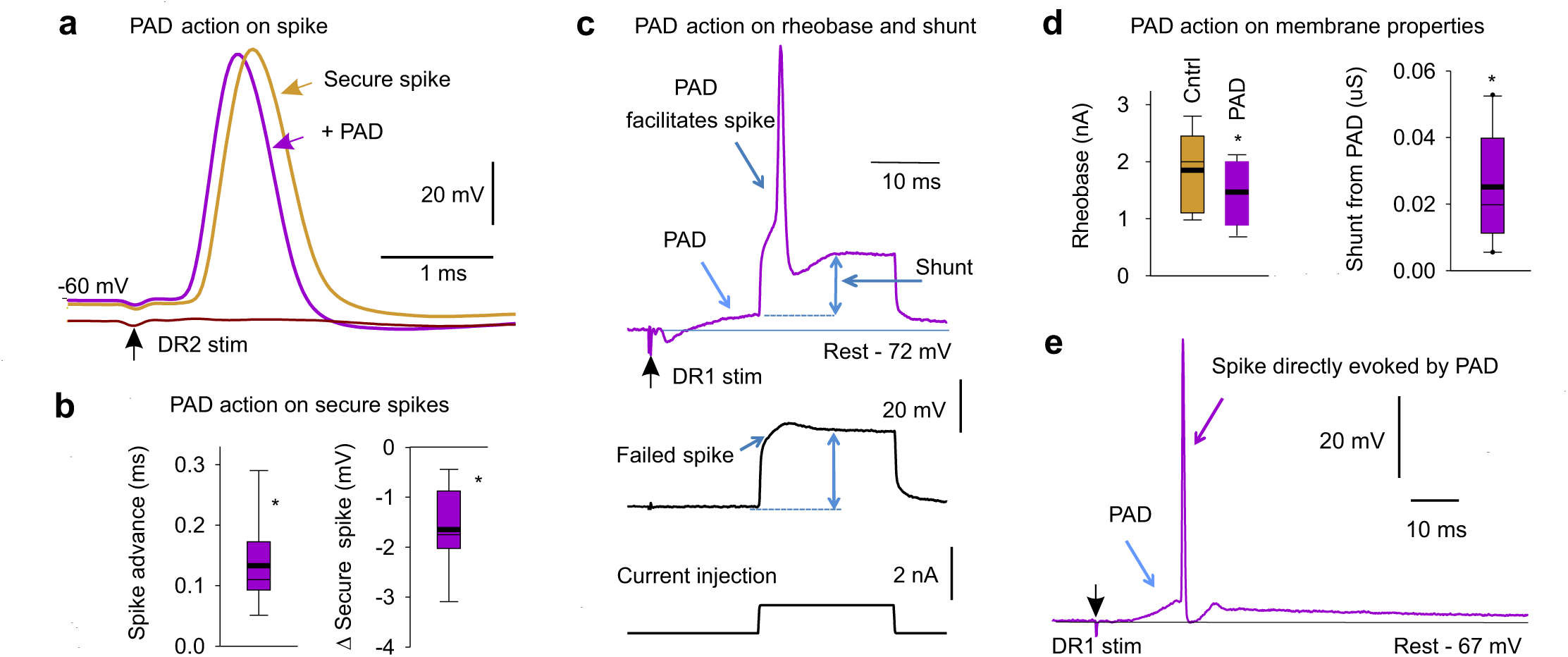
Other excitatory actions of GABA_A_ receptors on proprioceptive axons. **a**, Intracellular recording from a rat sacral S3 proprioceptive Ia afferent branch in the dorsal horn with a secure spike evoked by S3 DR stimulation at rest (DR2, 1.1xT, 0.1 ms, − 60 mV rest, rat; T: spike threshold in afferent volley). Sensory- evoked PAD initiated by stimulating an adjacent DR (DR1; S4 DR; 2xT, 0.1 ms pulse, as in Fig. 4) 10 ms prior to the DR2 stimulation only moderately influenced the spike (DR1 stimulation time not shown). It sped up the spike latency and rise time, reduced the fall time and slightly reduced the spike height. Hyperpolarization induced spike failure (lower trace), as in Extend Data Fig. 2a. **b**, Summary box plots of change in spike peak latency (advance) and height with prior sensory PAD activation as in (**a**). *significant change, *P <* 0.05, *n =* 26 axons, from 9 rats. **c**, Intracellular recording from an S3 proprioceptive Ia afferent branch in the dorsal horn with a brief current injection pulse just subthreshold to initiating a spike (near rheobase, bottom trace) at rest, only initiating a passive response with a small failed spike (middle trace). However, prior activation of PAD by stimulating an adjacent DR (DR1 as in A; S4 DR, 2xT, 0.1ms top trace) allowed the same current pulse to evoke a spike (above rheobase). The passive response to the current injection (double blue arrow; resistance R = V/I) was decreased during the PAD, corresponding to an increase in conductance, that contributed to a shunt (reduction) of the currents generating the spike, though this only caused about a 1% drop in spike height (1 mV; **b**). DCC recording mode, as in Extend Data Fig. 2a-d. **d**, Summary box plots of rheobase (current threshold from **c**) before and during PAD, and change in shunt (conductance = 1/R) with PAD, as in (**c**). * significant change with PAD, *P <* 0.05, *n =* 37 axons from 11 rats. **e**, By itself sensory evoked PAD sometimes initiated a spike on its rising phase, when the DR stimulation was large enough, demonstrating a direct excitatory action of GABA_A_ receptors, as previously reported^15^. These spikes propagate antidromically toward the DR; and are thus termed dorsal root reflexes (DRR). Example shown of intracellular recording from rat sacral S3 proprioceptive afferent branch in the dorsal horn, with PAD produced by a DR1 stimulation (S4 DR stimulation 3.5xT, 0.1 ms) evoking a spike that propagates out the DR (DRR). These DRR occurred in n = 11/120 axons from 15 rats (9% incidence). These PAD evoked spikes occur with a variable latency of 10 – 30 ms^15^ and thus make axons refractory for about 30 ms after the DR stimulation^132^. PAD evokes spikes with a high incidence (79%)^15^, but these spikes fail to propagate antidomically, yielding the 9% incidence we see. However, these spikes are more likely to travel othodromically (up to 79% incidence) and evoke EPSPs in motoneurons via the monosynaptic pathway^8^, and thus also produce a post activation depression of the EPSPs for many seconds. We thus kept the PAD low when examining the effects of PAD on EPSPs, to avoid these spikes and their subsequent inhibitory action (in Figs. 5 – 6).

**Extended Data Fig. 9.**
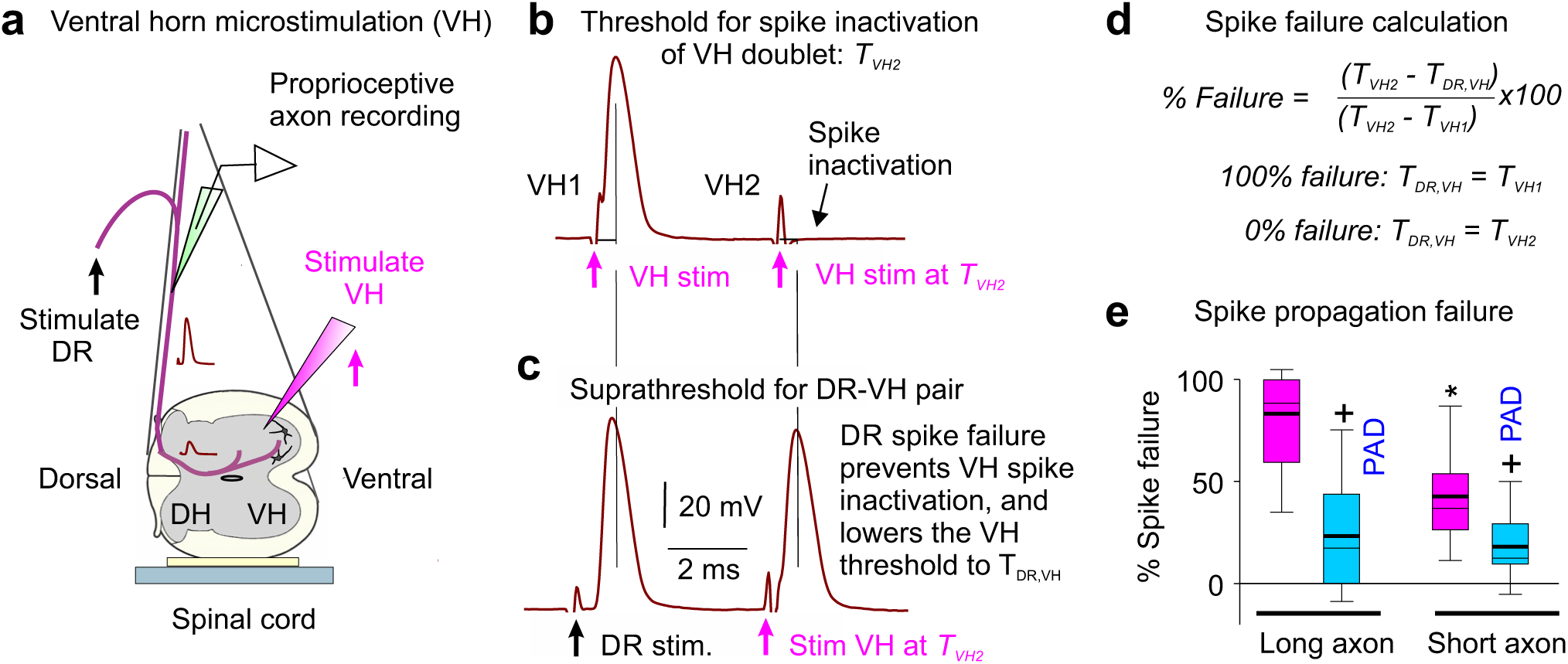
Estimating the overall spike conduction failure from the dorsal root to the motoneurons. **a**, Experimental setup to indirectly measure sensory axon conduction failure following DR stimulation, by examining whether failed axon segments are relatively less refractory to activation after failure, using a double pulse method adapted from Wall^30^. A tungsten microelectrode (12 MΩ) was placed in the ventral horn (VH) near the sensory axon terminals on motoneurons (S3 or S4 VH), to activate the branches/nodes of the axon projecting to the motoneuron that may have failed (VH stimulation). Spikes from VH or DR stimulation were recorded intracellularly in a proprioceptive axon penetrated in the dorsal columns. **b**, *VH threshold in refractory period*. Rapidly repeated VH stimulation (VH doublet; two 0.1 ms pulses) at an interval short enough to produce spike inactivation on the second stimulation (4 ms), with stimulus current adjusted to threshold for inactivation, T_VH2_. This T_VH2_ (∼15 uA) was always higher than the threshold VH stimulation for evoking a spike with the first stimulation, T_VH1_ (∼10 uA, not shown). Recorded in sacral S4 afferent resting at −72 mV, with doublets repeated at 3 s intervals to determine current thresholds. **c**, *VH threshold after DR stimulation*. Similar repeated activation of the axon in (**b**), but with the first activation from a DR stimulation (at 1.5x DR threshold) and the second from VH stimulation at the T_VH2_ intensity from (**b**). In this case the VH stimulation readily activated the axon spike, likely because the DR-evoked spike did not propagate to the VH, leaving the silent portion of the axon non refractory. Thus, this VH stimulation evoked spikes with a lower current than T_VH2_, with this lower threshold denoted T_DR,VH_ (∼ 12 µA, not shown). This DR – VH stimulation interval was deliberately set too short for the involvement of PAD (which rises in > 4 ms; Fig. 4). **d**, Computation of spike failure based on changes in VH stimulation thresholds. If the DR-evoked spike entirely fails to propagate to the VH, then the threshold for subsequently activating the VH (T_DR,VH_) should be the same as the threshold without any prior activation (T_VH1_ = T_DR,VH_), whereas if it does not fail then the threshold for activating the VH should be the same as with a VH doublet (T_VH2_ = T_DR,VH_). In between these two extreme scenarios, the DR evoked spike may only partially fail to propagate spikes to the VH; in this case T_DR,VH_ should be between T_VH1_ and T_VH2_, with the difference T_VH2_ - T_VH1_ representing the range of possible thresholds between full failure and conduction. Overall the % conduction failure can be thus quantified as: (T_VH2_ - T_DR,VH_)/(T_VH2_ - T_VH1_) * 100%, which is 100% at full failure and 0% with no failure. **e**, Average spike conduction failure to the VH in proprioceptive axons, and decrease following a DR conditioning stimulation that depolarized the axon (PAD). Box plots of failure estimated as in (**b-d**). Prior DR conditioning to produce PAD (via adjacent S4 or Ca1 DR stimulation at 3xT) reduced the failure estimated 20 ms later by the paired-pulse conduction testing (repeating DR – VH stimulations of **b-d**). DR conditioning itself lowered the thresholds for VH activation, as previously reported (not shown)^32^. We studied two lengths of axons: long axons (intersegmental, *n =* 11 axons, from 5 rats) with the VH stimulation one segment away from the recording site, and short axons (segmental, *n =* 12, from 5 rats) with the VH stimulation near the recording site, in the same segment. + significantly less failure with PAD and * significantly less failure with short compared to long axons, *P <* 0.05.

**Extended Data Fig. 10.**
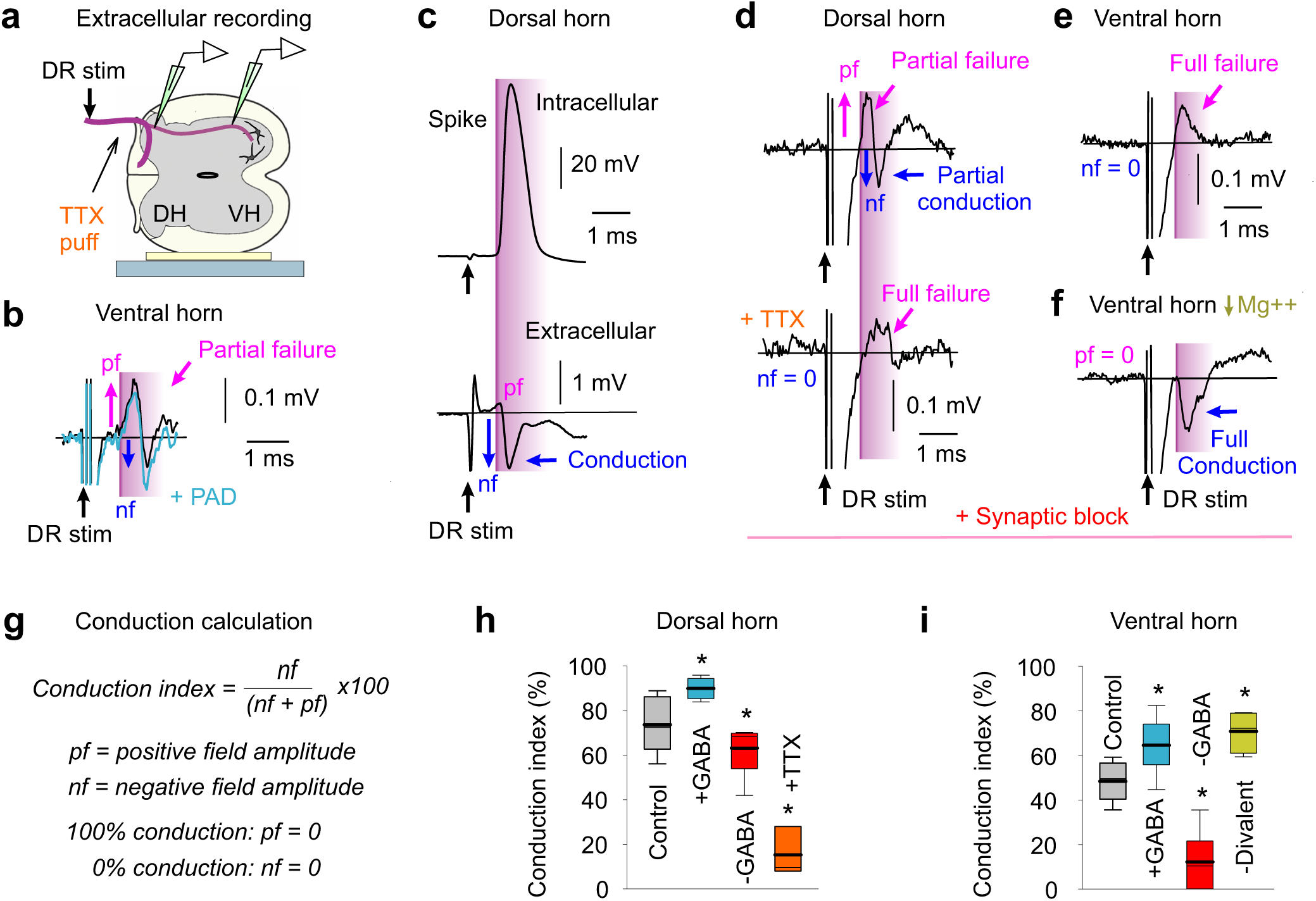
Axonal spike conduction estimated from extracellular recordings. **a**, Experimental setup to directly record spike conduction failure in proprioceptive axon terminals in the ventral horn (VH) following DR stimulation. Extracellular (EC) recordings from axon terminals in VH, with glass electrode positioned just outside these axons, and for comparison EC recording in the dorsal horn (DH). **b**, EC field recorded in VH after DR stimulation (S4 DR, 1.1xT; T: afferent volley threshold), with a relatively large initial positive field (magenta arrows, pf) resulting from passively conducted axial current from sodium spikes at distant nodes (closer to the DR; outward current at electrode), some of which fail to propagate spikes to the VH recording site; thus, this field is a measure of conduction failure, as demonstrated in (**c-f**) below. Following this, a negative field arises (blue arrow, nf), resulting from spikes arising at nodes near the electrode (inward current); thus, this field is a measure of secure conduction. Reducing conduction failure by depolarizing the axon (+PAD) with a prior conditioning stimulation of an adjacent DR (Ca1, 2xT, 30 ms prior), decreased the positive field (pf) and increased the negative field (nf), consistent with increased conduction to the terminals, and in retrospect the same as Sypert et al. (1980)^133^ saw in cat (their Fig. 4). Large stimulus artifacts prior to these fields are truncated. **c**, Control recordings from proprioceptive axons in dorsal horn (DH) to confirm the relation of the EC negative field (nf) to spike conduction. Intracellular (IC) recording from axon (sacral S4, resting at −64 mV) and EC recording just outside the same axon, showing the DR evoked spike (IC) arriving at about the time of the negative EC field (nf). There is likely little spike failure in this axon or nearby axons, due to the very small initial positive field (pf). EC fields are larger in DH compared to VH (G, 10x), and thus the artifact is relatively smaller. **d**, Locally blocking nodes with TTX to confirm the relation of the positive EC field to spike failure. EC recording from proprioceptive axon in the dorsal horn (S4), with an initial positive field (pf) followed by a negative field (nf), indicative of mixed failure and conduction. A local puff of TTX (10 µl of 100 µM) on the DR just adjacent to the recording site to transiently block DR conduction eliminated the negative field (nf) and broadened the positive field (pf), consistent with distal nodes upstream of the TTX block generating the positive field via passive axial current conduction, and closer nodes not spiking. Recordings were in the presence of synaptic blockade (with glutamate receptor blockers, kynurenic acid, CNQX and APV, at doses of 1000, 100 and 50 µM respectively), to prevent TTX spillover having an indirect action by blocking neuronal circuit activity, including GABA_axo_ neuron activity. This synaptic blockade itself contributed to some spike failure, consistent with a block of GABA_axo_ neuron activity, as there was a more prominent positive field (pf) compared to without blockade in (**c**). **e**, EC field recorded from terminals of proprioceptive axons in the ventral horn near motoneurons (S4), in the presence of an excitatory synaptic block that largely eliminates most neuronal circuit behaviour (with kynurenic acid, CNQX and APV, as in **d**). In this synaptic block negative fields were generally absent (nf = 0), and only prominent positive fields (pf) occurred (as with TTX block), suggesting that conduction to the VH often completely failed when circuit behavior is blocked, which likely indirectly reduces GABA_axo_ neuronal circuit activity and its associated facilitation of nodal conduction. **f**, Rescue of spike conduction to the ventral horn by increasing sodium channel excitability by reducing the divalent cations Mg++ and Ca ++ in the bath medium^78^. Same EC field recording as in **e**, but with divalent cations reduced (Mg^++^, 0 mM; Ca^++^, 0.1 mM). The positive field was largely eliminated (pf = 0) and replaced by a negative field (nf), consistent with elimination of conduction failure, and proving that the positive field is not a trivial property of axon terminals^76, 77, 133^. **g**, Conduction index computed from positive (pf) and negative (nf) field amplitudes as: nf / (nf + pf) x 100%, which approaches 100% for full conduction (pf ∼0; as in **c**) and 0% for no conduction (nf = 0; as in **e**). **h-i**, Summary of conduction index estimated from EC field potentials, shown with box plots. Without drugs present in the recording chamber, the axon conduction from the DR to the dorsal horn was about 70% (**h**, *n =* 17 axon fields, from 10 rats), consistent with Fig. 2, whereas conduction from the DR to the VH was only about 50% (**i**, *n =* 11, from 5 rats), suggesting substantial failure at the many branch points in the axon projections from the dorsal horn to the motoneurons. Increasing GABA_axo_ neuron activity with DR conditioning (PAD, 30 – 60 ms prior) increased conduction (+GABA, in both the DH and VH, *n =* 5 and 9 from 5 rats, as in **b**), whereas decreasing GABA and all circuit activity in a synaptic blockade decreased conduction (-GABA, in both DH and VH, *n =* 5 and 6 from 5 rats, as in **d-e**). TTX (*n =* 5 from same rats, **h**) or removal of divalent cations (Mg^++^ and Ca^++^, -Divalent, *n =* 5 from same rats in synaptic block, **i**) reduced or increased conduction, respectively (as in **d** and **f**). * significant difference from control pre-drug or pre- conditioning, *P <* 0.05.

**Extended Data Fig. 11.**
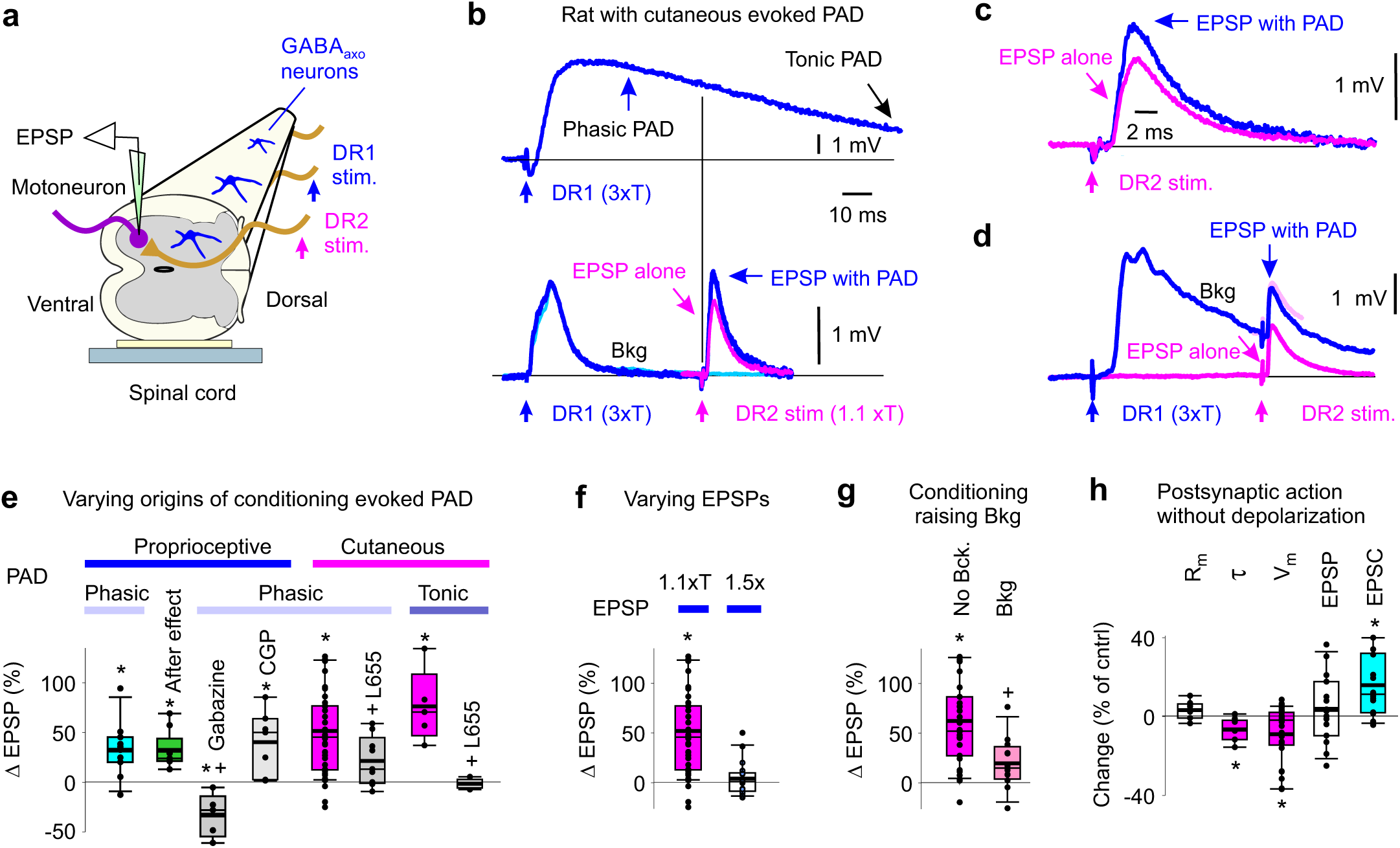
Sensory evoked facilitation of monosynaptic EPSPs by GABA_A_ receptors. **a**, Whole spinal cord ex vivo preparation for intracellular recording of EPSPs from motoneurons while stimulating dorsal roots (DRs). **b**, Monosynaptic EPSP in an S4 motoneuron evoked by a proprioceptive group I stimulation of the S4 DR (1.1xT, denoted DR2, lower traces; resting potential −75 mV: black line; T: EPSP threshold, similar current to afferent volley threshold), alone (pink) and 60 ms after (blue) a conditioning stimulation of cutaneous afferents in rat to evoked PAD (stimulation of the largely cutaneous Ca1 DR, 2.5xT; denoted DR1). Averages of 10 trials each at 10 s intervals. PAD evoked by the same cutaneous conditioning stimulation in a proprioceptive S4 DR afferent is shown for reference (top, recorded separately, as in Fig. 4b). **c**, EPSPs from (**b**) on expanded time scale. **d**, Similar to (**b**), but stronger conditioning stimulation (DR1, 3xT) evoking background postsynaptic activity (blue, Bkg) that lasted longer than 60 ms, and slightly inhibited the EPSP, likely from increased postsynaptic conductances shunting the EPSP (postsynaptic inhibition; light pink: overlay of EPSP alone) and masking nodal facilitation. **e**, Summary box plots of facilitation of EPSPs during phasic PAD evoked by either proprioceptive conditioning (S3 or contralateral S4 DR stimulation, 1.1xT, *n =* 11 motoneurons EPSPs in 5 mice, blue) or cutaneous conditioning (Ca1 DR stimulation, 2-3xT, in rats, *n =* 42 motoneurons/EPSPs in 10 rats, pink), and action of GABA_A_ and GABA_B_ antagonists (gabazine 50 µM, CGP55845 0.3 µM and L655708 0.3 µM grey, *n =* 5, 7, 9 EPSPs respectively in same animals, with again mice proprioceptive conditioning and rats cutaneous). EPSPs evoked in S3 and S4 motoneurons by DR2 (S3 or S4) stimulation at 1.1T, as in (**b**). Facilitation measured 60 ms post conditioning during phasic PAD (phasic condition indicated) and when postsynaptic actions of conditioning (Bkg) were minimal (as in **b**). After conditioning was completed EPSP testing continued and revealed a residual facilitation that lasted for 10 - 100 s (After effect, green, *n =* 9 EPSPs in 5 mice), due to a build up of tonic PAD, after which the EPSP returned to baseline (not shown), similar to post-tetanic potentiation^131^. Also, a brief high frequency cutaneous stimulation train (200 Hz, 0.5 s, 2.5xT) that led to a very long lasting depolarization of proprioceptive axons (Tonic PAD, example in Fig. 5g) caused a facilitation of the monosynaptic EPSP that lasted for minutes (average shown, tonic cutaneous condition), and this was blocked by L655708 (in rats, *n =*5 EPSPs in 4 rats). * *P <* 0.05: significant change with conditioning. + *P <* 0.05: significant change with antagonist. Raw data points shown to indicate occasional inhibition of the MSR by conditioning, but overall facilitation. ChR2 activation of GABAaxo neurons lacked these long tonic PAD-mediated after effects on the EPSP facilitation (Fig. 5e-f, Post), suggesting an additional source of GABA mediating after effects. **f**, Summary box plots of change in EPSP induced by cutaneous DR (DR1) conditioning (and associated phasic PAD) 60 ms prior to evoking the EPSP, with varying EPSP stimulation intensity. When the DR that evoked the test EPSP (DR2) was stimulated at an intensity that produced less than half the maximal EPSP height (1.1xT, ∼ 30% max EPSP, *n =*42, same data as in **e**) the facilitation of EPSP by conditioning was larger than when this DR2 stimulation was increased to produce a test EPSP near maximal (1.5xT, prior to conditioning, *n =* 18 EPSPs from same rats as in **e**). \**P <* 0.05: significant change with conditioning. This is likely because the stronger test stimulation reduced the headroom for increasing EPSPs by recruiting more proprioceptive axons, and increased self-facilitation prior to conditioning, the latter during repeated testing used to obtain EPSP averages. **g**, Summary of cutaneous facilitation of EPSPs from (**f**) (evoked by DR2 at 1.1xT), but separated into trials without (as in (**b**), *n =* 31 EPSPs, in 10 rats) and with (as in (**d**), *n =* 11 EPSPs in 10 rats) large background postsynaptic changes induced by conditioning that lasted up to and during the EPSP testing (at 60 ms post conditioning, Bkg). \**P <* 0.05: significant change with conditioning evoked PAD. + *P <* 0.05: significant reduction facilitation with increased background activity (Bkg). **h**, Remote postsynaptic inhibition from conditioning. Long lasting changes in intrinsic proprieties of motoneurons (S4 and S3) following a mixed proprioceptive and cutaneous conditioning DR stimulation (on S3 or contralateral S4 DR, 2.5xT, DR1) that only produced a transient postsynaptic depolarization that ended prior to EPSP testing (as in B), including a reduction in time constant (τ) and slight hyperpolarization of potential (V_m_), both measured at the time of EPSP testing (measured at 60 ms post conditioning, but in trials without EPSP testing; *n =* 15 motoneurons in 5 rats). At this time, there was little change in somatic membrane resistance (Rm) with conditioning, suggesting that conditioning induced postsynaptic activity at a remote location in distal dendrites of the motoneuron. Indeed, when we voltage clamped the membrane potential during monosynaptic testing (DR2 at 1.1-1.5xT) to directly measure the synaptic current (EPSC) and minimize that inhibitory action of postsynaptic conductance increases, we found that the conditioning stimulation (DR1) produced a larger facilitation of the monosynaptic EPSC than the EPSP measured in current clamp in the same motoneurons (same n = 15 motoneurons). These results are consistent with the facilitation of the EPSP being masked by postsynaptic inhibition from increases in remote dendritic postsynaptic conductances triggered by the conditioning stimulation. \**P <* 0.05: significant change with conditioning.

**Extended Data Fig. 12.**
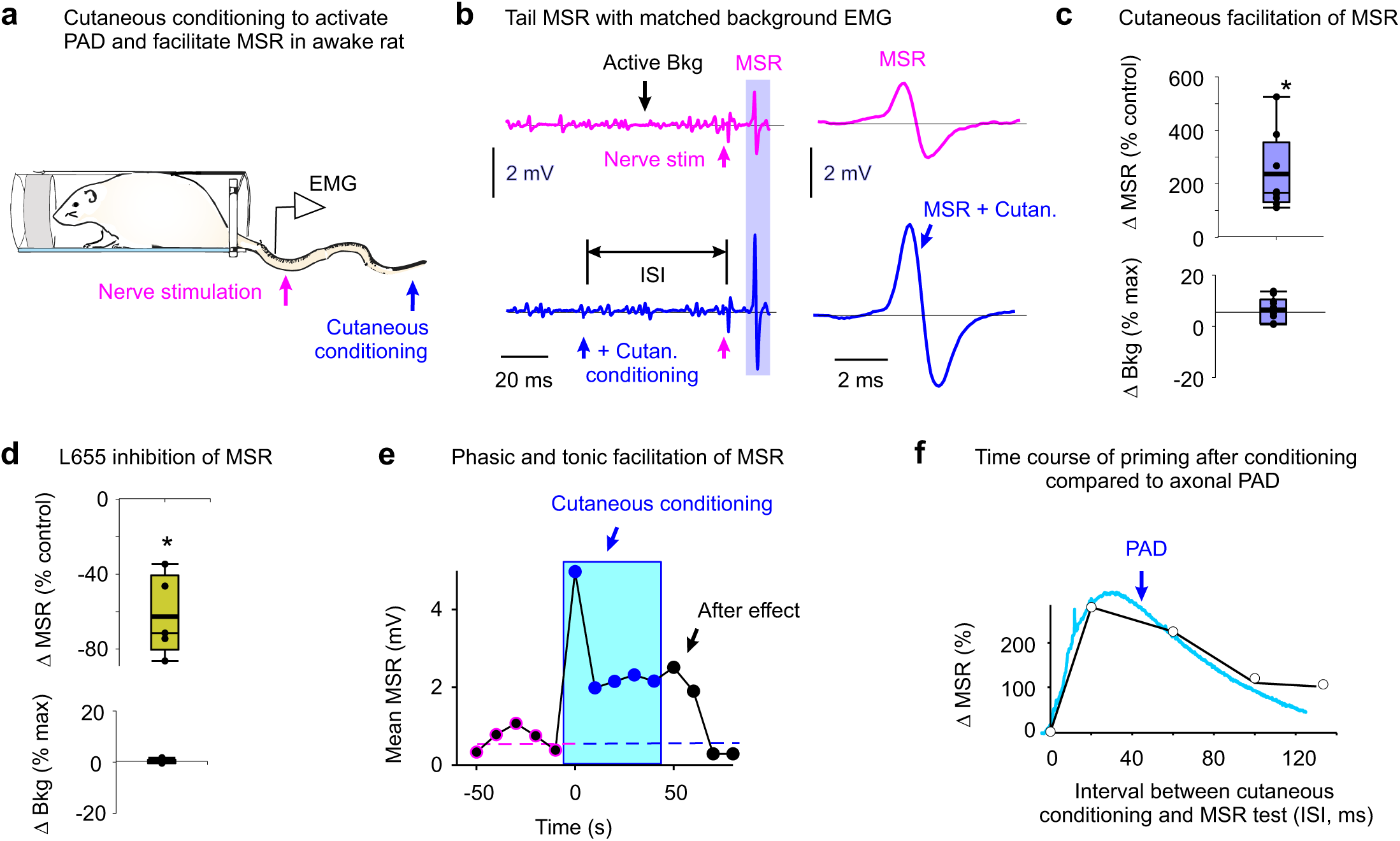
Facilitation of reflexes in awake rats. **a-c**, MSR recorded as in Fig 6, but in rat and with PAD instead activated with cutaneous conditioning (tip of tail, 0.2 ms, 2xT, 60 ms prior, 0.1 Hz repetition), at matched active Bkg EMG. * significant change with conditioning, *P <* 0.05, *n =* 8 rats (**c**). **d**, Decrease in MSR with L655708 (1 mg/kg i.p.) at matched Bkg EMG. Box plot. * significant change, *P <* 0.05, *n =* 5 rats. **e**, Typical MSR amplitude before, during and after conditioning as in (**a-c**) with after effect. **f**, Typical change in MSR with cutaneous conditioning as in (**a-c**) when the ISI is increased, compared to PAD (from Fig. 4). (**e,f**) similar results in *n =* 5/5 rats. **Summary of findings in awake rats**: Increasing GABA_axo_ neuron activity with a brief cutaneous stimulation (**a**) increased the MSR (**b-c**) during a period consistent with nodal facilitation by PAD (30 – 200 ms post stimulation; **f**). We again kept the conditioning stimulation small enough to not change the background (**b**) to rule out postsynaptic actions. Blocking GABA_A_ receptor tone (with L655708) decreased the MSR, at matched levels of background EMG (**d**), suggesting a spontaneous tonic PAD facilitating the MSR. Repeated cutaneous conditioning stimulation (trains) to induce a buildup in this tonic PAD caused an associated buildup of the MSR that outlasted the conditioning and its postsynaptic actions by many seconds (after effect; **e**).

**Extended Data Fig. 13.**
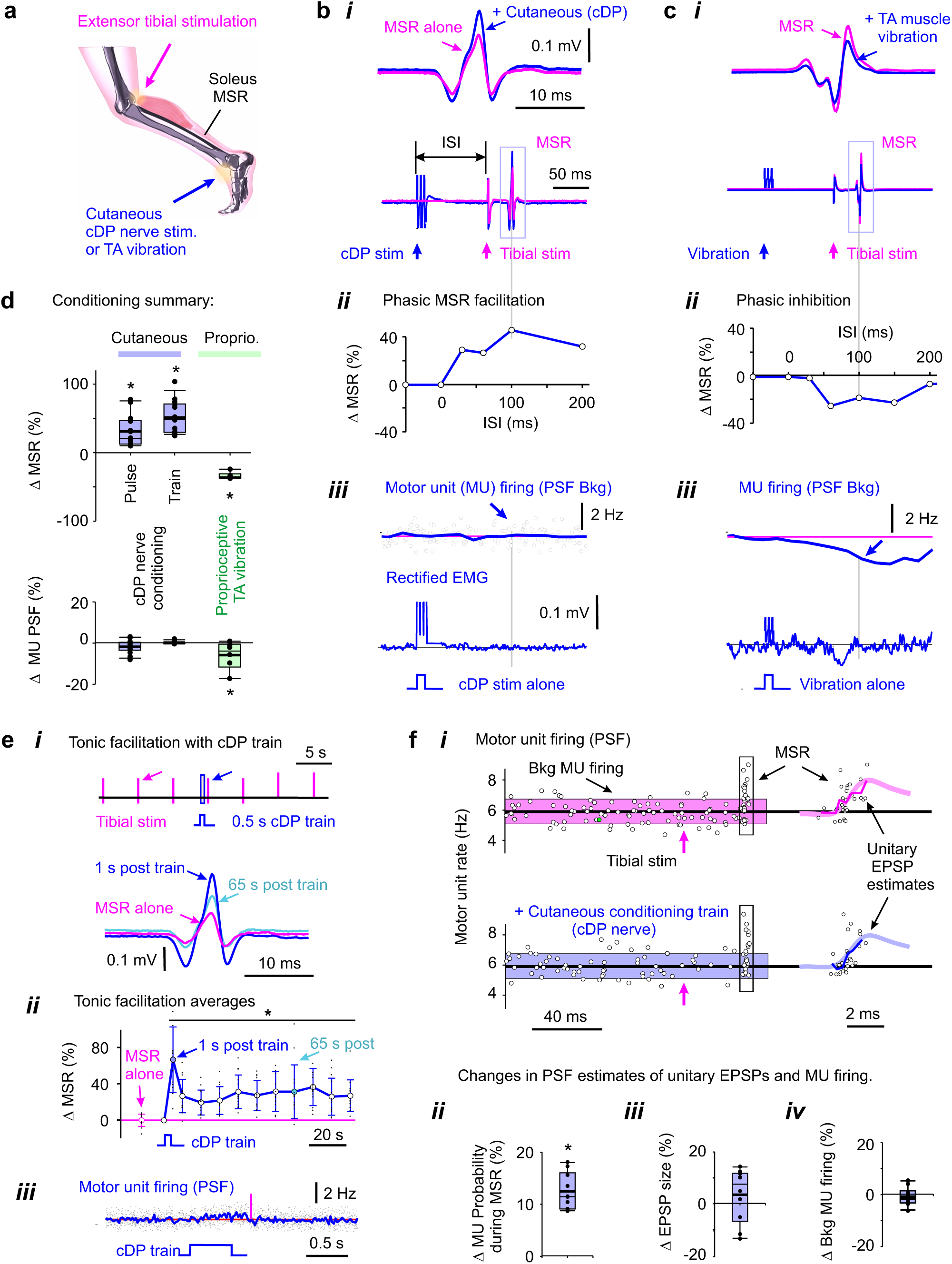
Facilitation of reflexes in humans. **a,** To estimate the role of GABA_axo_ neurons in humans we employed the sensory-evoked depolarization of proprioceptive axons by GABA_axo_ neurons (sensory-evoked PAD; Fig. 4), which is known to occur in humans^86^. For this we recorded the MSR in the soleus muscle in response to tibial nerve stimulation. **b,** MSR in soleus EMG evoked by a tibial nerve pulse (1.1xT, 0.2 Hz, **b***i*), and phasic facilitation of the MSR following a brief conditioning of the cutaneous branch of the deep peroneal nerve (cDP nerve) at varying intervals (ISIs, **b***ii*, 1.0xT, perception threshold T, at rest), and lack of changes in background (Bkg) motor unit (MU) activity or EMG evoked by conditioning alone (**b***iii*, peri-stimulus frequencygram, PSF; with weak contraction). **c**, Same as (**b**), but with proprioceptive conditioning evoked by a brief tibial anterior (TA) muscle tendon vibration, which alone inhibited MU activity (postsynaptic inhibition, PSF Bkg, **c***iii*). **d**, Summary box plots of changes in MSR and postsynaptic (MU) activity with brief conditioning (cDP, *n* = 14 subjects; or TA vibration, n = 6 subjects; as in **b-c**) and long cutaneous conditioning trains (**e**, *n* = 14 subjects). * significant change with conditioning, *P <* 0.05. **e**, Tonic increase in MSR (tonic facilitation) after 0.5 s cutaneous conditioning train (cDP, 1.1xT, 200 Hz) at rest (**e***i-ii*), without prolonged changes in MU activity induced by conditioning alone (**e***iii*, PSF in weak contraction). MSR evoked by tibial stimulation every 5 s, with averages from repeated conditioning shown in (**e***ii*). * significant change in MSR, P < 0.05, n = 5 subjects. **f,** Overlay of all MU firing rates (PSF) with repeated MSR testing (at 5 s intervals) during ongoing weak contraction, and effect of the 0.5 s cutaneous conditioning train (**f***i*). Summary box plots of increased probability of MU firing during MSR (**f***ii*), without changing estimated EPSP size (**f***iii*, PSF thin line; thick line unitary EPSP shape from Fig. 5j) or background MU firing (Bkg, **f***iv*). * significant change with conditioning, *P <* 0.05, *n =* 10 subjects. **Summary of findings in humans**: Increasing GABA_axo_ neuron activity with a brief cutaneous stimulation increased the MSR (**a, b***i***, d**) during a period consistent with nodal facilitation by PAD (30 – 200 ms post stimulation; **b***ii*). We again kept the conditioning stimulation small enough to not change the background EMG or single motor unit (MU) firing (**b*iii***) to rule out postsynaptic actions. When we instead increased PAD by a proprioceptive conditioning (via muscle TA vibration) the soleus MSR was inhibited (for up to 200 ms; **c***i-***c***ii*), as previously reported^90^. However, the vibration alone inhibited the ongoing MU discharge (**c***iii*), implying that this MSR inhibition was caused in part by postsynaptic inhibition, rather than PAD-mediated presynaptic inhibition^90^. Repeated cutaneous conditioning stimulation (trains) to induce a buildup in this tonic PAD caused an associated buildup of the MSR that outlasted the conditioning and its postsynaptic actions by many seconds (after effect; **d,e**). Finally, the probability of a single MU contributing to the MSR was increased by cutaneous conditioning (**f***i-ii*), without increasing the estimated EPSP amplitude or rise time (PSF; see Methods; **f***iii*) or changing in the MU firing prior to the MSR testing (**f***iv*; motoneuron not depolarized closer to threshold), consistent with decreased branch point failure (Fig. 5).

**Extended Data Fig. 14.**
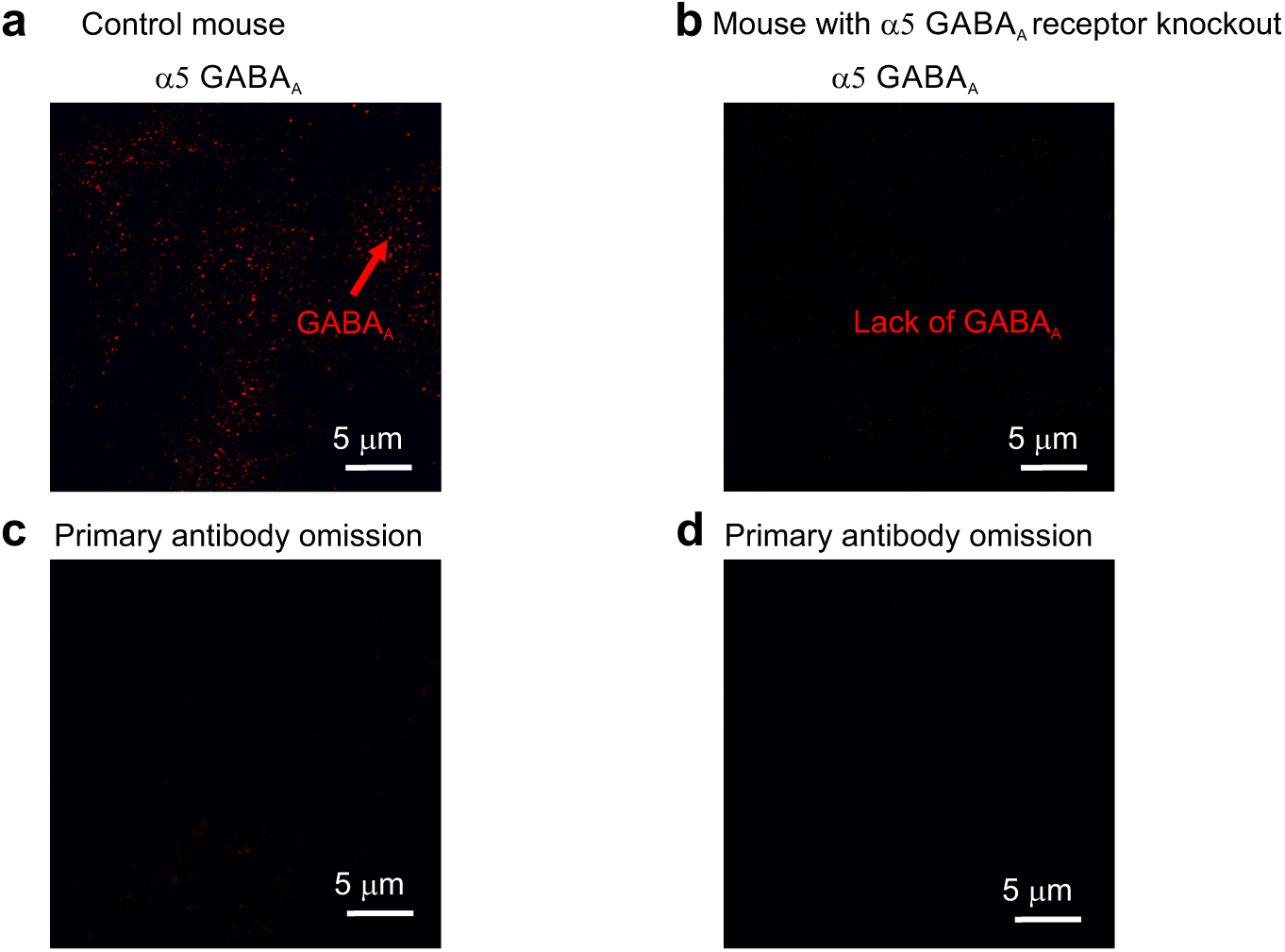
Lack of α5 GABA_A_ receptor immunolabelling after receptor knockout, verifying antibody selectivity. **a**, Immunolabelling of α5 GABA_A_ receptors with antibody to rabbit anti-α5 GABA_A_ receptor subunit (1:200; TA338505, OriGene Tech), as used in Fig 1 and Extended Data Fig 1. Images taken in hippocampal region of wildtype adult mouse brain where neuronal α5 GABA_A_ receptors are highly enriched. **b**, Lack of α5 GABA_A_ receptor immunolabelling in α5 GABA_A_ receptor knockout mouse (Gabra5 KO), from same region. **c-d**, Primary antibody omission controls in wild type and Gabra5 KO mice, respectively, where sections were processed as in (**a**) and (**b**), but no rabbit anti-α5 GABA_A_ receptor antibody applied. Tissue sections in (**a**) and (**b**) were processed for immunolabelled side-by-side on the same slide, and images were obtained with identical confocal microscope settings and displayed at the same brightness as in antibody omission controls of (**c**) and (**d**) where no labelling was observed.

**Extended Data Table 1.** Chronological list of evidence contradicting the classical concept of presynaptic inhibition of transmitter release from proprioceptive sensory axon terminals on motoneurons.

| Date | Contradictions in classic view of terminal presynaptic inhibition mediated by terminal GABA<sub>A</sub> receptors and PAD | Resolution of contradictions |
| --- | --- | --- |
| **1938** | *Primary afferent depolarization (PAD) directly evokes spikes in sensory axons, producing excitation rather than presynaptic inhibition.* Barron and Matthews (1938)<sup>22</sup> discovered that sensory nerve conditioning evokes a long depolarization in many other sensory afferents (primary afferent depolarization, PAD), which we now know is mostly GABA<sub>A</sub> mediated<sup>129</sup>. They and others noted that sometimes this PAD was large enough to directly induce axon spiking, even in vivo<sup>47</sup>, including spikes in the sensory axons mediating the MSR itself, raising a contradiction with the notion of GABA mediated presynaptic inhibition<sup>15</sup>. While these PAD-triggered spikes do not fully propagate antidromically out the DRs in many axons (they fail en route), they are actually initiated in most axons and more likely to conduct orthodromically<sup>15</sup>, making most axons and their motoneuron synapse refractory to subsequent testing (*post activation depression*). Indeed, numerous groups have shown that these spikes directly activate the MSR pathway<sup>6,8,134,135</sup>. Thus, these PAD-evoked spikes must inhibit afferent transmission in the MSR pathway by making axons refractory and producing post activation depression of their terminal synapse, even in humans where PAD evoked spikes occur<sup>86</sup>. This indirect inhibition is GABA<sub>A</sub> mediated and thus readily mistaken for presynaptic inhibition (sensitive to GABA<sub>A</sub> antagonists)<sup>136,137</sup>, even though the PAD-evoked spike is fundamentally excitatory. Even Eccles noted this issue, and showed that just the refractory period alone in the sensory axon inhibits the MSR<sup>8</sup>. | *Post activation depression from PAD evoked spikes inhibits the MSR and masks facilitation of the MSR by nodal facilitation.* We find that facilitation of the MSR by conditioning evoked PAD is always reduced when it is associated with a large enough conditioning stimulation to evoke spikes in sensory afferents, which likely results from post activation depression of axon transmission. This likely explains why Fink et al.<sup>5,16</sup> recently saw inhibition of the MSR with optogenetic or sensory activation of GABA<sub>axo</sub> neurons (see Fig 4.12c in Fink<sup>16</sup> for PAD evoked EPSC inhibiting the MSR). When looking for MSR facilitation, avoiding these spikes and post activation depression requires using weak conditioning stimuli, unlike previous studies<sup>8,17</sup>. |
| **1949** | *Post-tetanic potentiation (PTP) of the MSR increases sensory nerve conduction, but its mechanisms have remained elusive.* Lloyd (1949)<sup>138</sup> concluded that increasing conduction along sensory axons (not just terminals) contributed to the minutes of facilitation of the MSR seen after a high frequency nerve stimulation train (PTP). However, he supposed this might be due to hyperpolarization of the sensory axons, even though we now know that such trains depolarize axons via tonic PAD<sup>15</sup>. The tonic PAD from these bursts must overwhelm the hyperpolarization driven by Na-K pump activity<sup>139</sup>. Lloyd also concluded that PTP only occurred when the same nerve is used for the train (tetanus) as for testing the MSR. | *Repetitive nerve stimulation produces a tonic GABA<sub>A</sub> mediated depolarization (PAD) of axons that facilitates nodal conduction, and increases the MSR.* This PAD likely contributes to PTP, and is largest when the same nerve is tetanized, compared to other nerves, explaining why Lloyd missed the subtler facilitation from other nerves. |
| **1958** | *PAD is associated with a lowering of the threshold for activating spikes.* Early on Wall (1958)<sup>32</sup> noted that a conditioning nerve stimulation that depolarized sensory axons (PAD) was associated with a lower threshold to extracellularly activate these axons. Subsequently this was assumed to be due to the action of terminal GABA<sub>A</sub> receptors and presynaptic inhibition, and spike threshold changes were used to estimate the size of PAD<sup>129,140</sup>. | *PAD lowers the spike threshold via GABA<sub>A</sub> receptors at or near nodes assisting the sodium spike.* This is not related to presynaptic inhibition, but can still be used to estimate PAD as Rudomin and others have done. |
| **1957 - 1994** | *PAD is not correlated with inhibition of the monosynaptic reflex (MSR).* Shortly after Frank and Fortes discovered that the leg extensor muscle MSR is inhibited by a conditioning of a flexor nerve in cats (PBST; like Fig. 6)<sup>141,142</sup>, Eccles proposed the concept of presynaptic inhibition mediated by this conditioning depolarizing of the proprioceptive sensory axon terminals in the MSR pathway (PAD), simply because the MSR inhibition and PAD are somewhat correlated in time<sup>8</sup>. However, in retrospect PAD is far too brief to account for the much longer inhibition caused by this conditioning<sup>44,132</sup>, and some flexor nerve conditioning (a single PBST pulse) inhibits the MSR (Fig. 1 of Eccles, 1961<sup>8</sup>), even though it does not cause PAD in the extensor proprioceptors of the MSR at all<sup>143</sup>. | *PAD is correlated with nodal spike facilitation and facilitation of the MSR.* PAD causes facilitation of the MSR, explaining this correlation. When PAD is large and evokes axonal spikes, these cause post activation depression (detailed above) that should also be correlated with PAD, but is not due to presynaptic inhibition. Also, barbiturates used by Eccles and others potentiated GABA<sub>A</sub> receptor currents. |
| **1959 - 1993** | *Postsynaptic inhibition inevitably accounts for part of the inhibition of the MSR by flexor nerve conditioning.* In his initial short report Frank (1959)<sup>141</sup> correctly suggested that the early inhibition of the MSR by flexor nerve conditioning might be partly postsynaptic (rather than presynaptic), on motoneuron distal dendrites. Others dismissed postsynaptic inhibition because the decay times of the EPSP does not always change when the EPSP is reduced by conditioning, which they proposed indicated that there was no postsynaptic change in conductance in distal dendrites<sup>46,129</sup>. However, this method is likely not very sensitive, due to | *Postsynaptic inhibition masks facilitation of the MSR by nodal facilitation.* We find evidence for long lasting postsynaptic inhibition on distal motoneuron dendrites during nerve conditioning stimulation (including postsynaptic reductions in Tau, Vm and unitary EPSP heights and single |
|  | variability in unitary EPSP time course. Also, anatomically ~70% of GABA<sub>axo</sub> contacts on afferent terminals also contact motoneurons (in a triad), so postsynaptic inhibition is likely inevitable<sup>7,45</sup>. | MU firing). Crucially, minimizing postsynaptic inhibition requires using a small conditioning stimulation when looking for MSR facilitation, unlike previous studies<sup>17</sup>. |
| **1961 - 2014** | *Self-facilitation during repeated MSR testing reduces the possibility of observing facilitation with subsequent conditioning stimuli, leaving only inhibitory actions of conditioning.* Eccles and others knew that the same proprioceptive nerve stimulation that activates the MSR also depolarizes these proprioceptive afferents (PAD self-activation)<sup>8</sup>. Thus, just the act of repeatedly testing the MSR to find the average MSR prior to conditioning pre-activates PAD and produces self-facilitation of the MSR, reducing the headroom to observe changes in the MSR following a separate nerve conditioning stimulation that produces PAD. However, at the time it was not known that repeated nerve stimulation causes a tonic buildup of GABA and a tonic PAD that alters sensory transmission and MSR even at long repetition intervals of many seconds. Thus, Eccles and others used short test intervals (1 s) and strong maximal MSR test stimuli<sup>6,8,17</sup>, presumably assuming that there would be no interaction between test stimuli, which is not the case. In retrospect, these short test intervals and strong test stimuli must have preactivated tonic GABA, leaving little headroom to observe facilitation of the MSR (facilitation), and leaving mainly only inhibitory action possible. | *Self-facilitation masks facilitation of the MSR by a conditioning stimulation.* To observe facilitation of the MSR by a conditioning stimuli that produces a PAD it is important to use long test intervals (5 - 10 s) and small MSR test intensities (1.1xT) to minimize self activation of a tonic PAD prior to conditioning. While experimentally troublesome, self facilitation during repetitive activation is actually one of the main functions of PAD, allowing sensory axons to faithfully transmit spikes to motoneurons at high frequencies that would otherwise produce sodium spike inactivation. |
| **1980** | *Sensory axon terminal potentials at motoneurons are consistent with spike failure.* Early efforts to examine how spikes propagated to sensory axon terminals employed extracellular recordings (EC) near the motoneurons, called terminal potentials (Sybert et al. 1980)<sup>133</sup>. However, unlike EC recordings from near conducting axons (Fig. 2b), these terminal potentials lacked much of the obvious negative field associated with the action potential, and instead had a prominent positive field, followed by a smaller negative field (Extended Data Fig. 10 and Sybert<sup>133</sup>). This positive field has been shown in other axons to be indicative of spike propagation failure and result from the passive axonal current caused by the last non-failing node, similar to a FP, as demonstrated in motor axon recording<sup>74,76</sup>. Indeed, we found that even dorsal horn recordings could exhibit this positive field if the nearby dorsal root conduction is blocked with a microinjection of TTX (Extended Data Fig. 10d). Sybert<sup>133</sup> went on to show that with PAD evoked by nerve conditioning this positive terminal potential field was decreased, and incorrectly interpreted this as evidence for decreased spike conduction and thus supposed it was due to presynaptic inhibition. | *Positive terminal potential fields are decreased with PAD, indicative of decreased conduction failure, consistent with Sybert<sup>133</sup>.* There is a small negative field that follows the positive field in terminal potential recordings, representing spikes that actually reach the terminals. We quantified negative field and found it to increase with PAD, consistent again with increased spikes conducting to motoneurons (Extended Data Fig. 10). Blocking activity in the spinal cord with glutamate antagonists, which would include blocking GABA<sub>axo</sub> circuit activity, decreased this negative field. |
| **1988 - 1998** | *GABA<sub>B</sub> receptors cause presynaptic inhibition and related RDD.* Decades, after Eccles popularized the notion of GABA<sub>A</sub> mediated presynaptic inhibition, Curtis (1998)<sup>44,136</sup> concluded that the late part of the inhibition of the MSR by flexor nerve conditioning is instead GABA<sub>B</sub> receptor mediated, since it is reduced by the GABA<sub>B</sub> antagonist CGP55845 (as Fink also showed<sup>16</sup>), and as is RDD<sup>144</sup>. RDD is a rate dependent depression in the MSR during repeated testing. We suggest that RDD is partly mediated by a build up of GABA released by GABAergic neurons onto the terminals during this repeated MSR testing, though activity dependent homosynaptic depression likely also contributes<sup>145</sup>. | *GABA<sub>B</sub> mediated presynaptic inhibition masks facilitation of the MSR by GABA<sub>A</sub> receptors.* GABA<sub>B</sub> receptors are located on the terminals, and produce presynaptic inhibition (Fig. 5) and RDD (Bennett and Hari, unpublished results), which are reduced by GABA<sub>B</sub> antagonists (CGP55845) or silencing GABA<sub>axo</sub> neurons. |
| **1990 - 1998** | *GABA<sub>A</sub> receptors have direct postsynaptic inhibitory effects on many spinal neurons, making the actions of GABA<sub>A</sub> antagonists difficult to attribute to presynaptic inhibition.* By the 1990s Redman and others tried to confirm the role of GABA<sub>A</sub> receptors in presynaptic inhibition by locally applying the GABA<sub>A</sub> antagonists bicuculline to the spinal cord, and indeed found this drug or other antagonists reduced the inhibition of the MSR by flexor nerve conditioning<sup>17,44,136,146</sup>. However, we now know that this is indirectly due to bicuculline causing a widespread disinhibition of the spinal cord (including loss of post activation depression, detailed above) that leads to a convulsive spinal cord with very long lasting polysynaptic reflexes evoked by the nerve conditioning or the MSR testing itself, making pre and postsynaptic actions hard to distinguish. Further, we know that GABA<sub>A</sub> receptors mediate dorsal root reflexes and associated post activation depression of the MSR (see above point), and thus bicuculline and picrotoxin likely reduce the inhibition of the MSR via reducing post activation depression (see above), rather than changing presynaptic inhibition. | *GABA<sub>A</sub> receptor antagonists reduce the MSR, by reducing nodal facilitation.* Postsynaptic GABA<sub>A</sub> receptors have potent inhibitory actions on many spinal neurons involved in polysynaptic reflexes. However, minimizing these polysynaptic reflexes (by using weak test stimuli and blocking NMDA receptors, Fig 5c) reveals a direct inhibition of the MSR by GABA<sub>A</sub> antagonists, as does optogenetically silencing GABA<sub>axo</sub> neuron, consistent with GABA<sub>A</sub> receptors facilitating rather than inhibiting sensory transmission. |
| 1990 - 1995 | *PAD recorded in the dorsal roots cannot arise from terminal GABA receptors, due to spatial attenuation on the axon.* With advent of detailed anatomical and computer models of sensory axons<sup>25,147,148</sup> it became clear that signals like spikes or PAD are attenuated over short distances in axons, < 200 $\mu\text{m}$. This implies that PAD recorded on or near the DR is unlikely to bear any relation to terminal presynaptic inhibition, despite claims to the contrary<sup>6,90,129,146</sup>. | *Space constant $\lambda_s$ of sensory axons is about 90 $\mu\text{m}$.* Thus, the PAD recorded in the dorsal root must arise from GABA receptors at or near nodes, and not bear any relation to GABA action at the terminals 1000 $\mu\text{m}$ away. |
| 1994 - 1999 | *Shunting inhibition produced by axon terminal GABA<sub>A</sub> receptors is not adequate to produce presynaptic inhibition of the MSR.* Numerous invertebrate studies proposed that the conductance from GABA<sub>A</sub> receptors in terminals caused a reduction in spike height via its shunting action that contributed to presynaptic inhibition with nerve conditioning<sup>23</sup>. However, the effects of conditioning on spikes was small and terminals were not actually recorded from. Subsequently modelling considerations led to the conclusion that shunting inhibition is not adequate to produce presynaptic inhibition and calcium was somehow involved<sup>147</sup>, possibly further implicating GABA<sub>B</sub> receptors, as we see. Considering our estimated space constant $\lambda_s$ of ~90 $\mu\text{m}$, the small shunting inhibition of the spike height (1 mV) we observe is very unlikely to prevent the spike produced at a given node from activating a downstream neighbouring node, since nodes are ~50 $\mu\text{m}$ apart, leading to only about a 50% reduction in spike height at the downstream node (to ~40 mV, unless of course the node is failing), which is well above that needed to initiate a full nodal spike. Thus, spike propagation is very unlikely to be blocked by shunting inhibition. Also, terminal boutons are mostly on unmyelinated axons without sodium channels (passive, 3rd order), and so a 1% reduction in the spike arising from the last/closest node on the 2nd order branch will have little effect on the terminal depolarization (1%), ruling out substantial shunting inhibition of transmitter release from the terminal. | *GABA<sub>A</sub> receptors only slightly decrease spikes by shunting conductances, and otherwise assist nodal spike conduction in proprioceptive axons.* In non-failing secure spikes in sensory axons GABA<sub>A</sub> receptors lower the threshold for spike activation (rheobase) and speed the spikes, the latter by decreasing the time constant of the axon (RC). They do decrease the spike, but only by about 1%, consistent with shunting being unlikely to inhibit spike transmission to motoneurons. However, this does not rule out densely expressed GABA<sub>A</sub> receptors causing shunting and presynaptic inhibition in cutaneous afferents, as previously suggested<sup>15,73,149</sup>. |
| 1994 - 1998 | *Sodium spike inactivation from axon terminal GABA<sub>A</sub> receptor depolarization is not adequate to produce presynaptic inhibition.* Early poor quality recordings from sensory axons (resting near -50 mV from penetration injury)<sup>150</sup> led to the prevailing view that spike failure with depolarization (PAD) was much more common than we now find with better recordings (resting near -70 mV, Extended Data Fig. 3b). Further, Redman later questioned this view, and it seems unlikely for the MSR pathway<sup>17,137</sup>. | *Physiological PAD depolarizations do not block proprioceptive sensory axons spikes, and instead prevent them from failing in the MSR pathway.* However, this does not rule out GABA<sub>A</sub> receptors causing spike inactivation in other axons<sup>15</sup>. |
| 1995 - 1998 | *Physiological GABA<sub>A</sub> receptor activation is unlikely to produce branch point failure in the sensory axons of the MSR pathway.* Over the years sensory axon conduction failure has been occasionally noted from indirect observations<sup>29,31,43,73,131,151-153</sup>. Wall and others<sup>25,73,149</sup> questioned whether this failure could be increased by GABA. However, Wall thought GABA should inhibit rather than assist spikes by inactivating sodium channels. However, Wall was misled by two issues. First, at the time low quality recordings from sensory axons may have led to the misconception that spike failure with physiological depolarizations (like PAD) was common<sup>150</sup>, unlike what we observe. To be fair, Wall was studying cutaneous, as well as proprioceptive, afferents, which are more densely innervated by GABA receptors<sup>15</sup>, making spike inactivation by PAD more likely<sup>73</sup>. Second, by this time GABA<sub>A</sub> and associated PAD had been firmly entrenched as synonymous with presynaptic inhibition. | *GABA<sub>A</sub> receptors help prevent branch point failure and thus facilitate sensory transmission in the MSR.* Computer simulations by Walmsley and others<sup>25,147</sup> have led to the conclusion that physiological GABA<sub>A</sub> receptor conductances cannot stop spikes from propagating past a node. Instead, we report here that they help prevent spike failure near branch points, including in our computer simulations. |
| 1996 - 2018 | *Lack of GABA<sub>A</sub> receptors on proprioceptive Ia axon terminals.* Extrasynaptic $\alpha 5$ GABA<sub>A</sub> receptors are lacking at most proprioceptive axon terminals in the ventral horn<sup>15</sup>. Synaptic GABA<sub>A</sub> receptors also appear to be lacking from these terminals, though only indirectly studied<sup>5,6,24</sup>. GABA<sub>B</sub> receptor immunolabelling had not been investigated in these Ia afferents, though it has in others (Aβ)<sup>154</sup>. | *GABA<sub>A</sub> receptors are mostly at nodes, whereas GABA<sub>B</sub> receptors are at terminals in large proprioceptive afferents.* |
| 2005 - 2014 | *GABAergic innervation of axons.* Recently, GAD2 expressing GABAergic neurons were identified that make axoaxonic connections onto presynaptic terminals of proprioceptive axons<sup>5-7</sup> (termed GABA<sub>axo</sub> here). Previously, Walmsley found GABAergic P-boutons contacting nodes of these axons. Subsequently, Kolta and Zytnicki again found GABAergic contacts near branch points of mammalian afferents<sup>149,155</sup>, as did Cattaert in crayfish<sup>23</sup>. | *A key role of GABA<sub>axo</sub> neurons is to innervate proprioceptive afferent nodes via GABA<sub>A</sub> receptors and ventral terminals via GABA<sub>B</sub> receptors, producing nodal facilitation and presynaptic inhibition, respectively.* |
| 2018 | *GABA<sub>axo</sub> neuron activation by sensory conditioning does not depolarize proprioceptive axon terminals.* Direct recordings from the fine terminals of proprioceptive afferents reveal that during sensory conditioning the terminal is not depolarized during the long PAD recorded on dorsal roots<sup>15</sup>. | *GABA<sub>axo</sub> neuron activation depolarizes nodes.* Dorsally located nodes produce the PAD recorded in dorsal roots. |

**Extended Data Table 2.**
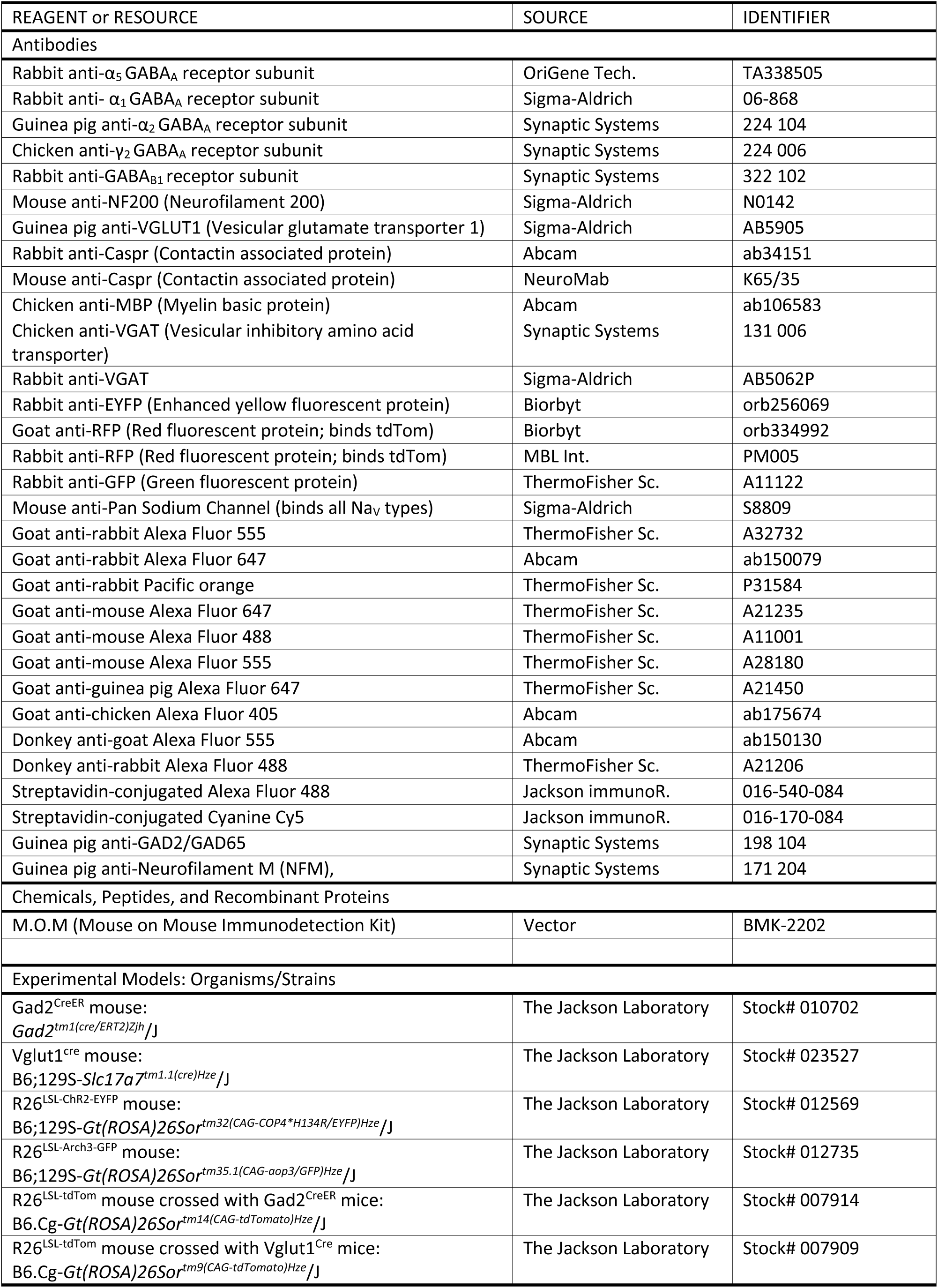
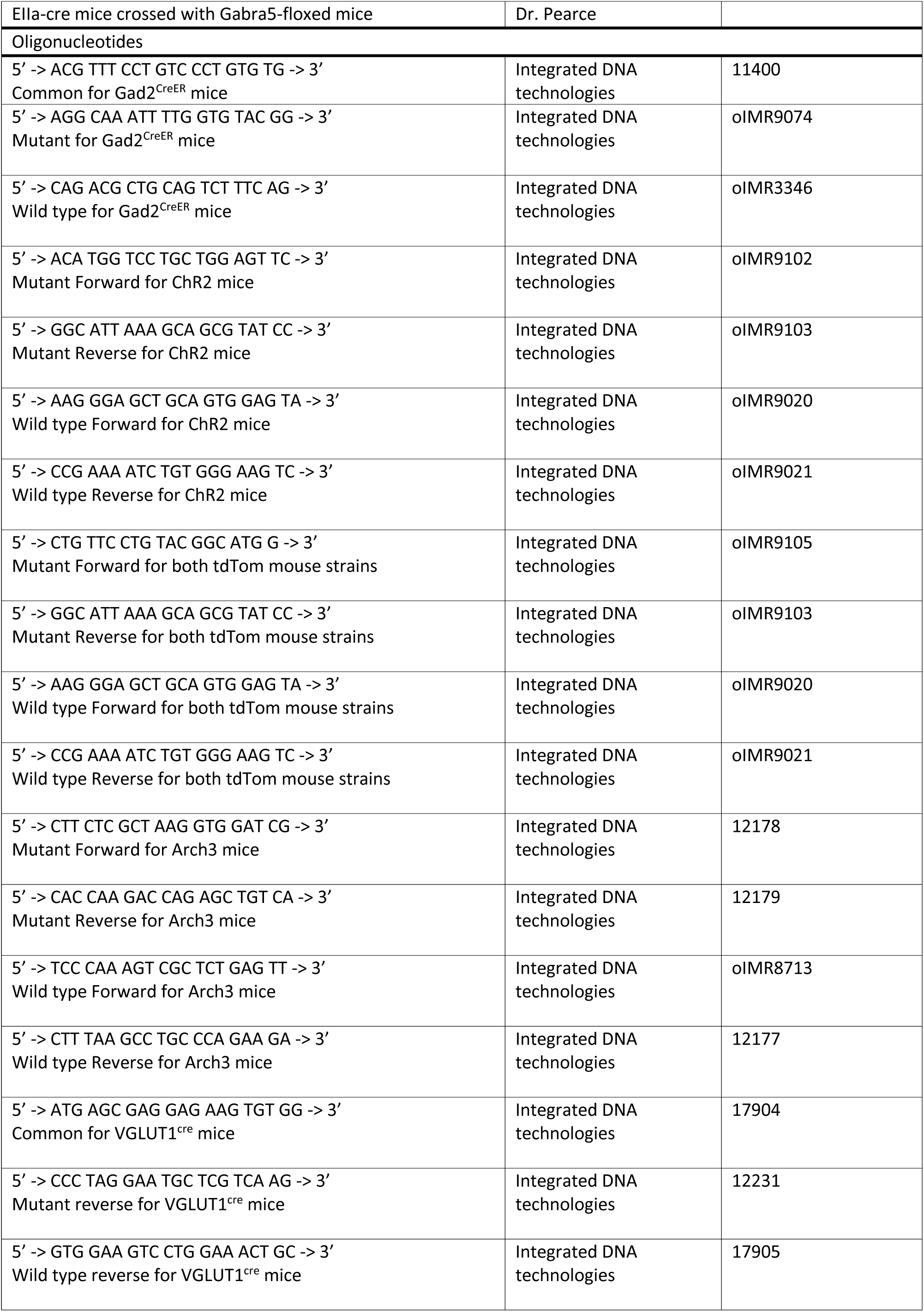
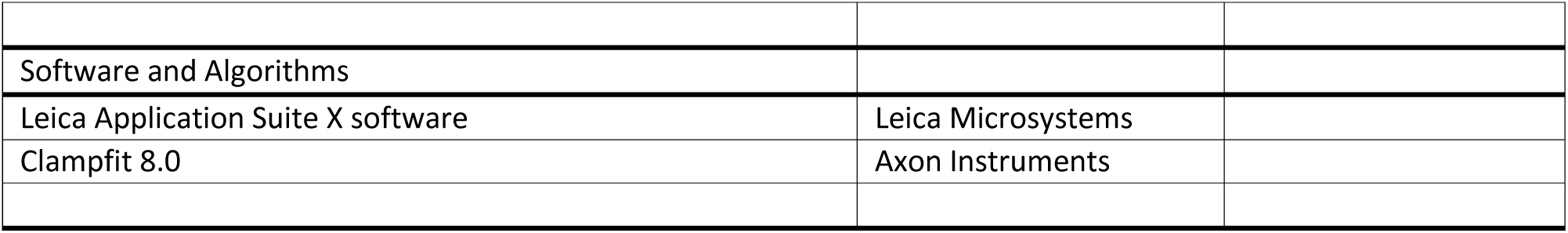
Resources used in Methods.

